# Analysis of cellular and synaptic mechanisms behind spontaneous cortical activity *in vitro*: Insights from optimization of spiking neuronal network models

**DOI:** 10.1101/2021.10.28.466340

**Authors:** Jugoslava Aćimović, Tuomo Mäki-Marttunen, Heidi Teppola, Marja-Leena Linne

## Abstract

Spontaneous network bursts, the intervals of intense network-wide activity interleaved with longer periods of sparse activity, are a hallmark phenomenon observed in cortical networks at postnatal developmental stages. Generation, propagation and termination of network bursts depend on a combination of synaptic, cellular and network mechanisms; however, the interplay between these mechanisms is not fully understood. We study this interplay *in silico,* using a new data-driven framework for generating spiking neuronal networks fitted to the microelectrode array recordings. We recorded the network bursting activity from rat postnatal cortical networks under several pharmacological conditions. In each condition, the function of specific excitatory and inhibitory synaptic receptors was reduced in order to examine their impact on global network dynamics. The obtained data was used to develop two complementary model fitting protocols for automatic model generation. These protocols allowed us to disentangle systematically the modeled cellular and synaptic mechanisms that affect the observed network bursts. We confirmed that the change in excitatory and inhibitory synaptic transmission *in silico*, consistent with pharmacological conditions, can account for the changes in network bursts relative to the control data. Reproducing the exact recorded network bursts statistics required adapting both the synaptic transmission and the cellular excitability separately for each pharmacological condition. Our results bring new understanding of the complex interplay between cellular, synaptic and network mechanisms supporting the burst dynamics. While here we focused on analysis of *in vitro* data, our approach can be applied *ex vivo* and *in vivo* given that the appropriate experimental data is available.

**New & Noteworthy:** We studied the role of synaptic mechanisms in shaping the neural population activity by proposing a new method to combine experimental data and data-driven computational modeling based on spiking neuronal networks. We analyze a dataset recorded from postnatal rat cortical cultures *in vitro* under the pharmacological influence of excitatory and inhibitory synaptic receptor antagonists. Our computational model identifies neurobiological mechanisms necessary to reproduce the changes in population activity seen across pharmacological conditions.

## Introduction

The emergence of spontaneous activity in form of network bursts is a hallmark of functional maturation of mammalian sensory cortices particularly at postnatal developmental stages [1–5]. These bursts have been observed in cortical networks, including the developing cortex *in vivo*, see e.g. [6–11], cortical slices *in vitro* [12–17], and dissociated cortical cultures *in vitro* [18–24]. This type of network activity is thought to prime cortical circuits for incoming sensory information [11]. The activity develops in parallel with the structural and synaptic connectivity [25] and abnormalities in the process lead to neurodevelopmental disorders [15,25,26]. The interaction of excitation and inhibition through synaptic transmission is essential for the proper functioning of these neural circuits and impaired synaptic functions have been associated with a number of neuropsychiatric disorders and neurological diseases [26–30]. Earlier studies have examined the impact of glutamatergic AMPA and NMDA receptors [31–34] or GABA_A_ receptors [23,32,35,36]. However, the effects and interaction of GABA_A_R on predominantly either AMPA or NMDA receptor-mediated network activity have not been studied in cortical cultures prior to [24,37] (but see a study in hippocampal cultures in [38]). In order to explain fully the functions and mechanisms of network bursting it is essential to combine efficiently experimental protocols and data with the data-driven computational modeling.

We propose a new data-driven computational modeling formalism that combines spiking neuronal network models with a rich set of experimental data described in detail in our previous study [24]. In [24], we quantify the changes in network burst structure induced by pharmacological blocking of individual or combinations of synaptic receptors (AMPAR, NMDAR, GABA_A_R). In this new computational study, we disentangle the synaptic, cellular and population level mechanisms contributing to these experimentally measured changes in burst structure. We focus on two measures to characterize the burst structure, burst length and burst size. Burst length (BL) is the time between the onset and offset of a network burst. Burst size (BS) is the total number of spikes, recorded from all active electrodes, normalized to the number of active electrodes. The proposed computational approach permits to study the role of individual and combinations of biophysical mechanisms in a well-controlled setup and to establish their causal impact on network activity. As a result, we identify the experimental findings that can be explained solely by reduction in synaptic transmission and we also propose the minimal set of biophysical mechanisms necessary to reproduce all experimental data.

To achieve these goals we developed two model fitting protocols named the ‘flexible protocol’ and the ‘constrained protocol’. In each of them, we fitted a spiking neuronal network model to a subset of the experimental data, and then assessed how well the fitted model reproduces the remaining experimental data. The model consists of spiking neurons [39,40] randomly connected via synaptic connections that explicitly incorporate the pharmacologically manipulated receptor types (see [24]), AMPAR, NMDAR and GABA_A_R [41–45]. This model, although reduced to a small subset of biophysical mechanisms, is flexible and can exhibit various dynamical regimes across the parameter space (see e.g. [46]; or analysis of networks of integrate-and-fire neurons in e.g. [47–49]). We fitted the model parameters using intensive simulations, genetic algorithm and multi-objective optimization [50,51], and then applied additional criteria to select the best parameter set among the fitting results. The model equipped with this parameter set is called ‘the best fitted model’ in what follows. In the context of computational neuroscience, this methodology is typically used to construct multi-compartmental single neuron models [50,52].

We developed the ‘flexible protocol’ to test whether we can achieve a quantitatively accurate reproduction of all experimental data, extracted from the network burst activity in each experimental protocol. We allowed maximal flexibility by fitting one spiking network to each pharmacological condition. This way we compensate for the possible cellular, synaptic and changes in extracellular conditions induced by receptor blocking in our experimental preparation. Prior to model fitting we performed *in silico* blocking of synaptic receptors consistent with the considered pharmacological condition. Using this protocol, we reproduced the experimentally obtained burst lengths and sizes in four out of five conditions, and the correct burst lengths in the fifth condition.

The ‘constrained protocol’ allowed exploring which experimental findings can be attributed solely to the reduced synaptic transmission induced by pharmacological blocking. We fitted one network model to the data obtained from all pharmacological conditions. *In silico* blocking was still performed according to [24] but the cellular and synaptic mechanisms were not allowed to adapt to each individual condition. In this case, we were able to reproduce the relative changes in burst lengths and sizes across conditions as seen in our experimental study. More precisely, our fitted model was capable of reproducing the increase or decrease in these burst measures across pharmacological conditions, but not the exact experimentally recorded values. Our results obtained with multiple model fittings suggest that burst measures could be better reproduced when single-neuron bursting/adaptation is allowed in the cell model. On the other hand, allowing the heterogeneity across neuronal and synaptic types did not significantly improve model fitting.

In conclusion, we developed new model fitting protocols to explore the biophysical mechanisms behind network bursting in cortical networks *in vitro.* Our results reveal the impact of individual biophysical mechanisms and describe how the interactions between these mechanisms give rise to the properties seen in the experimental data. In what follows, we first describe our methodology, including a brief overview of our experimental setup, data analysis and details of the computational methodology. Next, we summarize the experimental data and findings that we reproduce and analyze using computational methodology. Result sections 2 and 3 present the outcomes of the ‘flexible protocol’, and result section 4 and 5 presents the outcomes of the ‘constrained protocol’. For each presented result, we analyze the fitted model parameters and establish the robustness of our results. We finish with discussion of our findings.

## Methods

### Experimental data

All the experimental data used to construct and fit computational models in this study were recorded, analyzed, and presented in our earlier publications [24,37]. Here, we re-analyzed this previously published data into the format suitable for automatic model fitting. A part of the data is already made publicly available through our previous publication [24]. The data needed to reproduce the results of this paper are shared as supplementary information.

#### Experimental procedures

The preparation and maintenance of cell cultures, pharmacology, and microelectrode array recordings were described in [24,37]. Briefly, primary cells were derived from the prefrontal cortices of postnatal (P0) Wistar rats of either sex. Tissue was mechanically chopped and enzymatically trypsinized and dissociated. Cortical cells including neurons and astrocytes were re-suspended into minimum essential medium, seeded at densities of 2,000 cells/mm2 onto MEAs (MultiChannel Systems Ltd., Reutlingen Germany) and cultured until 20-25 DIV when all recordings were done. A total of 13 cell cultures from three different preparations were used in the study: 6 cultures were used for pharmacological experiments of NMDR blockade (10μM, D-(-)-2-Amino-5-phosphonopentanoic acid, D-AP5) followed by GABA_A_R blockade (10 μM, Picrotoxin, PTX), and 7 in experiments of AMPAR blockade (10 μM, 2.3-Dioxo-6-nitro-1,2,3,4 tetrahydrobenzo-[f]quinoxaline-7-sulfonamide, NBQX) followed by GABA_A_R blockade (10 μM PTX). Blocking the function of these receptor types allowed characterization of their contribution to network activity during electrophysiological recordings. All drugs were directly applied with a pipette into the culture medium inside the dry incubator. Each cell culture was used to conduct a three-hour experiment (Fig 1A) inside a dry incubator: **1)** one hour recording of the control (CTRL) condition, **2)** application of NMDAR or AMPAR antagonist to culture medium, **3)** another one hour recording, **4)** application of GABA_A_R antagonist to disinhibit the culture, and **5)** the third hour of recording. The list of applied reagents and their impacts are given in Fig 1B (all pharmacological antagonists were purchased from Sigma-Aldrich, Steinheim, Germany).

**Fig 1:**
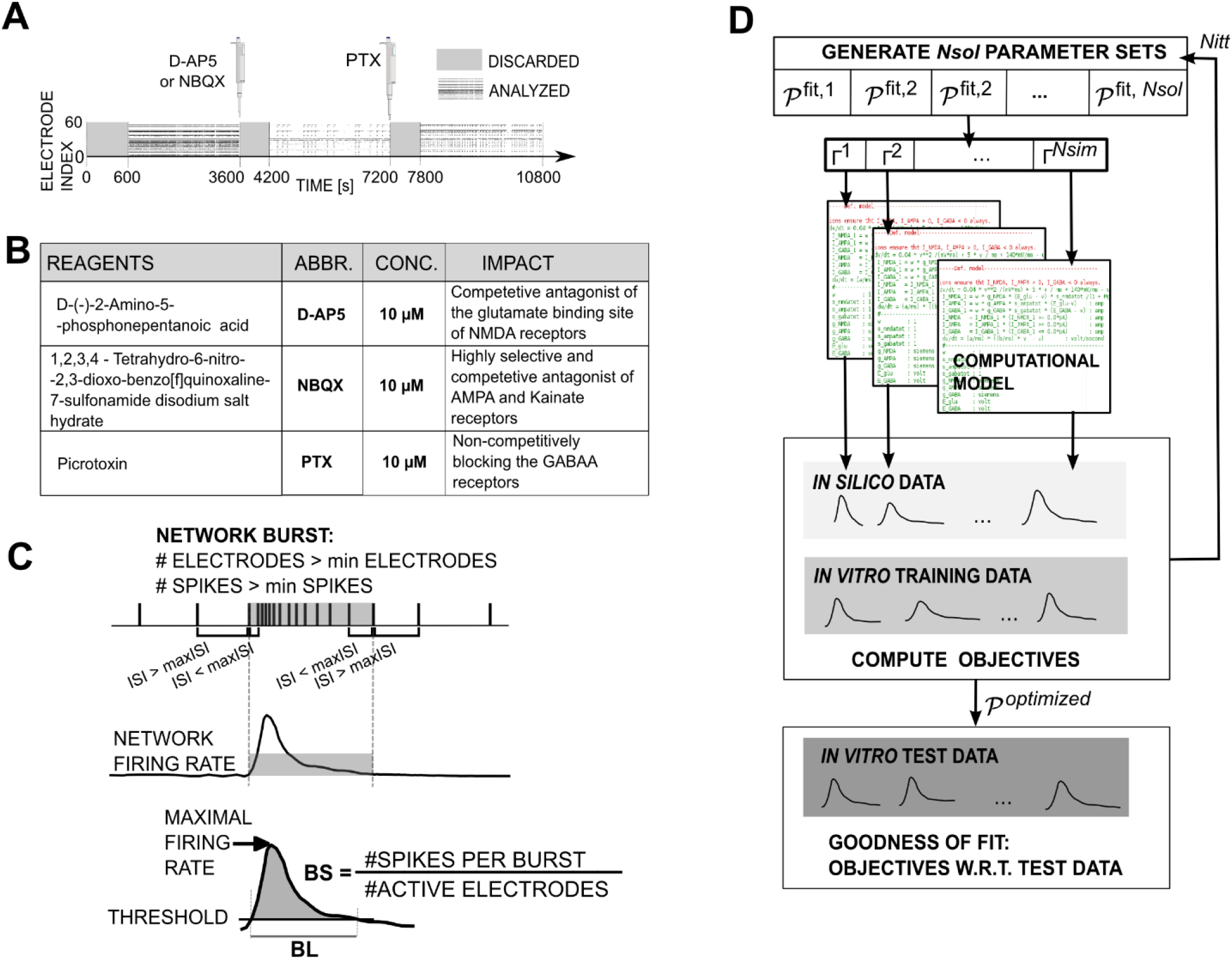
Summary of experimental and computational methods. **A:** Illustration of the experimental protocols used to collect the data. The network bursts *in vitro* were recorded using the standard MEA chip containing 60 electrodes (y-axis). First, the control recordings were collected for one hour. Next, either AMPARs or NMDARs were pharmacologically blocked and the activity was recorded for the next hour. Finally, the culture was disinhibited by applying GABA_A_Rs blocker and the final hour of recording was done under this pharmacological condition. For each one-hour recording, the first 10 minutes were removed (shaded intervals) and only the latter, stabilized activity was used for model fitting. **B:** The table lists reagents used for pharmacological blocking of glutamatergic and GABAergic receptors. **C:** Summary of the data analysis used to process spike trains and extract burst measures. Top - burst detection based on spike timings in the population spike trains (combined spikes from all electrodes). Middle - burst profile. Bottom - extracting burst size (BS; the number of spikes in a burst normalized to the number of electrodes activated during that burst) and burst length (BL; time between the onset and offset of a burst) from the burst profile. **D:** Model fitting protocol. In each iteration (*n* = 1.. *N*_*itt*_) of the model fitting algorithm a set of *N*_*sol*_ parameter sets is generated by genetic algorithm. For each of those parameter sets 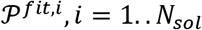, the *N*_*sim*_ models are generated by drawing a random connectivity scheme with the fixed probability of connection between pairs of neurons. Simulating these models results in *N*_*sim*_ network bursts and corresponding BL and BS estimations. That *in silico* data is compared to the training data set (70% of the available *in vitro* data) to evaluate the objective functions. After completing all iterations, the final model is selected and its success is evaluated against the test data set (30% of the available *in vitro* data).

### Microelectrode array recording and data preprocessing

A detailed description of the recording methodology is presented in our earlier publications [24, 33]. In short, the extracellular network activity was recorded from cortical cultures grown on microelectrode arrays (MEAs; Multi-Channel Systems Ltd, Reutlingen, Germany, MCS). The collected recordings were preprocessed and the extracellular spikes were detected as described in [24]. The first ten minutes of the recordings were discarded to remove the effect of initial perturbation caused by drug application (see Fig 1A). The remaining 50 minutes were analyzed using MATLAB toolboxes MEA-tools [97] and FIND [98]. The spike times detected at individual electrodes were combined into population spike trains and further analyzed using own MATLAB scripts [24].

### Burst measures extracted from experimental data

The 50 minute recordings contained 4-722 network bursts in different cultures and pharmacological conditions. We extracted individual bursts using a standard detection method (Fig 1C). The interspike-intervals (ISI) between the successive spikes in a recording were computed: beginning of a burst corresponds to the first pair with ISI ≤ 100ms, ending of a burst corresponds to the last pair satisfying this condition. We kept only the sufficiently large bursts, those that spread over at least 10 electrodes. Burst detection accuracy was validated through manual inspection of the obtained bursts. In total, we extracted 4913 bursts from the control condition of 13 cultures, between 155 and 722 bursts per culture. The experiments with NMDAR blocking resulted in 1368 network bursts, between 34 and 704 bursts per culture, and the experiments with additional disinhibition gave 1342 bursts, 75-305 per culture. The blocking of AMPARs dramatically reduced the number of bursts. However, we still recorded 120 network bursts from 7 cultures, 4-34 per culture, which provided enough data for the model fitting described in what follows. Disinhibition of these cultures increased the number of bursts to 500, 20-132 per culture.

Next, we computed the burst profiles by binning the population spike rates and then convolving them with a Gaussian kernel [24]. We described the burst structure using two measures (Fig 1C): **1)** the network burst size (BS), i.e. the number of spikes within a burst normalized to the number of active electrodes, **2)** the network burst length (BL), the time when the burst profile exceeds a threshold value (set to 1/16 of the burst profile maximum, adapted from [53]). The adopted definition provides a robust measure of BLs and reduces the trial-to-trial variability particularly affecting the burst tails. Next, we evaluated the distribution, means, medians, and percentiles (25th and 75th) for these two measures across cultures and pharmacological conditions. The results obtained from different pharmacological conditions were compared using statistical tests (U tests, p=0.05). In order to use this data for model fitting, we pooled all results (all bursts from all cultures) for the same pharmacological condition. In Result sections 2 and 3, 70% of the data was used for training, i.e. for fitting the computational model while the remaining 30% was used for testing, i.e. to evaluate the goodness of fitting.

#### Computational Model

We constructed a spiking neuronal network composed of N=100 point neurons, *N*_*E*_=80 excitatory and *N*_*I*_=20 inhibitory. In what follows, we outline the model in equations (1–7) and emphasize dependency on model parameters. A detailed description of the model in the standardized tabular format according to [54] is given in the Appendix 1.

We adopted a standard neuron model [39,40] described by equations (1–2) and the spike-reset condition:

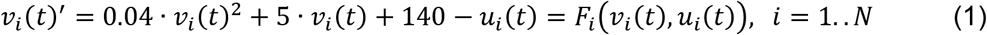

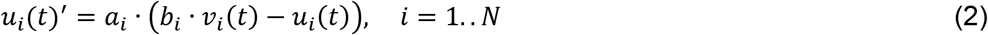

The spike-reset condition is given as follows: whenever 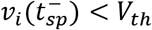 and 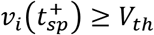 a spike is generated at time *t*_*sp*_ and the membrane-potential and adaptation variables are reset as 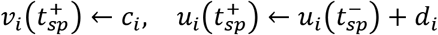.

### This neuron model depends on the following parameters

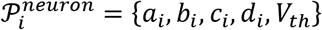. Embedded into a network model, the expression for the cell membrane potential becomes:

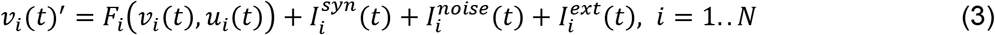

The cell membrane dynamics is driven by the three currents: 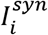- the sum of all synaptic currents, 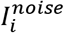- the noisy current that accounts for nonspecific sources of membrane fluctuations, 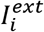- the external stimulus (when exists). 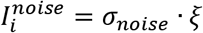 is a random variable depending on the variance 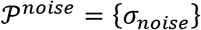 and a normally distributed random number 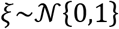. We used a standardized simulator (see below) which ensures accurate numerical integration of the random variable *ξ*. For all neurons, the noise was set to a small value that can affect the subthreshold dynamics but cannot induce spiking on its own (tests with zero noise were also done). The external stimulus in the form of a pulse current was injected into a few neurons (1 or 10) to simulate a small perturbation that initiates bursting. This stimulus is defined by duration and amplitude of the pulse 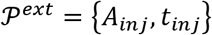.

Synaptic current incorporates the inputs from the considered glutamatergic and GABAergic receptors:

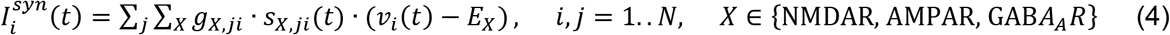

Here, *g*_*X*,*ji*_ is the maximal value of the synaptic conductance for the receptor type X (NMDAR, AMPAR or GABA_A_R) on a synapse from neuron j to neuron i. The variable *s*_*X*,*ji*_ defines the collective dynamics of synaptic receptors of type X on the incoming synapse from neuron j to neuron i. The AMPAR and GABA_A_R conductances are increased instantaneously at each incoming spike and decay exponentially with the time constants *τ_AMPA_* and 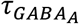 between spikes. The NMDAR is described according to a two-dimensional phenomenological model from [41,42]:

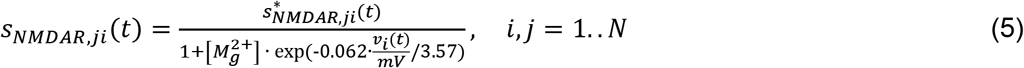

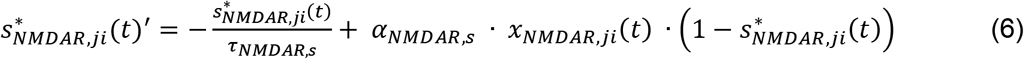

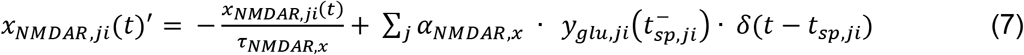

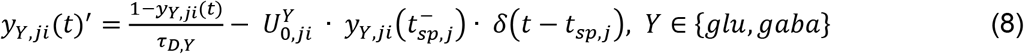

All three receptor models are updated at each presynaptic spike *t*_*sp*,*ji*_ for an amount equal to 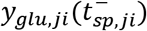 for glutamatergic and 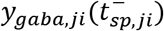 for GABAergic synapses. This amount accounts for the presynaptic depletion of synaptic resources which might be substantial during a prolonged burst and can be scaled by a constant (*α*_*AMPAR*_ and 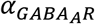 for AMPA and GABA_A_ receptors, and *α*_*NMDAR,s*_, *α*_*NMDAR,x*_ for NMDA receptors). The model has been used before e.g. in [43], the equation (8) is equivalent to the short-term depression in [55]. The variables *y*_*glu*,*ji*_(*t*) and *y*_*gaba,ji*_(*t*) are instantaneously decreased at each presynaptic spike by constant amounts 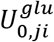 and 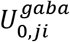 respectively. Between spikes they exponentially increase towards their maximal values with the time constants *τ*_*D*,*gl,u*_ and *τ*_*D*,*gaba*_. The s and y variables take values between 0 and 1.

### The synaptic model is determined by the following parameters

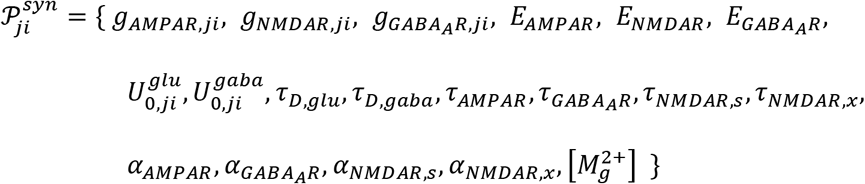

Note that the index ‘ji’ means that each individual synapse receives a different value for this parameter. The index is absent for parameters that are set to be the same across all synapses.

The presence or absence of a synapse between a pair of neurons is defined by the binary connectivity matrix 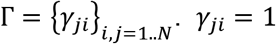 if there is a synapse from neuron *j* to neuron *i*, *γ*_*ji*_ = 0 otherwise. Thus, the variable *S*_*X*,*ji*_ exists and contributes to synaptic currents only if *γ*_*ji*_ = 1 and X is the appropriate receptor type. In this study, Γ is a random connectivity matrix where each neuron randomly connects to 20% other neurons in the network.

#### Model implementation

The model was implemented using Brian 2.0 simulator (https://brian2.readthedocs.io/en/stable/, [56,57]) and the NEURON simulator version 7.5-7.7 (https://neuron.yale.edu/neuron/). We used Python 2.7 with Numpy version 1.14.4, and Scipy version 0.14.0 (except for computing U-test where Scipy 0.14.1 was used). The code used for preparing this publication is available in the ModelDB data repository (access will be provided to the reviewers, the repository will be made open access once the manuscript is accepted for publishing). Summary of the content of this repository is given in supplementary material S6. The intensive model fitting was done using the Linux cluster at Tampere University (https://wiki.eduuni.fi/display/tutsgn/TUT+Narvi+Cluster) and CSC high-performance cluster Puhti (project 2003397). The illustrations presented in the paper are produced using the same simulators running on a personal computer (Lenovo G50-70, Intel(R) Core(TM) i7-4510U CPU @ 2.00GHz).

### Model fitting to the experimental data

The described spiking network model depends on neuronal 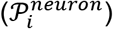, synaptic 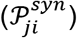, noise 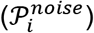, and external current 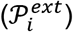, parameters. In addition, the connectivity matrix Γ determines the pathways of synaptic interactions. Some of the parameters were equal for all neurons or synapses, while others were allowed to vary across neurons or synapses. In this case we generated individual parameters from a (truncated) Gaussian distribution, with mean and variance of that distribution being treated (and possibly fitted) as model parameters. A subset of model parameters was fitted to the experimental data 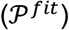, while other parameters were fixed prior to model fitting. We fitted the parameters that control cellular excitability and dynamical regime (regular spiking vs. single neuron bursting), the parameters that control the release of presynaptic resources, and the parameters of postsynaptic currents: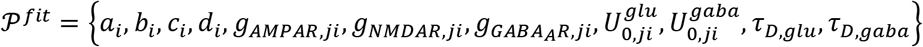.

In some of the tests and for the parameters that differ across neurons or synapses we fitted both the mean and the variance of the parameter distribution. The list of all parameters, the fixed parameter values, and the examples of fitted values are documented in detail in Appendix 1.

The model parameters were selected using iterative model fitting that employs a multi-objective optimization algorithm from the literature [50,52]. The algorithm iterates the following steps (see Fig 1D) *N*_*itt*_ times: **1)** Generate *N*_*sol*_ sets of model parameters, 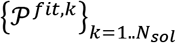. In the first iteration, all the parameters are chosen randomly within the specified ranges of values. In all successive iterations, the parameters are determined according to the employed genetic algorithm, i.e. half of the solutions are inherited from the previous iteration and other half generated by applying crossover and mutation operations to those inherited solutions. **2)** For each 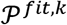 generate *N_sim_* random connectivity matrices 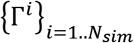 and simulate the models to obtain *N*_*sim*_ bursts for each parameter set. The previously described small perturbation, i.e. injecting a short pulse into a few neurons, triggers a single network burst most of the time. The cases that failed to produce a clear network burst (e.g. because they contained one or several persistently active neurons) were penalized. Generating a new connectivity matrix for each simulation averages out the details in connectivity schemes. **3)** The obtained bursts were analyzed to extract the measures of network burst structure (lengths, BLs, and sizes, BSs) as described before. Selecting a sufficiently big *N*_*sim*_ allowed estimating statistics of these burst measures. **4)** Compute the objective functions used to assess the goodness of fitting. Based on objective functions reject the worst half of the parameter sets and pass the other half to the next iteration. **5)** Once the *N*_*itt*_ iterations are completed, select the best parameters and evaluate the objective functions against the test part of the experimental data set. The parameter with the best performance on the test set are shown in the results section.

For the **multi-objective optimization algorithm,** we used a publicly available Python package downloaded from https://projects.g-node.org/emoo/ and described in [50]. The package adapts the non-dominated sorting genetic algorithm II (NSG-II) for model fitting in neuroscience. The details of the algorithm, including the descriptions of crossover, mutation and criteria for pareto-optimal solutions, are given in the literature [58]. The NSGA-II has been used in various optimization tasks, and in the context of neuroscience for fitting ion-channel densities in multi-compartmental neuron models [50,52] or protein concentrations in biochemically detailed synapse models [59]. The crossover and mutation parameters and the probability of mutation per parameter were fixed to *η_c_* = 20, *η_m_* = 20, *p_m_* = 0.5 (for precise meaning of these parameters see [50,58]. The population size was *N*_*sol*_ = 200 (flexible protocol) or *N*_*sol*_ = 1000 (constrained protocol). The number of iterations *N*_*itt*_ = 20 − 40 was selected to ensure the convergence of the algorithm. To validate our results of the constrained protocol, we used an alternative multi-objective optimization algorithm, NSGA-III [60], as implemented in the Platypus package (https://platypus.readthedocs.io/). In NSGA-III, the population size cannot be explicitly set but is determined by a division parameter (parameter p in [60]) and the number of objective functions. In our validation experiments, we set the number of divisions p=6, which resulted in population size *N*_*sol*_ = 210.

The objective functions used for model fitting were designed to minimize the distance of BLs and BSs between *in vitro* and *in silico* data while preserving the relative differences seen across pharmacological conditions in [24]. Exact formulation of the objective functions for all considered tests is given in Appendix 2. We formulated them by carrying out a series of trials and keeping those that produced the best fits to the data. A distance between two burst measures was computed either as an absolute difference between their means (constrained protocol) or as Jensen-Shannon divergence (JSD) between their distributions (flexible protocol), see detailed described in S2. The JSD takes values on the interval zero to one, with zero corresponding to the perfectly aligning distributions and one to the maximal divergence between two distributions. In order to simplify computation of JSDs, we estimated all distributions using the same support and bins for histograms. Moderate changes of bins did not significantly alter JSD computation.

## Results

### Section 1: Glutamatergic and GABAergic receptors modulate length and size of network bursts

In this section, we review the experimental findings from [24] used as the motivation and a starting point for this computational study. In Fig 2, we summarize the statistics of burst lengths and sizes across pharmacological conditions and cultures and assess the significance of the differences seen across the conditions. Fig 2A-D show the distribution of BLs and BSs across five pharmacological conditions. The leftmost panels correspond to the control data, the middle panels to AMPAR/NMDAR blocking experiments, and the rightmost panels to the data collected after disinhibition by GABA_A_R blocking. The bars in Fig 2E-H correspond to the average BLs and BSs also shown as vertical lines in Fig2A-D. Data from individual cultures are compared using the 1-sided U-test and significance was assessed at p = 0.01 and p = 0.05 levels. The angular brackets are drawn in Fig 2E-H if the null hypothesis was rejected for majority of cultures. One star (*) indicates that it was rejected for one considered significance level (0.01< p <0.05) and two stars (**) indicate that it was rejected for both significance levels (p < 0.01). The number of cultures that confirmed this statistical result is indicated next to the stars. All p and n values are listed in Table 1.

**Table 1:**
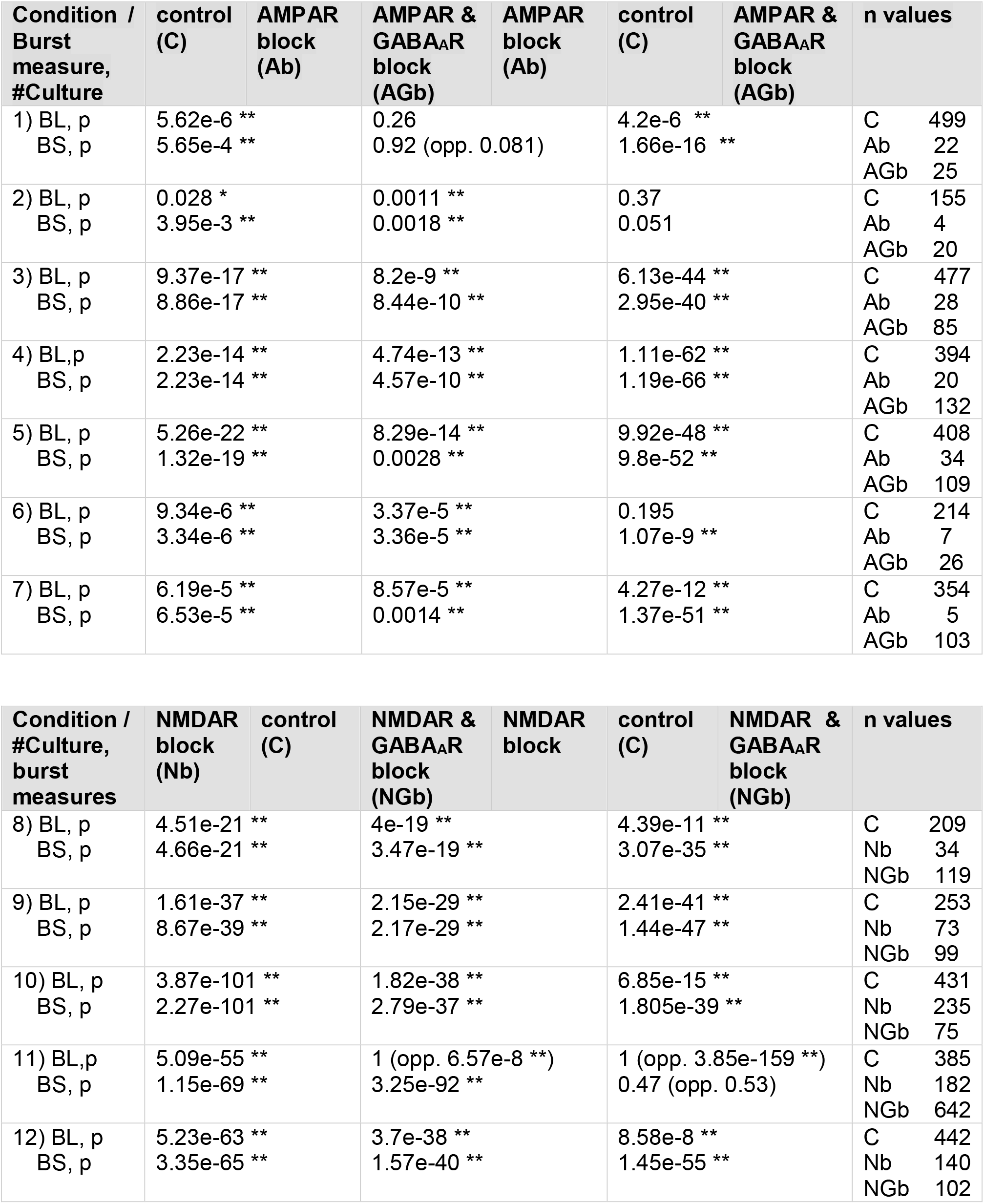
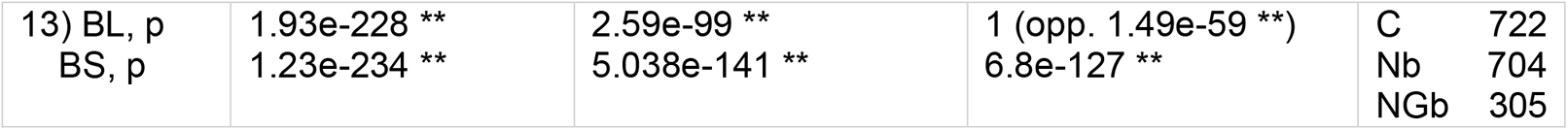
Summary of statistical tests done to compare BL and BS across pharmacological conditions, these results are also illustrated in Fig 2E-H. The rightmost column gives the number of bursts in each culture and condition, i.e. the sample size (n) used in statistical tests. Abbreviations: C – control, Ab – AMPAR block, AGb – AMPAR and GABA_A_R block, Nb – NMDAR block, NGb – NMDAR and GABA_A_R block. In each column, the first condition has **significantly lower** BL and BS than the second condition in majority of considered cultures. Each column gives the p-value (1-sided U test) for BL comparison and for BS comparison. **\*\*** indicates significant result for both considered significance levels, i.e. p<0.01. **\*** indicates significant result for one significance level, i.e. 0.01 < p < 0.05. In some cultures, we found the opposite trend from the dominant one. In those cases, we give the p-value obtained when testing the opposite hypothesis using 1-sided U test; significant results are marked by stars as before.

**Fig 2:**
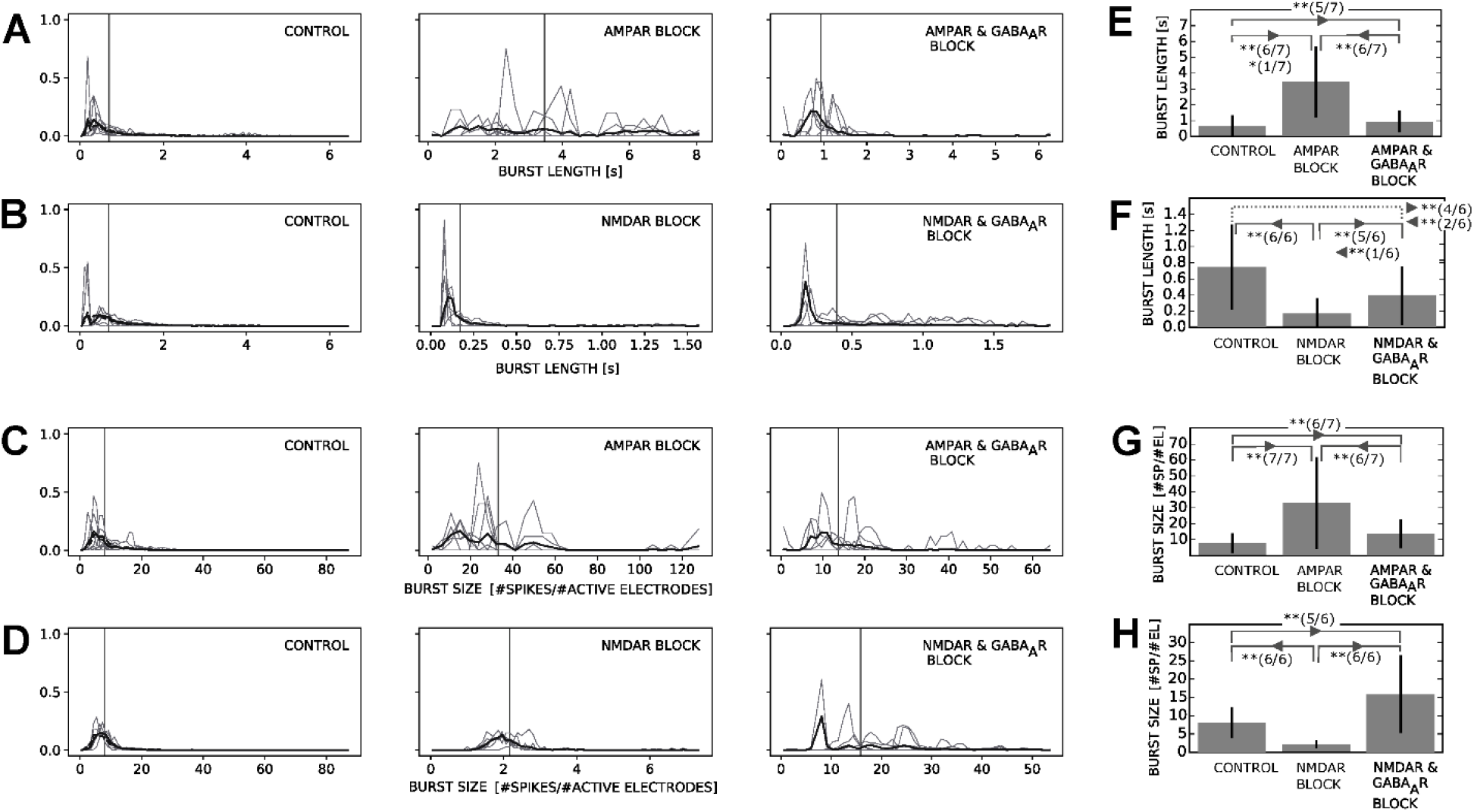
Statistics of experimentally measured burst lengths (BL) and sizes (BS): **A-D: Distribution of BLs and BSs for each pharmacological condition.** Individual cultures are represented by thinner grey lines, while the tick black lines correspond to data pooled from all the available cultures (7 cultures for AMPAR blocking, 6 for NMDAR blocking). In panels illustrating the control data, the interrupted line corresponds to all 13 cultures pooled together. Vertical bars are mean values obtained from the pooled data. **A:** Distribution of BL for control, AMPAR blocked and AMPAR and GABA_A_R blocked condition. **B:** Distribution of BLs for control, NMDAR blocked, and NMDAR and GABA_A_R blocked condition. **C:** Distribution of BSs for the same cultures and pharmacological conditions described in A. **D:** Distribution of BSs for the same cultures and pharmacological conditions described in B. **E-H: Anaysis of the data from panels A-D. Bars represent means and whiskers indicate mean standard deviation. The results of statistical tests (see Table 1) are indicated at each panel.** Lines on the top connect the pharmacological conditions where statistical tests show significantly different BLs or BSs. The arrow points from the condition with lower to the condition with higher BLs or BSs. The number of stars indicates if the significance was established for one or both considered significance levels (p=0.01, p=0.05). The p-values and the sample size for each test are given in Table 1. The number between the brackets indicates how many cultures passed the test out of the total number of considered cultures. **E:** Analysis of the BLs from panel A. **F:** Analysis of the BLs from panel B. The dashed line connecting control with NMDAR & GABA_A_R blocked condition indicates that we could not establish a trend in BL values as 4 (out of 6) cultures pointed at one direction (BL significantly increased when blocking NMDAR and GABA_A_R) and 2 (out of 6) in the opposite direction (BL significantly decreased when blocking NMDAR and GABA_A_R). Note that one culture also showed an opposite trend when comparing BLs in NMDAR blocked and NMDAR & GABA_A_R blocked conditions. In this case, we still concluded that the overall trend can be established as 5 out of 6 cultures show consistent trend (with very low p-values) and the means and variances of the pooled data from all cultures do not contradict this trend (notice the opposite when comparing control and NMDAR & GABA_A_R block conditions). **G:** Analysis of the BSs from the panel C. **H:** Analysis of the BSs shown in panel D.

Blocking of fast AMPARs suppressed the overall capacity to generate network bursts (in total 120 bursts were recorded from 7 cultures, compared to 2471 recorded from the same 7 cultures in control) and significantly increased the length and size of bursts (Fig 2E, 2G). Both, BL and BS were significantly bigger in AMPAR-block condition than in control condition in 7/7 cultures (6/7 for BL if significance is established for p < 0.01; see rows 1-7 in Table 1, column 2 for p-values, column 5 for n-values). Subsequent disinhibition decreased BL and BS (Fig 2E, 2G) in 6/7 cultures and in one culture we could not establish significance (Table 1, rows 1-7, column 3). Still in AMPAR block and disinhibited condition the levels remained significantly higher than in control condition (Fig 2E, 2D). BL in AMPAR and GABA_A_R was significantly higher than in control in 5/7 cultures, and in 2/7 cultures we could not establish significance. BS in AMPAR and GABA_A_R blocked condition was significantly higher than in control in 6/7 cultures, and in 1/7 cultures we could not establish significance (Table 1, rows 1-7, column 4).

The result of blocking NMDARs, which are contributing to prolonged maintenance of network bursting, was significant reduction in BLs and BSs in all 6/6 cultures (Fig 2F, 2H; see Table 1, rows 8-13, column 2 for p-values, column 5 for n-values). Disinhibition significantly increased the size and duration of bursts compared to the NMDAR blocked condition (Fig 2F, 2H; Table 1, cows 8-13, column 3). Disinhibition increased BS in 6/6 cultures, and BL in 5/6 cultures (the opposite trend was found in 1/6 cultures). The BS significantly increased also when compared to the control (Fig 2H) which was confirmed in 5/6 cultures (in 1/6 cultures we could not establish significance; Table 1, rows 8-13, column 3). The difference in BLs between the NMDAR and GABA_A_R blocked conditions and control was not established (Fig 2F). BL was significantly higher in NMDAR and GABA_A_R blocked than in control in 4/6 cultures, and significantly lower in 2/6 cultures (Table 1, rows 8-13, column 4). Taken together, these findings indicate significant receptor-blockade induced differences in network bursts arising from the interplay of the three considered receptor types. To investigate further the mechanisms of this interplay we adopted a computational modeling approach presented in the following sections.

### Section 2: *In silico* models of cell cultures separately fitted to each pharmacological condition reproduce well the experimental findings

The flexible protocol was developed to fit one model for each pharmacological condition. The model is shortly presented in Methods while all details are given in the standardized format [54] in Appendix 1, Tables A-E. We first blocked the synaptic receptors consistently with the modeled pharmacological condition by setting the corresponding peak conductances to zero. The parameters controlling cellular excitability (Izhikevich neuron model), the parameters determining the short-term synaptic depression, and the peak conductances of the remaining active receptors were fitted to the experimental data. The parameters of cellular excitability were randomized across model neurons, and we fitted the mean and variance of their Gaussian distribution (see Methods, all implementation details will be shared in modelDB repository). The parameters of the two neuronal types (excitatory and inhibitory) and the four synaptic types (excitatory-excitatory, excitatory-inhibitory, inhibitory-excitatory, inhibitory-inhibitory) were different and fitted simultaneously. In total, 28 parameters were fitted for AMPAR/NMDAR and GABA_A_R blocked condition, 32 for AMPAR/NMDAR blocked condition, and 36 for control condition.

The goal was to select a model that best reproduces the distributions of BLs and BSs measured *in vitro* (and shown in Fig 1A-D). The model selection was based on two objectives: **1)** the distance between BL distributions measured *in silico* and *in vitro*, **2)** the distance between BS distributions *in silico* and *in vitro* (see Methods and Appendix 2, Table C). The model fitting error was expressed as the distance between distributions computed using the Jensen-Shannon divergence (JSD). The BL and BS distributions were extracted from *N*_*sim*_ = 100 model simulations; each of those models had own randomized parameters for the neuron model and own random connectivity, and all models shared the same (non-randomized) synaptic parameters. Additional source of randomness was a weak Gaussian noise that affected the subthreshold regime but was not strong enough to initiate bursting on its own. A burst was initiated by applying a perturbation (in a form of short current pulse) to one randomly selected neuron (see Methods and model implementation in modelDB/S6). Prior to perturbation, the model was set to the state that correspond to a longer period of inactivity similar to an inter-burst interval: the cell membrane variables were in the steady state, the presynaptic variables were at their maximum and the postsynaptic variables at their minimum. In such a model, population bursts were efficiently introduced at the onset of perturbation, rapidly spread to the entire network, and then decayed to the baseline level.

The model fitting converged to the solution within *N*_*gen*_ = 31 generations for both objectives, as shown in Fig 3 (BL fitting error – Fig 3A, BS fitting error – Fig 3B). In this figure, we show the errors for the 50 best models (according to the model fitting algorithm ranking) and for the three model fitting trials, i.e. 150 models in total. The median errors are shown as orange bars and the boxes spread between 25^th^ to 75^th^ percentiles. As shown in Fig 3, we identified the first generation for which both objectives reach the steady state and those generations are marked by arrows. Note that some error variability remains, although much less than in earlier generations. This is a consequence of randomness in our model, which inevitably causes randomness in the extracted burst measures.

**Fig 3:**
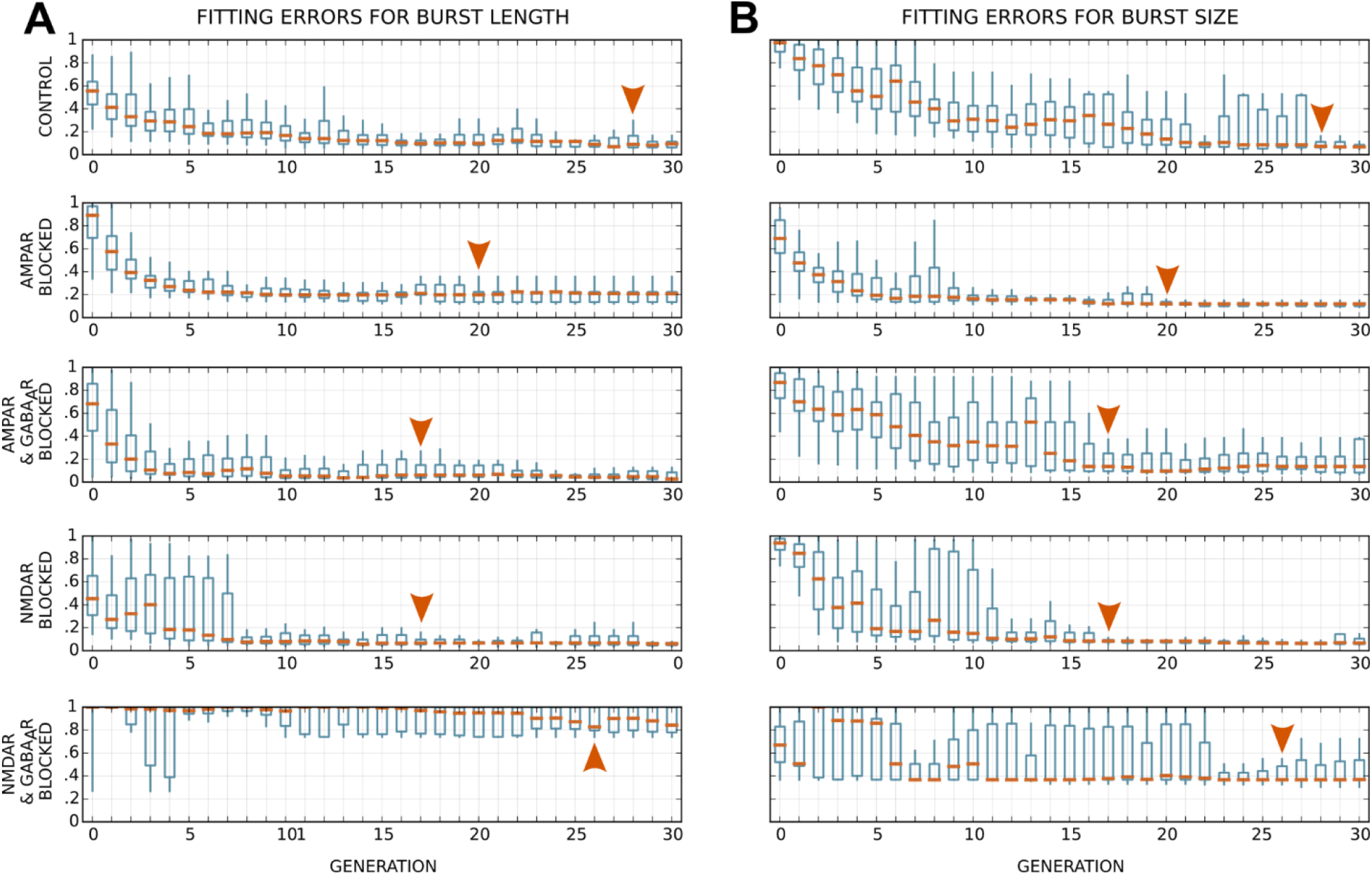
Convergence of the flexible protocol. **A:** Errors of the BL fitting across generations 0-30. **B:** Fitting errors for the BS across generations 0-30. The generations are indicated on the x axes. In all panels, y axes show fitting errors computed as Jensen-Shannon divergence (JSD) between the burst measure distribution extracted from simulation and the one obtained from experimental recordings. JSD=0 – two identical distributions, JSD=1 – maximally diverging distributions, e.g. two distributions with non-overlapping supports. Each row of panels corresponds to one pharmacological condition given as the label of the y axes. The medians (orange bars), 25^th^ and 75^th^ percentiles (boxes) and the extent of error values (whiskers) are shown for 50 best models in 3 model fitting trials (150 models in total) for each generation and each pharmacological condition. The arrows point at the first generation in which the model fitting converges to its steady state.

For the generation chosen above and for each fitting trial, eight candidates for the best model were selected as follows: the two models with the best fitness function according to the model fitting algorithm, two that minimized the BL error, two that minimized the BS error, and two that minimized the sum of BL and BS errors. This way, we obtained 24 candidate models for each pharmacological condition. These models were compared to the test data set, the 30% of experimental data that was not used for model fitting. The majority of these models gave small fitting errors, so we introduced additional selection criteria. For each model, we evaluated whether the obtained BLs and BSs follow the same distributions as the experimental test data using the 2-sided Kolmogorov-Smirnov test (KS) at p=0.05 significance level. We found at least one model passing the test for 4 out of 5 pharmacological conditions. In cases where we found more than one such model or if we found none of them, we selected the model with the minimal sum of BL and BS fitting errors. This procedure provided *the best fitted model* per pharmacological condition. Additional analysis of the BL and BS errors, when compared to the test data set, for all generations and separately for each model fitting trial is shown in the supplementary material S3, Fig S3-1.

The selected model parameters are listed in S1, Table H, and model properties are analyzed in Fig 4. Panels Fig 4A-D show the distribution of BLs and BSs in the same format as the one used in Fig 2A-D for easier comparison. The figure shows good agreement between the experimental (green line) and the simulated data (orange line), with JSD values between 0.046 and 0.2218, in all but one condition. The exception is BS distribution in NMDAR and GABA_A_R blocked condition that largely exceeded the experimentally measured data. In panels Fig 4E-H we compared the *in silico* distributions of BLs and BSs across conditions using the same statistical test as in Fig 2 (1-sided U-test with significance level established for p=0.01 or p=0.05). The sample size was n=100 for all pharmacological conditions and burst measures studied *in silico,* thus it is not separately indicated for each test below. The analysis confirmed that the fitted models reproduce the same trends seen in Fig 2 (indicated by angular brackets at the top). Precisely, BL and BS increased in the AMPAR blocked condition compared to control (p = 4.12e-24 and p = 1.22e-28, respectively), further disinhibition decreased both BL and BS (p = 7.09e-16 and p = 1.51e-21, respectively), but the values remained higher than in control condition (p = 5.98e-9 for BL, p = 5.07e-10 for BS). Blocking of NMDARs led to decrease in BL (p=3.37e-23) and BS (p=1.27e-34) compared to control, while additional disinhibition resulted in the increase in BL (p=1.99e-11) and BS (p=1.27e-34) compared to the NMDAR blocked condition. BS in the NMDAR and GABAAR blocked condition exceeded the values obtained for control condition (p = 1.28e-34), while BL in NMDAR and GABAAR blocked condition was found to be smaller than in control (p = 7.9e-15). This last result differs from our experimental finding due to using pooled data from all cultures in model fitting. When comparing the experimentally obtained BLs in individual cultures the results were inconclusive, 4/6 cultures showed the increase and 2/6 decrease in BL in NMDAR and GABAAR blocked condition compared to control. However when comparing the pooled data from all six cultures we also found significantly smaller BL in NMDAR and GABAAR blocked condition compared to control (p = 2.297e-111, n_NMDAR-GABAAR_=1342, n_CONTROL_ =2442).

**Fig 4:**
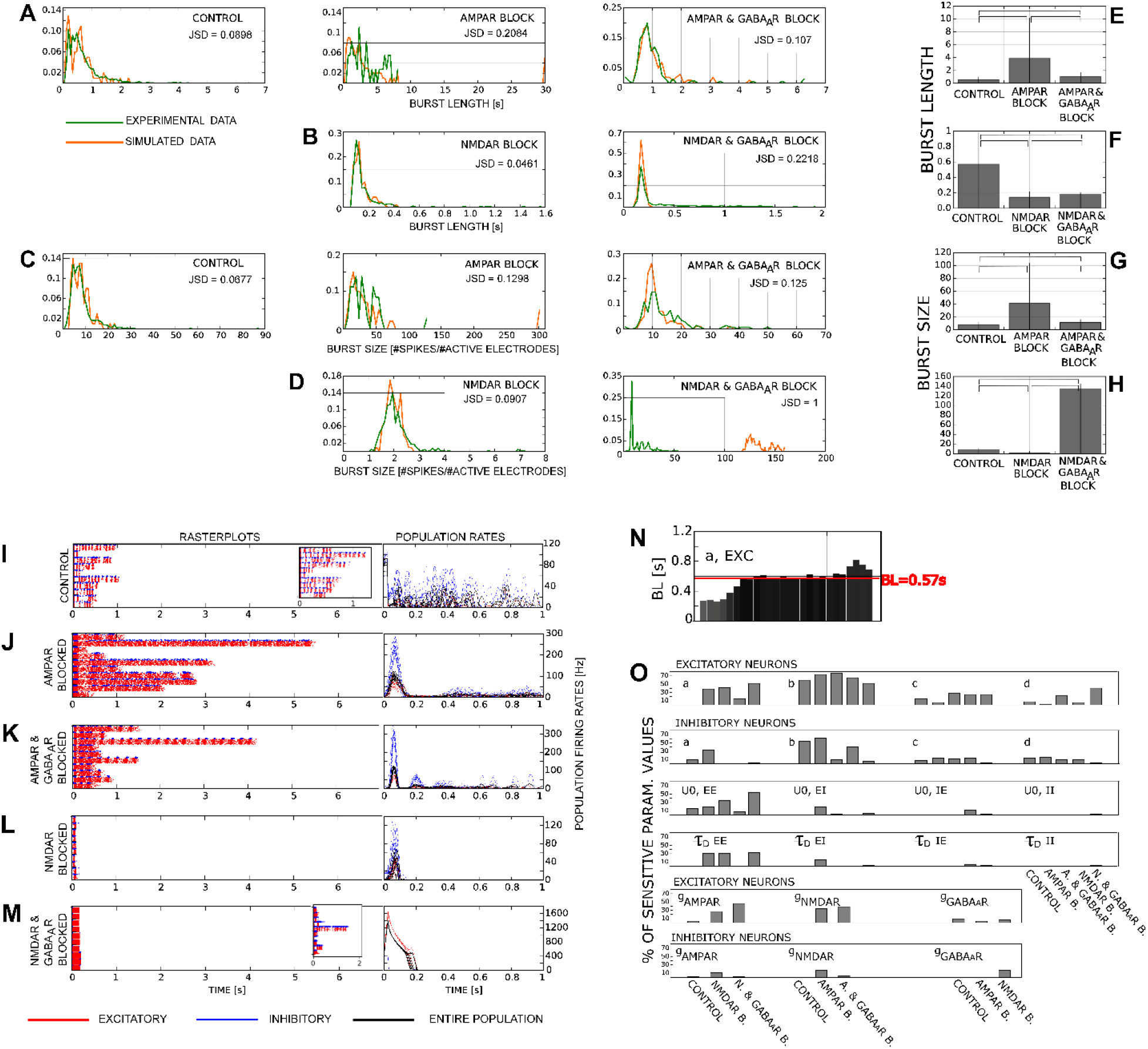
Results obtained using the flexible protocol. **A-D:** Distributions of BLs and BSs for the experimental data and the simulations of the models fitted using the flexible protocol. The green lines show the experimental data (in Fig 2, the data obtained by pooling all available cultures) and the orange ones show the simulated results. The similarity between the experimental and simulated data is quantified using JSD (shown at each panel, JSD=0 for two identical distributions, JSD=1 for two maximally diverging distributions). **E-H:** The mean values (bars) and variances (whiskers) of BLs and BSs across conditions. The simulated BLs and BSs are compared across conditions the same way as it was done for the experimental data. The angular bracket at the top of each panel connect those conditions where the null hypothesis was rejected (1-sided U-test) at significance level p=0.01 and p=0.05. In all considered cases, the significance was established for both levels, thus the mark ** is omitted from the panels. **I-M**: Leftmost panels show the rasterplots of the simulated network bursts for each pharmacological condition. The panels to the right show population spiking rates computed from the corresponding rasterplots. Red - excitatory neurons, blue – inhibitory neurons, black - entire neuronal population. **N:** An example of sensitivity analysis for the parameter ‘a’ in excitatory cells. The bars show the average BL obtained from 100 simulations of the model with varied parameter. The examined parameter values included 1%, 15%, 20%, 25%, 50%, 80%, 90%, 95%, 98%, 99%, 99.5%, 99.8%, 99.9%, 100%, 100.1%, 100.2%, 100.5%, 101%, 102%, 105%, 110%, 120%, 150%, 200%, 300%, 400% of the best fitted value. **O:** The panels illustrate model sensitivity to the variation of each parameter. The sensitivity is expressed as the percent of models where the variation of one parameter led to significant deviation in BL or BS, such that the JSD between the best fitted and altered model exceeded 0.5 (JSD ∈ [0,1]). The JSD is computed over 100 evaluations of the BL or BS. The varied parameter is indicated next to the corresponding results, each bar corresponds to one pharmacological condition (indicated at the x axis).

Panels Fig 4I-M illustrate the burst structure obtained by simulating the best fitted models. The left-hand side of these panels shows 5 rasterplots per pharmacological condition obtained using the best fitted model for that condition. The cellular parameters and the network connectivity were drawn from the random distributions defined by the best fitted model. The panels to the right illustrate the corresponding population rates for the excitatory cells (red), the inhibitory cells (blue) and the entire population (black). We repeated the same simulations while fixing all model parameters and randomizing only the connectivity scheme (Fig 4I – inset panel illustrates the obtained result for the control condition). The variability in burst properties persisted (as seen in Fig 4I) demonstrating the role of randomized connectivity in reproducing the variability in experimental data. However, the coefficient of variation (variance divided by mean of a burst measure computed across 10 simulations of the same model) averaged over all burst measures and pharmacological conditions decreased from 0.044 to 0.0108 when fixing model parameters.

Next, we explored reasons for poor BS fitting in NMDAR and GABA_A_R blocked condition. In our model for this condition, the fast AMPARs are the only mediator of synaptic transmission. We explored whether an incomplete blocking of GABA_A_R could rescue the model and better represent the data. Thus, we repeated the fitting with only NMDAR-blocking being effective, while keeping both AMPAR and GABA_A_R functional. The fitting converged to the steady state within 30 generations and the obtained model provided better fit to the experimental data (see S3, Fig S3-2B). We also repeated the fitting with all synaptic receptors being functional, which resulted in successful model fitting with BL and BS distributions similar to the experimental data (rasterplots shown as an inset in figure Fig4M and S3, Fig S3-C for details of this model fitting). In addition, we went on to fit each pharmacological condition using this full model without synaptic blocking. These fittings quickly converged to the solution in less than 20 generations confirming the flexibility of this model (see S3, Fig S3-3, Fig S3-4 for the fitting convergence and the analysis of fitted models). In conclusion, adding additional flexibility by allowing synaptic transmission through all receptors is sufficient to reproduce the experimental data.

We verified the robustness of our selected models through sensitivity analysis summarized across burst measures and pharmacological conditions in Fig 4N,O. The sensitivity was tested by varying one-by-one model parameter within the wide range from 1% to 400% of the best fitted value. We first compared the burst measures obtained in the model with altered parameter, and in the best fitted model. Fig 4N illustrates one example of such analysis – the average BL (across 100 model simulations) obtained when varying one model parameter in excitatory cells (parameter ‘a’ in Izhkevich model) and in control condition (BL in the best fitted model is marked by red line). The results for other parameters, pharmacological conditions and burst measures are given in S3, Fig S3-5. The analysis showed that a variation of up to ±20% of the original value did not result in deviations from the best fitted model, while larger variations induced larger differences. Next, we computed the JSD between the 100 burst measure evaluations in the altered model and in the best fitted model and marked the cases where JSD exceeds 0.5 (for the JSD computation see S2, Table B). For each parameter and pharmacological condition we counted such cases across all considered parameter variations (other than the best fitted value) and for both burst measures. These counts are shown in Fig 4N as the percent of the total number of examined JSDs. In summary, parameters of excitatory cells induced bigger sensitivity than those of inhibitory cells. For both types of cells, ‘b’ (a parameter in Izhkevich model neuron) did not tolerate large deviations from the best fitted value, except for inhibitory cells in disinhibited conditions that were robust with respect to the larger variations of ‘b’. The parameters of excitatory-excitatory synapses (‘EE’ in Fig 4N) induced more sensitivity than parameters of other synapse types. Expectedly, the sensitivity to variation of glutamatergic peak conductances became higher as we added more pharmacological blockers, i.e. as we removed synaptic mechanisms in the computational model.

### Section 3: Diverse parameter sets provide good reproduction of the experimental data

The results of the previous section show that our model fitting protocol possesses enough flexibility to reproduce the experimental data. We further examined the fitted models and addressed the following questions: **1)** Is the parameter variability beneficial when reproducing any burst measure in any particular pharmacological condition? **2)** Can we identify any consistency between parameter changes and BL and BS changes across pharmacological conditions? In the previous section, we allowed heterogeneity in neuron model parameters but kept the synaptic parameters constant across the synapses of the same type. Here we include two alternatives: **a)** models with no parameter variability, **b)** models with no variability in neuronal parameters but with variability in synaptic parameters, namely in the parameter controlling the depth and time course of presynaptic depression 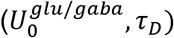 and the three parameters for peak conductances of synaptic receptors 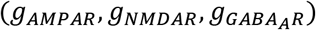. Detailed results obtained for a) and b), comparable to Fig 3 and Fig 4, are presented in the supporting material S3, Fig S3-6 to S3-9.

Fig 5 summarizes these results and evaluates them against the experimental data. The distribution of *in silico* BLs (Fig 5A-E) and BSs (Fig 5F-J) is shown for each pharmacological condition (cyan - no variability in model parameters, orange - variability in neuronal parameters, magenta - variability in synaptic parameters) and the quantitative comparison to *in vitro* data (green) is expressed as JSD (upper right corner). *In vitro* and *in silico* data were also compared using 2-sided Kolmogorov-Smirnov test and stars in figure Fig 5A-J mark the cases where the null hypothesis cannot be rejected. * indicates the cases where significance was established for p>0.01, and ** for p > 0.05. Comparing the histograms in Fig 5A-J showed no major difference between the three model fitting outcomes. In NMDAR and GABA_A_R blocked condition the model with variability in synaptic parameters found better tradeoff although the selected model was still distant from the experimental data. JSD averaged across all conditions and burst measures (individual conditions and burst measures are shown in Fig 5A-J) showed no major difference: variation in neuronal parameters led to the average JSD of 0.2109, in synaptic parameters to the average JSD of 0.1952 and in models with no variability to the average JSD of 0.229. Similarly, statistical tests (see Fig 5A-J, p-values given in figure caption) showed no clear advantage of presence or absence of variability in model parameters. When reproducing BLs the variability in neuron model parameters resulted in the average JSD (across pharmacological conditions) of 0.1408, in models with synaptic parameter variability in the average JSD of 0.2132, and in models with no parameter variability in average JSD of 0.1613. When reproducing BSs the average JSD was 0.2967 when parameter variability was not allowed and 0.2809 and 0.1772 when parameter variability existed in neuronal and synaptic parameters respectively (excluding NMDAR and GABA_A_R blocked condition gives 0.1208 (no variability), 0.1012 (variability in neuronal parameters), and 0.1166 (variability in synaptic parameters)).

**Fig 5:**
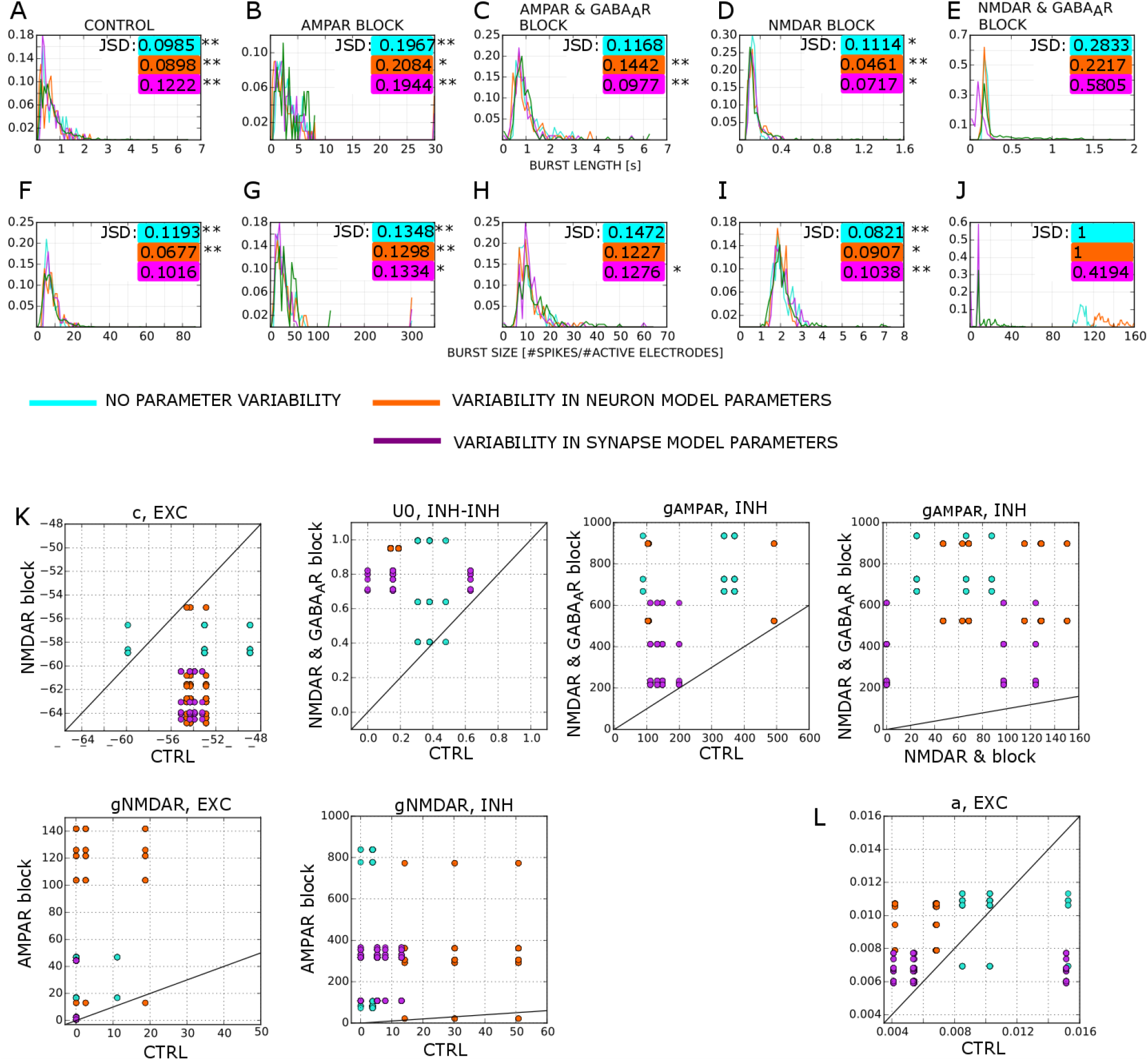
Parameter variability in models fitted by flexible protocol. Model fitting employed the flexible protocol and the following alternatives in parameter handling: **1)** no variability in model parameters allowed (cyan), **2)** the variability allowed only for neuronal parameters (orange), **3)** the variability allowed only for the synaptic parameters (purple). For each pharmacological condition and each of the options 1-3 we repeated model fitting three times and selected the best fitted model as described before. **A-E**: BL distributions for the model fittings 1-3 and for the experimental data (green) for five pharmacological conditions. JSD between the simulated and the experimental data is shown in the upper right corner (color code is the same as for distributions). * and ** indicate the cases where simulated and experimental data belong to statistically indistinguishable distributions for one (0.01<p<0.05) or both (p>0.05) significance levels (2-sided Kolmogorov-Smirnov test; **A:** p = 0.51, 0.089, 0.081 (n_sim_=100, n_exp_=4943); **B:** p = 0.162, 0.025, 0.162 (n_sim_=100, n_exp_= 120); **C:** p = 0.0055, 0.1, 0.481 (n_sim_=100, n_exp_= 500), **D:** p = 0.0226, 0.794, 0.0488 (n_sim_=100, n_exp_= 1368), **E:** p = 1.69e-17, 1.31e-14, 8.98e-50 (n_sim_=100, n_exp_= 1342)). **F-J**: BS distributions. JSD between the simulated and the experimental data is shown in the upper right corner. * and ** mark the same as before. **F:** p = 0.055, 0.307, 0.008 (n_sim_=100, n_exp_= 4943); **G:** p = 0.3, 0.989, 0.025 (n_sim_=100, n_exp_= 120), **H:** p = 0.007, 0.009, 0.031 (n_sim_=100, n_exp_= 500), **I:** p = 0.655, 0.01002, 0.364 (n_sim_=100, n_exp_= 1368), **J:** p = 2.92e-81, 2.92e-81, 3.13e-36 (n_sim_=100, n_exp_= 1342)). **K:** Distribution of the parameters consistent across pharmacological conditions (i.e. in >95% of cases bigger in one condition than in another). Diagonal lines indicate equal parameter in both pharmacological conditions. **L:** An example of a parameter that does not pass the criterion used to select the parameters in panel K.

Next, we analyzed the parameters selected through the model fitting procedure and considered in the test phase. Those included the two sets with maximal fitness function (computed by the multi-objective algorithm), the two sets that minimize either the fitting error of BL or BS, the sum of the two fitting errors. While neuronal excitability clearly changed from one pharmacological condition to another (see Fig 4I-M, and support material S3, Fig S3-7C, Fig S3-9C, Fig S3-12) these changes were achievable through diverse choices of parameter sets, resulting in a wide parameter ranges in Fig 5. Diversity in neuronal parameters was also evident when recording the cell membrane potential in response to constant current stimulus; both excitatory and inhibitory neurons exhibited both tonic and chattering type of spiking activity without clear preference across pharmacological conditions (see Fig S3-10).

We examined if there is any consistency in parameter changes across pharmacological conditions. We considered the pairs of successive pharmacological conditions, as before (e.g. ‘control’ and ‘AMPAR blocked’, ‘AMPAR blocked’ and ‘AMPAR & GABA_A_R blocked’ etc.). For each pair of conditions, we compared all possible pairs of considered models. We looked for the parameters that are bigger in one pharmacological condition than in other for at least 95% of model pairs. The parameters that satisfy this criterion are shown in Fig 5K. Among them, one is the post-spike reset threshold in the excitatory neurons (c in Izhkevich model), the parameter that affects spiking patterns and burstiness, one is a presynaptic model parameter (U_0_ in inhibitory to inhibitory synapses), and four are the peak conductance values of the synaptic receptors. The gAMPAR for inhibitory neurons tends to be smaller in control and NMDAR blocked condition than in NMDAR & GABA_A_R blocked condition. Both excitatory and inhibitory g_NMDAR_ are smaller in control than in AMPAR blocked condition. The dots in Fig 5K show the parameter values in the two compared models for the pharmacological conditions shown in two axes. For comparison, Fig 5L shows the same plot for the parameter a in excitatory cells, an example that did not pass the criterion used to select the parameters shown in Fig 5K. To conclude, the flexible protocol found versatile parameter sets that satisfied the multi-objective optimization criteria. In spite of variability in properties of individual neurons (Fig S3-10) network-wide synchronization during a bursting event resulted in relatively consistent time course of model variables (see S3, Fig S3-11).

### Section 4: A single model fitted to all pharmacological conditions reproduces changes in burst lengths and sizes across the conditions

An intrinsic assumption behind flexible protocol employed in sections 2 and 3 is that *in silico* pharmacological blocking of one synaptic receptor type (AMPAR, NMDAR or GABA_A_R) may also affect the conductances of other synaptic receptors as well as non-synaptic ion channel conductances through altered activation of certain cellular pathways. We tested whether the data from Fig 2 could be explained without this assumption. We sought for a single model capable of reproducing all experimental data from all conditions when subjected to *in silico* synaptic blocking, assuming that the blockade of the synaptic receptors only affected the underlying synaptic conductance. A naïve attempt at fitting the model to the control data and then blocking combinations of synaptic receptors in the already fitted model failed to reproduce the experimental data (see supplementary document S4). Clearly, a successful model requires fitting to a larger dataset including all pharmacological conditions of interest.

We developed constrained protocol to fit the same computational model as before (see Methods and Appendix 1) to the same experimental data by minimizing five objective functions that evaluate differences between the simulated and experimental BLs and BSs (see Appendix 2, Table C). The following were changed with respect to the previous sections: **1)** Fig 4 shows that flexible protocol cannot reproduce well the BSs in the NMDAR and GABA_A_R blocked condition, but also that it does not prohibit reproducing the changes in burst measures across conditions. Thus, the constrained protocol is designed to fit the BLs explicitly and the changes in BSs relative to control condition **2)** To ensure success in fitting the increased number of objectives we evaluated more parameter sets in each generation (400 instead of 200), as well as more fitting iterations (30 instead of 3). In order to reduce the computational cost, we abandoned fitting entire BL and BS distributions and focused on reproducing mean trends in the data (see Fig 4E-H). Estimating mean BLs and BSs required 3 model simulations in the training phase (100 model simulations were needed in the flexible protocol), while we used 10 model simulations in the test phase for more precise estimation of means. **3)** The distances between simulated and experimental data were computed as absolute values between means (Table B, Appendix 2). The number of completed generations was increased to 40-50 to ensure the convergence for all five objectives.

The design of constrained protocol, described above, ensured the evolution of all objectives as shown in Fig 6A. The evolution was the most rapid during the first 10 generations, after which more modest improvements were attained. During initial generations, the model fitting algorithm selected some parameter sets that produced ceaseless bursts composed of dense and long-lasting spiking activity in all pharmacological conditions. As these candidate models were not representative of the experimental data, and more importantly, greatly increased time needed for the objective function evaluation, we discarded them by setting a large value (1e8) for each objective whenever a ceaseless burst was encountered in one of the two first simulated conditions (NMDAR and GABA_A_R blocked or AMPAR and GABA_A_R blocked). These cases represented some of the maximal (and at times also the 75th percentile) objective function values during early generations, shown in Fig 6A. In the test phase, we re-estimated the objective function values for the final parameter sets using a larger number of repetitions (30 instead of 3). The three panels in Fig 6B represent the total error, the error made when reproducing BLs, and the error in reproducing BSs (normalized to control data) computed for all considered pharmacological conditions. The models are sorted in increasing order for each of the three types of errors (thus a model does not retain the same position in all three panels) to better illustrate the range of error values obtained for the best parameter sets.

**Fig 6:**
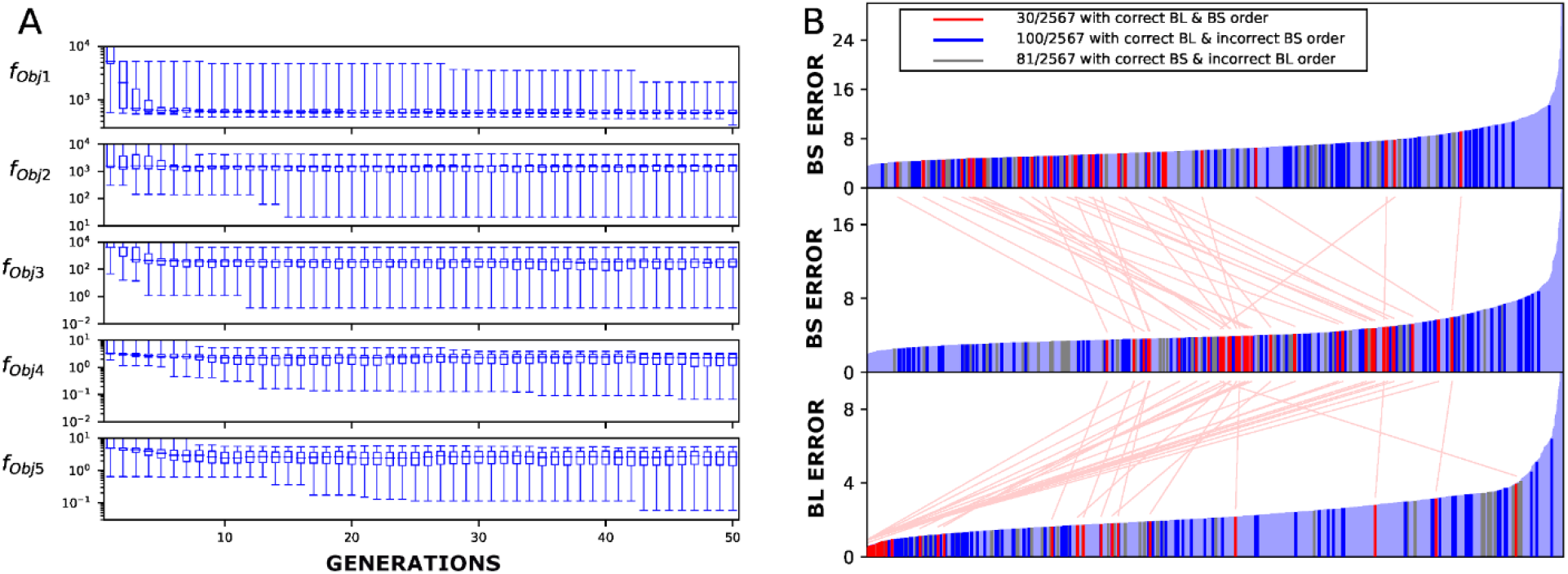
Model fitting using the ‘constrained protocol’. **A:** Evolution of the multi-objective optimization functions (see Appendix 2, Table C) pooled across 10 repetition of the model fitting. x-axis - the generation number, y-axis - the minima (bottom tick), 25th percentiles, medians (long tick), 75th percentiles, and maxima of the objective function value distributions **B:** Error values for the best fitted parameter sets from 10 repetitions of the model fitting. A total of 2567 unique parameter sets were obtained from the multi-objective optimization. The bottom panel shows BL errors (i.e. sums of BL errors divided by the standard deviation of the corresponding data), the middle panel shows BS errors, and the top panel shows the sum of the two. The dark blue bars represent parameters sets that predicted the accurate change of BLs across pharmacological conditions, the gray bars represent those that predicted the correct changes in BSs, and the red bars those that predicted both of them correctly. The light blue bars represent parameter sets that did not correctly predict changes in either of these measures. The fitness values are ordered by increasing value in each panel separately - the connections between the red bars show the identities.

The highlighted red, blue and grey bars show the models that managed to reproduce the changes across pharmacological conditions shown in Fig 2 for both burst measures (red), for BLs only (blue), and for BSs only (grey). The 30 models reproduced the correct order of both BLs and BSs. We ranked these 30 models based on the stability with which they reproduced the experimental data (based on how many individual repetitions fulfilled the conditions of acceptable fit), and the model that yielded the smallest total error (i.e. summed errors of predicted BLs and BSs normalized by the underlying STD in the experimental data) among the 3 most stable models was selected as the best fitted model.

The best fitted model is further analyzed in Fig 7. Rasterplots and population firing rates are shown in Fig 7A-E. The synaptic mechanisms governing the network-level dynamics are shown in Fig 7F-J, including the time courses of the available synaptic resources and the gating variables of AMPAR, NMDAR and GABA_A_R currents (small panels to the right also illustrate dynamics of the adaptation variable in model neurons). The best fitted model highlights possible mechanisms behind the experimental observations. Starting from the simplest condition, Fig 7E,J illustrates the dynamics in the networks with AMPAR-only mediated synaptic transmission. In these networks, a dense and relatively short network burst is generated, leading to a rapid use of most of the releasable synaptic resources (Fig 7J, pink) and a rapid build up of adaptation current (Fig 7J, right panel in black). Fig 7D,I show the activity in networks with AMPAR and GABA_A_R mediated synaptic currents. The presence of inhibitory currents leads to faster cessation of network bursts, compared to the networks with AMPAR-only currents. Consequently, the synaptic resources are not nearly as depleted and the adaptation current remains smaller than in networks with AMPAR-only currents. Fig 7C,H describes activity in networks with NMDAR-only mediated synaptic currents. These networks are characterized by much slower activation of neurons and thus a slower use of synaptic resources (Fig 7H, pink) and slower build up of adaptation current (Fig 7H, right, black) compared to the AMPAR-mediated networks. This allows the activity to be sustained over a longer time than in AMPAR mediated networks. Fig 7B,G shows the results of *in silico* AMPAR blocking. Compared to the simulations of AMPAR and GABA_A_R blocking (Fig 7C,H), the activation of the excitatory neurons is dampened (Fig 7B, right) due to the GABAergic currents caused by the inhibitory population. This leads to long (often non-ceasing within the simulation time) network bursts, where the depletion of synaptic resources (Fig 6G) is slightly less complete than in the absence of GABAergic currents (Fig 6H). Network activity in the control condition, shown in Fig 6A, is affected by all of the above contributions of different synaptic currents. The beginning of the network burst is similar to that in NMDAR-blocked networks, but NMDAR-mediated currents allow the network to be activated again, after a short period of silence caused by either GABAAR-mediated currents or synchronized afterhyperpolarization periods of excitatory neurons, or both (Fig F). This pattern is repeated several times, until the synaptic resources are largely depleted (Fig 6F, pink). The spike trains in control condition express a “bursts within a burst” pattern of activity. The high-activity periods are followed by (in a partly overlapping manner) high-frequency firing of the inhibitory population.

**Fig 7:**
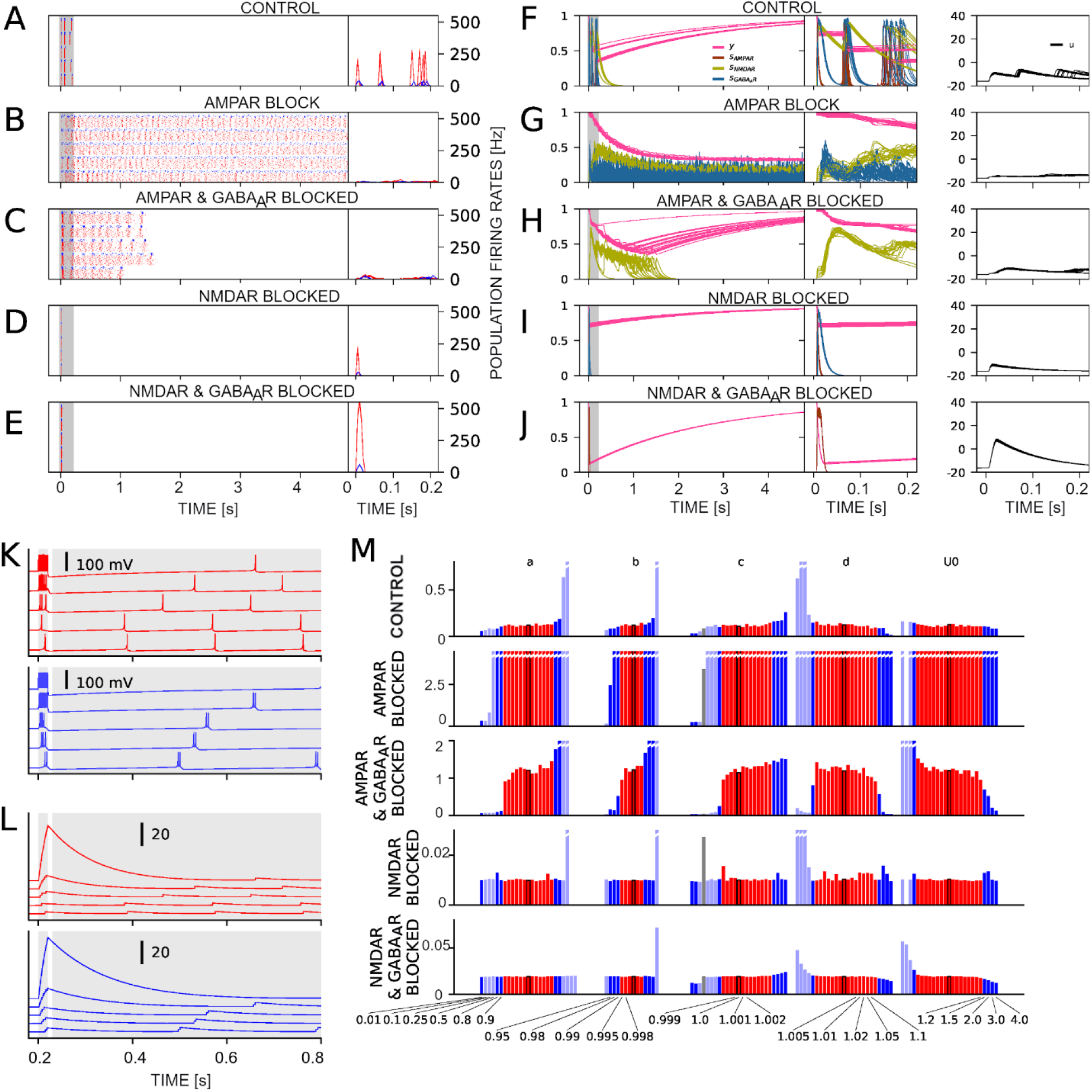
Synaptic, cellular and network activity in models fitted using the constrained protocol. **A-E:** Left: Spiking activity (red - excitatory neurons, blue - inhibitory neurons) in the best fitted model for different pharmacological conditions. x-axis – simulation time, y-axis - neuron index (80 excitatory, 20 inhibitory neurons). Five repetitions of the same model (with different random network topologies) are shown above each other. Right: Population firing rates zoomed in on the gray area in the left-hand panels. The data from five simulations are overlaid on each other. **F-J:** Time courses of the synaptic gating variables and fraction of the releasable synaptic resources. The y-axis shows the average quantity over excitatory (*s*_*AMPAR*_, *s*_*N*MDAR_, *y* ^*glu*^) or inhibitory 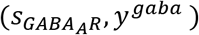 synapses. Data from five simulations are overlaid on top of each other. The right-hand panels zoom in on the gray area in the left-hand panels. **K-L:** Single neuron responses to a DC pulse with an amplitude 0.5 (bottom), 1.5, 4.5, 15, or 100 nA (top), starting at time t=200ms and lasting 20ms, followed by a long DC pulse with amplitude 0.5nA starting at 230ms. The upper panel (K) shows the membrane potentials of a single model neurons, and the lower parts of the panels show adaptation current variable (u in Izhikevich model; red-excitatory, blue-inhibitory). **M:** BL sensitivity to variation of the best fitted model parameters. Parameters a, b, c, d, (Izhkevich neuron model) for excitatory neurons, as well as *U*_0_ are shown. The five panels correspond to BLs (in ms) in the five different pharmacological conditions. One parameter at a time was multiplied with a coefficient ranging from 0.01 to 4 (see all tested coefficients at the bottom of the figure). The BLs obtained from the best fitted model are shown in the middle of each plot (bars with the thick black edge). The color coding is the same as in Fig 5B: red bars - parameter sets that reproduced the correct order of both BL and BS, dark blue bars - correct BL but incorrect BS, grey bars - correct BS but incorrect BL.

We further examined the membrane potential and the adaptation variable of the excitatory and inhibitory neurons in the optimized model in Fig 6K,L. This figure describes the responses of single model neurons (parameters given in Table I, Appendix 1) to short square-pulse currents of different amplitudes followed by long square-pulse current of amplitude 0.5nA. Both excitatory (red) and inhibitory (blue) neurons expressed a behavior where a strong build-up of the adaptation current (Fig 6L) delays the reactivation of the neuron (Fig 6K). The delay caused in the inhibitory neuron was somewhat larger than in the excitatory neuron due to the slower decay of the adaptation variable (Fig 6K-L). Moreover, the inhibitory neuron exhibited chattering firing behavior in response to prolonged stimulus due to its large membrane-potential reset value while the excitatory neuron fired single action potentials (see Fig 7K, for time > 0.4s).

To ensure the robustness of the best fitted model and to analyze the contribution of different model parameters we performed the same sensitivity analysis as before. Fig 6M summarizes sensitivity to the excitatory neuron model parameters (a, b, c, d) and the parameter controlling the depth of presynaptic depression (*U*_0_) while all other parameters as well as the sensitivity of BS are shown in supplementary document S5, Fig S5-1, S5-2, S5-3. Similarly as in Fig 4N-R, the BL was most sensitive to neuron model parameters, in particular the firing threshold parameter b for the excitatory neurons, where a modest change of 1% changed the reproduced order of burst lengths. The sensitivity to Izhkevich model parameters was particularly large in the GABA_A_R-blocked conditions, where a 5% decrease in b and c in the excitatory neurons decreased the burst lengths by >80%. Sensitivity to *U*_0_ had a large effect on the two conditions without inhibitory neurotransmission (NMDAR and GABA_A_R blocked, AMPAR & GABA_A_R blocked) suggesting that synaptic depression played a large role in cessation of the bursts in these conditions (Fig 6J,H) and a smaller role in other conditions (Fig 6F,G,I). The BS sensitivity to parameter changes followed similar trends across pharmacological conditions (supplementary document S5, Fig S5-1). The neuron model parameters for the inhibitory population had smaller effects: apart from the firing-threshold parameter b, the parameters could be altered by ±20% without affecting the qualitative reproduction of the data (S5, Fig S5-2, S5-3). The BLs (S5, Fig S5-2) and BSs (S5, Fig S5-3) were also robust against small changes in synaptic conductance parameters.

In summary, a single spiking network model, adjusted to all experimental data using the constrained protocol and subjected to *in silico* blocking of synaptic receptors can reproduce the changes in BLs and the relative changes in BSs compared to control seen in the experimental data in Fig 2. However, there could be other parameter sets that reproduce the data by alternative fundamental mechanisms – for example, in one model the burst cessation of Fig 6C and 6E could be mediated by strong depression and in another model by strong adaptation currents. In the following section, we examine which properties are indeed important for qualitative reproduction of the experimental data and which ones can be compensated by other mechanisms.

### Section 5: Computational model predicts the importance of single-neuron bursting/adaptation but not of heterogeneity across neuronal population for reproducing the experimental data

We employed the constrained protocol from the previous section to further examine individual cellular, synaptic and network mechanisms and assessed how significant they are for successful reproduction of the data from Fig 2. In Table 2, we compiled a list of mechanisms that might have a fundamental role in reproducing the features of network bursts. To understand how important each of the neuronal or synaptic mechanisms is, we ran the optimization with and without the considered mechanism. Our rationale was that if the mechanism were crucial for the phenomena observed in the experiments, the data would not be reproduced by the model lacking the mechanism while it would be reproduced by the model containing the mechanism.

**Table 2:** List of abstractions of neuronal and synaptic mechanisms and the implementations for their exclusion.

| Mechanism | Hypothesized effect on network dynamics | Parameter assignments that exclude the mechanism |
| --- | --- | --- |
| Short-term depression | The probability of initiation or maintenance of a network burst depends on previous activation time of the synapses [55] This may make the distinction between short and sustained bursts more robust. | $U_0^{glu/gaba} = 0$ for all synapses in the network model |
| Adaptation and burstiness of single neurons | Bursting and chattering types of single-neuron dynamics may provide a period of silence or lower firing activity after a high-activity period. This could help in synchronization of the population [39,61], leading to an additional means of ceasing the network burst [62]. | $a = 0.02, b = 0.254,$<br>$c = -65mV, d = 2 \frac{V}{s}$<br>These parameters produce tonic firing [39]. They are used for all model neurons including excitatory and inhibitory neurons. |
| Variability in intrinsic neuron dynamics | The spiking times within bursts become more variable and certain cells fire longer trains within network bursts than other cells. This may facilitate (or hinder) the emergence of another period of burst activity [62], leading to sustenance (or cessation) of the spiking activity. | $\sigma_a = \sigma_b = \sigma_c = \sigma_d = 0$<br>The variability in neuronal parameters is set to zero for all model neurons. |

Table 2 lists the considered mechanisms, their putative effect on network dynamics, and the parameter choice that effectively excluded the mechanism *in silico*. The first mechanism we considered was the short-term depression that accounts for depletion of presynaptic resources at each transmitted spike. In the network bursting regime, where many neurons co-activate to produce relatively short intervals of dense activity rapid depletion of presynaptic resources might lead to faster termination of bursting [55]. We excluded this mechanism by setting the parameter *U*_0_ to zero for each synapse, both glutamatergic and GABAergic synapses, which resulted in a model with infinite presynaptic resources. In a similar fashion as before, we fitted the model to all experimental data. Surprisingly, some of the fitted parameter sets reproduced the changes in BLs and BSs induced by blocking of synaptic receptors (see supplementary document 5, Fig S5-4). The total error among the acceptable 12 parameter sets was larger in this setting than in the fittings shown in section 4 where synaptic depression was included (S5, Fig S5-5, 5.8 ± 0.7 vs. 5.3 ± 1.0, p=0.02<0.05 U-test, n=12 vs. 30). The simulations obtained with the best parameter set (i.e. the smallest total error) without synaptic depression are illustrated in S5, Fig S5-5 (corresponding to Fig 7 in the previous section). These simulations reproduced the data less robustly, since repetitions of the test phase with parameters with little or no change sometimes failed to reproduce the correct BL and BS relations (S5, Fig S5-5M).

When we disabled single-neuron adaptation as described in Table 2 (i.e. used the tonic spiking version of the Izhikevich model for all neurons), we could not reproduce the relations between burst lengths and sizes (S5, Fig S5-6). By contrast, when we allowed single-neuron adaptation but disabled the variability of Izhikevich model parameters within the population (by setting all four parameter variances to zero) and fitted the model containing homogeneous excitatory and homogeneous inhibitory population we were able to reproduce the orders of burst lengths and sizes (S5, Fig S5-7). The total errors from the 42 acceptable parameter sets were not significantly different than the acceptable parameter sets obtained in section 4 by allowing the variability (5.0 ± 0.9 vs 5.3 ± 1.0, p=0.08>0.05, n=42 vs. 30). The properties of the best among these models are presented in Fig S5-8. This result suggests that the advantage of having a freely chosen magnitude of variation in the intrinsic neuron parameters is minor compared to the additional difficulty in the optimization imposed by the increase in parameter space. We reached the same conclusion using flexible fitting protocol, which aimed at reproducing both the mean trends and the variability in experimental data (see Fig 5A-J).

It suffices to show a single acceptable model to demonstrate that a certain set of mechanisms can reproduce given data with required accuracy, but it is harder to be sure that a certain set of mechanisms is incapable to reproduce it. For further evidence that adaptation of neurons was necessary, we re-ran the optimization without single-neuron adaptation using more relaxed limits of model parameters. Namely, we increased the maximum search values of all synaptic conductance parameters by 150%. This optimization also failed to produce acceptable fits (Fig S5-9). We went on to study whether single-neuron adaptation of excitatory or inhibitory neurons alone was sufficient to reproduce the correct orders of burst lengths and sizes. Acceptable fits were found both when excitatory neurons alone were allowed single-neuron adaptation (Fig S5-10, S5-11) and also when inhibitory neurons alone were allowed the adaptation (Fig S5-12, S5-13). In the former case, the total error was significantly larger than in the optimization of section 4 (7.7 ± 0.1 vs 5.3 ± 1.0, p=0.03<0.05, n=2 vs. 30), but in the latter case the difference was non-significant (7.1 ± 0.3 vs 5.3 ± 1.0, p=0.06>0.05, n=2 vs. 30).

The importance of different mechanisms suggested here relies on the performance of the used multi-objective optimization algorithm. It has been shown that NSGA-II does not perform optimally with a large number of objective functions (more than two or three [51]). To confirm our results, we re-optimized the parameters using NSGA-III algorithm, which outperforms NSGA-II when multiple objective functions are used [60]. We used 5 repetitions for each set of mechanisms. We obtained similar results with NSGA-III when it comes to reproducing the correct order of burst lengths and burst sizes: acceptable models were obtained when all mechanisms were present (Fig S5-14), and also when synaptic depression (Fig S5-15) or variability of Izhikevich parameters (Fig S5-16) was excluded, but not when only tonically firing neurons were used (Fig S5-17).

Despite the capacity to reproduce the experimental data, the characteristics of the accepted models may be different when different sets of mechanisms were included. To explore the strategies in which the models with different sets of mechanisms achieved the acceptable fit to the data, we compared the parameter value distributions of all acceptable parameter sets across the tested sets of mechanisms (all mechanisms present, without synaptic depression, without variability in neuron model parameters, without adaptation, and without adaptation in excitatory or inhibitory neurons). We found no large differences between the parameters from acceptable fits (Fig 8A). The smaller range of parameters in the models lacking adaptation in excitatory or inhibitory neurons is most likely explained by the smaller number of acceptable fits (Fig 8B). Larger numbers of repetitions for each optimization should be made to draw further conclusions. We also quantified the prevalence of chattering-type firing activity in excitatory (Fig 8C) and inhibitory (Fig 8D) neurons among the acceptable models by simulating the response of the single-cell models to a square-pulse current of 1.0 nA for 2 seconds: the neuron was considered chattering if a burst of at least 2 action potentials, separated by a maximum of 15 ms, was encountered after 500 ms of stimulation. The excitatory neurons in general displayed chattering firing less often than the inhibitory neurons, with the exception of the two last sets of mechanisms where one of the two populations was fixed to be tonically firing (Fig 8C-D). Finally, there were differences in whether the predicted activity for AMPAR & GABA_A_R-blocked only showed ceaseless burst activity or whether the firing activity died out in at least some of the 20 repetitions (Fig 8E). When synaptic depression and single-neuron adaptation were included, 3% or 21% of acceptable parameter sets in optimization tasks where all mechanisms were included (3%) or where variability of Izhikevich parameters was excluded (21%) contained ceasing bursts also in this condition (Fig 8E). By contrast, when synaptic depression was excluded or when one of the neuron populations had fixed neuron parameters inducing tonic firing, all optimization tasks only produced acceptable parameter sets that led to ceaseless bursting activity in this condition (Fig 8E). This suggests that synaptic depression may be important for quantitative reproduction of the long-lasting bursting activity as observed in the experimental data although it is not needed for qualitatively correct reproduction of the data.

**Figure 8:**
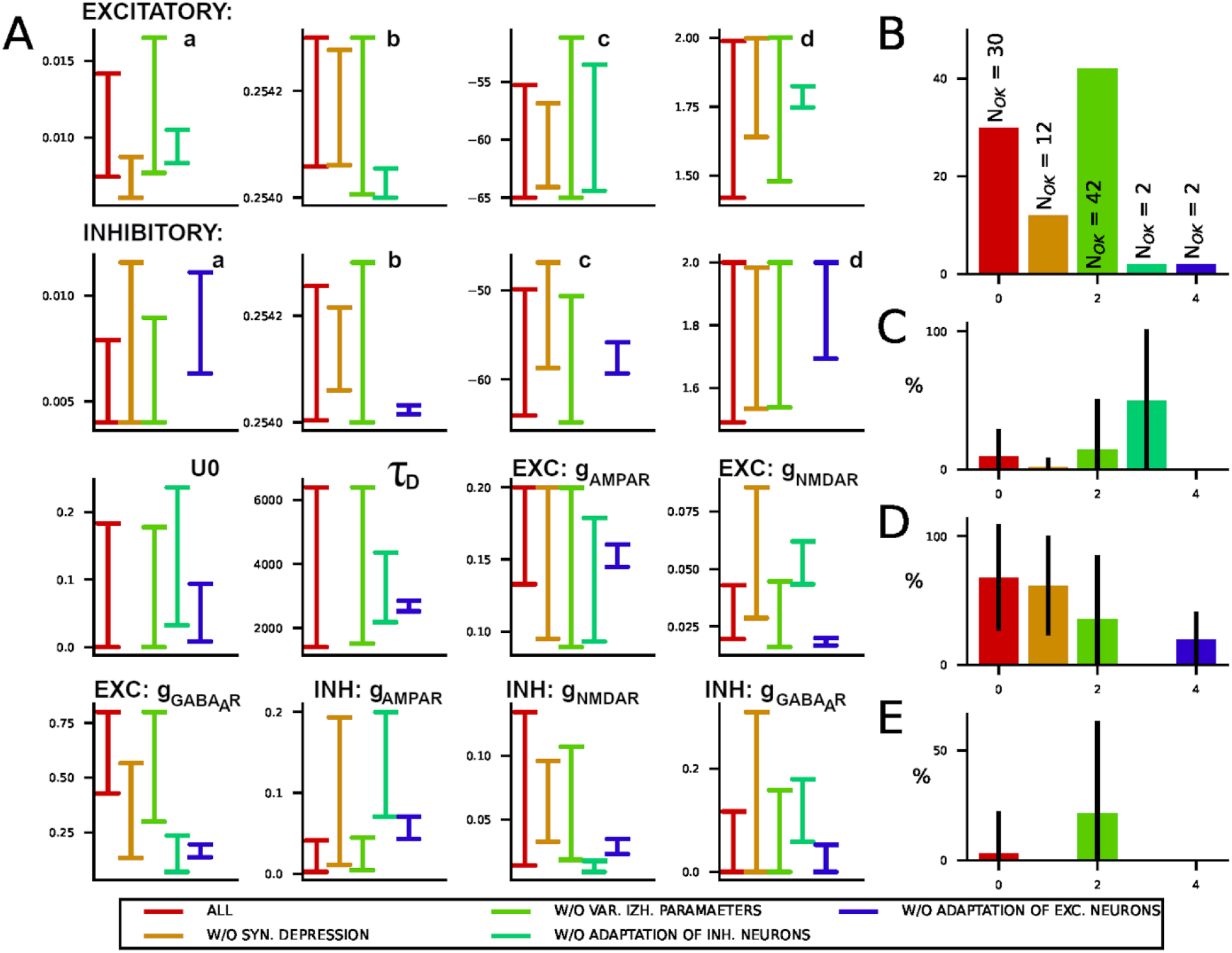
Performance of constrained protocol when different mechanisms are included into the fitted models. **A:** Parameter ranges of acceptable parameter sets obtained from the five multi-objective optimization tasks with different mechanisms included into fitted models. **B:** Numbers of acceptable fits obtained from the five tasks. **C:** Percentage of acceptable fits where the excitatory neuron showed chattering-type of firing behavior in a single-cell simulation where the neuron was stimulated with a 1.0nA long-lasting square-pulse current (out of all acceptable fits). **D:** Percentage of acceptable fits where the inhibitory neuron showed chattering-type of firing behavior. **E:** Percentage of acceptable fits where the bursts in the AMPAR & GABAAR-blocked condition ceased (in at least one out of the 20 repetitions) within simulation time (5 sec).

In summary, we tested a range of mechanisms embedded into the computational model and assessed how crucial they are for the success of reproduction of the experimental findings.

We found that the exclusion of single-neuron adaptation hinders the model’s capacity to reproduce the data, and exclusion of synaptic depression or partial exclusion of adaptation exclusion from excitatory or inhibitory neurons alone increases the total error but does not hinder qualitative reproduction of the experimental data. On the contrary, excluding the variability in the neuronal model parameters did not impair the model optimization.

## Discussion

In this paper, we present a new data-driven modeling formalism to study how the excitatory and inhibitory synaptic drives modulate the structure of spontaneous network bursts. In our earlier publication [24] we quantified the changes in burst structure induced by blocking of individual or combinations of synaptic receptors (AMPAR, NMDAR, GABA_A_R). In this new computational study, we identify the synaptic, cellular and network level mechanisms sufficient to reproduce the experimental findings. We demonstrate that the sole blocking of synaptic transmission, consistent with the pharmacological protocols in [24], can explain the changes in burst structure seen across the pharmacological conditions. When we adjust both the synaptic transmission and the cellular excitability to each pharmacological condition, we can faithfully reproduce the quantitative description of the burst structure. To the best of our knowledge, our study is the first to employ a systematic data-driven computational modeling to combine the spiking network models with the experimental network-level MEA data collected under several pharmacological conditions.

The proposed data-driven modeling formalism consists of the following steps. **1)** In [24], we measured and characterizing various aspects of network burst structure in dissociated cortical cultures *in vitro*. Here, we focus on two measures of network activity, namely the burst length (BL) and burst size (BS). This was done for each of the five pharmacological protocols, namely the control, blocking one type of glutamatergic receptors (AMPAR or NMDARs), and blocking of GABA_A_Rs in AMPAR- or NMDAR-blocked networks **2)** Constructing the spiking network model that incorporates the considered synaptic receptor types as well as the reduced set of cellular mechanisms. **3)** Formulating multi-objective optimization tasks and solving them using the algorithm from the literature [50,52,58,60]. **4)** Defining two complementary model fitting protocols. The first, flexible protocol, fitted a separate model to each pharmacological condition and obtained a good fit for experimentally recorded BLs and BSs in 4/5 conditions, and for BLs in the remaining condition. The second, constrained protocol fitted one model to all conditions and reproduced the relative changes in BLs and BSs across conditions. **5)** Using the constrained protocol we explored which cellular and synaptic mechanisms are necessary to reproduce the findings from [24]. In what follows, we present our findings and challenges related to each of the steps 1-5. We finish with discussion about applicability of our approach to different models and experimental preparations.

We measured and characterized various aspects of network activity in our earlier study [24]. Here, we opted to focus on two measures of network activity, the burst length and the burst size. Increasing the number of burst measures improves the description of the experimental dataset but also increases the number of criteria for model optimization and the complexity of the optimization task. We used a conventional burst detection method based on statistics of spike times to identify network bursts both *in vitro* and *in silico*. Alternative methods are available in the literature, but to the best of our knowledge, none of them is fully automatic and perfectly accurate (for a systematic evaluation of burst detection methods see [63]). Our modeling approach requires fully automatic *in silico* burst detection done inside the model fitting algorithm. Occasional detection errors affect model selection in random algorithm steps, and then propagate through all successive steps in multi-objective optimization and may impact the results at each stage of the analysis. Here, we applied a pragmatic solution to the problem, we employed a conventional and fast burst detection method of average precision, but we incorporated additional steps into model fitting to remove all ‘suspicious’ models early on and prevent error propagation. This tradeoff gave good result and the obtained *in silico* bursts had structure consistent with the experimental ones.

The neuronal network model used in our study consists of spiking neurons randomly connected through synapses that incorporate the short-term presynaptic dynamics and phenomenological models for AMPA, NMDA and GABA_A_ receptors. This type of spiking networks have been used earlier for theoretical studies of network bursts *in vitro,* see [45,46,53,64–73] and for a review [74]. In addition, the mean-field approximation was also used to study network bursts *in vitro* in [75–77]. We adopted a spiking neuron model [39,40] that permits easy tuning of cellular excitability, sensitivity to synaptic inputs and implicitly accounts for some conditions in the extracellular environment, but does not explicitly represent cellular mechanisms. It is a computationally light model that can be efficiently optimized to reproduce properties of the extracellularly recorded spiking patterns (e.g. tonic spiking vs bursting, the number and frequency of spikes per neuron). To our best knowledge, our study is the first that combines multiple pharmacological protocols, data analysis and model fitting to the data to construct spiking network models. An earlier study proposed a top-down model fitting to a single objective and single (control) condition that searches for the global optimum via systematic analysis of the parameter space for mean-field and spiking network models in [76]. In the absence of biophysically detailed models specific for dissociated cortical cultures, we adopted the generic model described above however our approach can easily be extended to incorporate additional biophysical model complexity.

The here developed neuronal network model consists of 100 model neurons, 80 excitatory and 20 inhibitory. This ratio between the number of excitatory and inhibitory neurons is supported by experimental observations [78–80]. A typical dissociated cortical culture consists of 10.000 - 100.000 cells [81]. A standard 60-electrode MEA can be expected to capture the activity of 2-3 proximal neurons per electrode [81], which gives 120-180 recorded neurons in total. Thus, the number of neurons in our model heavily underestimates the size of cortical cultures, but only slightly the number of neurons recorded using the standard MEA. However, in a small model the number of synaptic inputs to each neuron is largely underestimated. This is likely compensated by stronger synaptic coupling between model neurons, which might reflect in altered variability in input and output spiking patterns of each neuron. On the network level, this might lead to shorter and more abrupt synchronous events within network bursts seen in some of our results (see Fig 4I-M, Fig 7A-E), instead of longer and more spread out activity found in the experimental data.

The structure of network bursts might also be affected by properties of connectivity scheme, which is selected to be a generic random connectivity in our model. In dissociated cultures *in vitro*, computational studies emphasize the functional role of complex connectivity [45,68] and computational modeling predicts the non-random connectivity [82]. However the experimental studies supporting these computational predictions are scarce (but see [23,79]). Careful mapping of connectivity in dissociated cultures is technically challenging. Having in mind these arguments, exploring the role of complex connectivity is omitted from this study and is left to future work. The connectivity scheme in our models is described by a single parameter, the probability of connection between a pair of neurons, which is set to 20% similarly as in earlier studies (see e.g. [69]). In a network of 100 neurons connected with 20% probability, on average 1980 synaptic contacts are chosen out of 9900 potential contacts (self-connections excluded), which gives 10^2149.5^ possible connectivity schemes, and even more given that the number of synaptic contacts randomly fluctuates around 1980. The choice of connectivity defines the feasible pathways of activity propagation in the model, which affects burst structure and induces randomness to BL and BS statistics (see rasterplots in Fig 4I, and the inset of Fig 4I). The BL and BS statistics is estimated from multiple simulations of the same model but with a new randomly generated connectivity in each simulation. This approach is consistent with our experimental data, namely we obtain the BL and BS statistics by pooling data from 13 cell cultures. The neuronal connectivity varies from culture to culture and by pooling data we randomize the connectivity of our experimental preparation. An alternative to our approach would be to fit one model with the fixed connectivity to the data collected from one culture. However, from the model fitting perspective, this choice would lack generalizability. Each fitted model would represent well the data from one culture, but would not be able to generalize well to the data from other similar cultures. Thus, in this study we adopted a simple connectivity scheme, which is treated as a source randomness in the model.

The results are obtained through extensive simulations of models selected by a genetic algorithm, estimating statistics of *in silico* BLs and BSs, and evaluating goodness of each model by comparing *in silico* and *in vitro* data according to multiple criteria. This methodology is common in computational neuroscience and was used for data-driven modeling of individual neurons (e.g. see [50,52]) and recently for detailed synaptic mechanisms [59]. To the best of our knowledge, it has not been used to integrate spiking network models with the experimentally measured network-level activity from multiple pharmacological conditions before. The lower-dimensional more computationally tractable mean-field models were fitted to the data from dissociated cell cultures in control condition [76], and spiking network models were compared to the data from multiple pharmacological protocols but without model fitting (see for example [45,69]). Our adopted method allowed to integrate data from multiple preparations and imposed no constraints on selecting the modeled mechanisms. Based on this approach, we developed two model fitting protocols.

The flexible protocol provided one model per pharmacological condition by performing *in silico* blocking of receptors and then fitting the remaining parameters to the data from that one condition. By fitting each pharmacological condition separately, we compensated for possible changes in synaptic, cellular, and extracellular properties induced by synaptic. The obtained five models were able to reproduce the entire distributions of the experimentally measured BLs and BSs for 4 out of 5 pharmacological conditions and the distributions of BLs for the fifth condition.

The constrained protocol fitted one model to the data from all five pharmacological conditions, thus assuming that blocking affects only the targeted receptor. Here, we blocked synaptic receptors during model fitting, i.e. for each model evaluated during fitting we blocked the five considered combinations of synaptic receptors and obtained five sets of BLs and BSs. All five sets were then used to construct objective functions and evaluate similarity between *in silico* and *in vitro* data. This model successfully reproduced the increase or decrease in BLs and BSs across pharmacological conditions. To conclude, we were able to reproduce the exact experimentally measured BL and BS distributions when fitting one model per pharmacological condition, and thus allowing the model to adapt to each condition separately. However, for reproducing the relative change in BLs and BSs across pharmacological conditions, one model fitted to all data and subjected to *in silico* synaptic receptor blocking was sufficient, suggesting that the relative changes across conditions seen in Fig 2 might result from the changes in synaptic transmission.

We further tested under what conditions this conclusion holds. Our tests suggest that the single-neuron adaptation might be an important mechanism. After removing this mechanism from the model we were not able to reproduce the experimental data, at least for the model and the parameter ranges considered in our study. The neuronal adaptation is known to be important for maintaining the appropriate level of neuronal excitability [39,40] and might have important contribution to network burst termination [73]. In addition, in our fitted models, the presence of adaptation supported the emergence of specific spiking patterns. Namely, in the acceptable parameter sets the inhibitory neurons tended to exhibit single-cell bursting while the excitatory neurons were mainly tonic spiking (Fig 7K, Fig 8C-D). On the contrary, we were able to reproduce the experimental data using the models without the pre-synaptic depression, however the models equipped with this mechanism exhibited more robust burst termination and were less likely to produce ceaseless bursting. Similarly, introducing variability of model parameters was not needed for successful model fitting. This was also found when using the flexible protocol, which aimed at reproducing not only mean trends in the data but also the exact *in vitro* recorded BL and BS distributions that typically include significant variability in the data (see Fig 5).

Finally, we inspected the parameters obtained from model fitting. As expected, when using genetic algorithms, diverse parameters produced successful models (Fig 5, Fig 8). Genetic algorithms tend to maximize the variability in the selected candidate parameter sets in order to optimize the exploration of the parameter space. These methods do not guarantee a global solution to the fitting task, so a number of fitting trials might be reaching possibly very diverse local optima. Additionally, by fitting a spiking neuronal network model to the experimental data we projected much richer dynamics of the biological *in vitro* networks onto a lower-dimensional set of equations, and consequently the solution to the fitting problem might be non-unique. We explored the changes in individual model parameters across pharmacological conditions, and found, at least in the model and parameter ranges considered in this study, some parameters that change consistently across pharmacological conditions (see Fig 5K). Among them, the peak conductances of the synaptic receptors where more common than other parameters.

Focusing on the automatic fitting of models of low computational complexity that incorporate only the minimal set of cellular, synaptic and network mechanisms led to excluding many other biophysical properties that might affect network burst dynamics. For example, the blockage of NMDARs significantly reduces the calcium influx to the neurons, which can lead to (de)phosphorylation and exo/endocytosis of many ion channels and thus alter also the intrinsic neuron excitability [83]. Blockage of AMPARs and GAB_A_Rs may have similar impacts through the mechanisms of homeostatic plasticity [83]. Non-linearities in dendritic processing, such as presence of a hot zone of calcium channels and its coupling to SK channels, might have significant effect on spike timing and integration of synaptic inputs [52] with potential to modulate burst structure. Moreover, non-neuronal cells present in cell cultures, including astrocytes, respond to synaptic activity and can signal back to neurons [84]. Finally, the high-activity firing seen during network bursting can alter the extracellular ionic concentrations, leading to altered intrinsic excitability of the neurons [85]. Glial cells are also known to affect the extracellular ionic concentrations and consequently the neuronal excitability, e.g. by regulating potassium concentrations [86–88] but also the concentration of other ions in the extracellular space. Fitting an individual model using flexible protocol to each pharmacological condition allows adapting excitability of model neurons and implicitly accounting for some of the listed mechanisms. However, even this approach resulted in excessive BSs in NMDAR and GABA_A_R blocked condition. The BSs in this condition were correctly reproduced when transmission was allowed also over additional synaptic mechanisms. In our model, that was done by allowing active GABAARs or GABAARs and NMDARs, which should be blocked at least partially according to the pharmacological protocol. Thus, this result might also point at the lack of some other crucial mechanisms.

The mechanisms listed above can also be explored by our model fitting framework, as the framework does not impose hard limitations to model complexity or to the considered experimental data. Including additional synaptic mechanisms or more than two cell types is straightforward. Incorporating more complex neuron types, e.g. a two-compartment neuron model to account for dendritic mechanisms, is also possible. However, a more complex model would increase the computational load of model fitting which should be taken into account when defining the model fitting protocol. For example, fitting the mean trends in burst measures, as done in our constrained protocol, is less computationally demanding than fitting their distributions, as done in our flexible protocol. To illustrate this, the flexible protocol needed about three and a half days to evaluate 200 models in each of 31 generations, while the constrained protocol needed two days to evaluate 400 models for between 40 and 50 generations. Incorporating different measures of network bursting activity (see other measures considered in [24]) will not increase the computational load of individual model evaluation but might require evaluating more parameter sets and more generations to achieve good fitting. Including the intra-burst measures (e.g. the duration of intra-burst intervals, or the average spiking activity between bursts) would increase the simulation time of individual models. In this study, we simulated each model as long as needed to reproduce a single burst in any pharmacological protocols. Estimating *in silico* inter-burst intervals would require simulating one burst and as well as a period until the next burst, which is substantially longer but populated with sparse activity. Finally, the available data should be considered, a model fitting task exploring a large parameter space or fitting many objectives requires sufficiently big training data set.

In summary, we propose a new data-driven model fitting framework to develop rodent and human models of neural networks in a systematic manner using experimental data. Our results using this framework advance understanding of the complex interplay between cellular and synaptic mechanisms in shaping network activity. Additionally, our study emphasizes the need for systematic data analysis and careful evaluation of the model fitting protocol against the available data. While here we focused on the analysis of *in vitro* data, our approach can be applied to similar analyses of network dynamics *ex vivo* and *in vivo* given the appropriate experimental data is available.

## Supporting information

Appendix 1, model

Appendix 2, objective functions

Supplementary data 3, flexible protocol

Supplementary data 5, constrained protocol

Supplementary document 6, description of data and code

## Acknowledgements

This research has received funding from the Academy of Finland and ERA-NET NEURON (https://www.aka.fi/en/, decision Nos. 297893 and 318879 to M-LL) and (decision numbers 330776 and 336376 to T.M.), and the European Union’s Horizon 2020 Framework Programme for Research and Innovation (https://ec.europa.eu/programmes/horizon2020/en) under the Specific Grant Agreement Nos. 785907 (Human Brain Project SGA2) and 945539 (Human Brain Project SGA3) to M.-L.L. Partial support from UiO: Life Science Convergence Environment (4MENT) and Research council of Norway (248828) to T.M is acknowledged. H.T. acknowledges support from Graduate School of Tampere University of Technology, Tampere Graduate School in Information Science and Engineering, Finnish Foundation for Technology Promotion, Finnish Cultural Foundation (Central and Pirkanmaa Regional funds), and Finnish Brain Foundation sr. The CSC – IT Center for Science, Finland and TCSC – Tampere University Computing Center provided the computing resources that made our study possible.

## Author Contributions

J.A., T.M.-M., H.T. and M.-L.L. jointly contributed to conceptualization, formal analysis, investigation, selection of methodology, prepared the resources necessary for this study and contributed to validation of the obtained results. H.T. prepared the experimental data and managed data curation. J.A. and T. M.-M. developed the methodology and the code needed for this study, obtained and visualized the results. J.A. prepared the initial version of the manuscript and all authors contributed to preparation of the final version of the manuscript. J.A. and M.-L.L. were responsible for project administration. M.-L.L. supervised the project and acquired funding needed for this study.

## Supporting information

### Appendix 1 - Details of model description

Details of the considered computational model presented in a standardized format according to [54]. Tables A-F contain description of various parts of the model according to the utilized format. Table G lists model parameters that were kept fixed across all fitting trials. Table H lists parameters for the best models obtained using flexible protocol. Table I lists the best parameters obtained using the constrained protocol.

### Appendix 2 - Burst measures, distance measures, objective functions

Details of the objective functions used in two applied model fitting protocols. Table A summarizes the selected network burst measures - the burst length (BL) and size (BS). Table B presents distance measures used to compare *in silico* and *in vitro* data. Table C gives two objective functions used in flexible protocol and five objectives of the constrained protocol.

### S3 – Additional results for flexible protocol

Fig S3-1 illustrates the convergence of the test phase for the flexible protocol algorithm convergence in the training phase is shown in Fig 3. Fig S3-2 shows control tests for NMDAR and GABA_A_R blocked protocol. Allowing synaptic transmission through either GABAAR or both blocked receptors permits fitting of both BLs and BSs. Fig S3-3 and Fig S3-4 show the full model, with all synaptic receptors being active, allows fitting of both BL and BS for each pharmacological condition. Fig S3-4A,B shows 10 spike trains and population firing rates corresponding to 10 best models from 10 model fitting trials (i.e. each rasterplot is produced by a model with different parmeters). Fig S3-4C shows BL and BS distributions for these models as well as the range of JSD values obtained when comparing these distributions to the experimental data. Unlike in other figures, we here show results of individual fitting trials, instead of one best result from all trials, as an illustration of model fitting procedure and variability of the obtained results. Fig S3-5 complements Fig 4N-R by showing sensitivity analysis for all other burst measures and pharmacological conditions. Fig S3-6 and Fig S3-7 analyze the model with no variability in Izhkevich model parameters but with variability in pre- and post-synaptic parameters. Fig S3-8 and Fig S3-9 analyze the model with no variability in any of the parameters. Fig S3-10 illustrates diversity of spiking patterns in model neurons fitted using flexible protocol. Fig S3-11 illustrates dynamics of model variables for the five pharmacological conditions.

### S4 - Naïve approach in fitting one model to all pharmacological protocols

We tested how well a single model fitted to the control data only can account for all pharmacological protocols. A model fitted to the control condition was subjected to *in silico* blocking consistent with our pharmacological protocols. Evaluating similarity between *in silico* and *in vitro* data demonstrated that this straightforward approach to model fitting cannot account for all considered conditions. This test served as the motivation for the work presented in sections 4 and 5.

### S5 – Additional results for constrained protocol

Fig S5-1,2,3 complement Fig 7M and show sensitivity analysis for all other model parameters not presented in Fig 7M. Fig S5-4,5 analyze model in which presynaptic depression is excluded (and thus the model has infinite presynaptic resources). Fig S5-6 shows fitness error values for the model with single-neuron adaptation removed in both excitatory and inhibitory neurons. Fig S5-7,8 analysis of the model with no variability in model parameters. Fig S5-10,11,12,13 analyses fitting of models where single-neuron adaptation is excluded in either excitatory or inhibitory neurons (but not in both). Fig S5-14,15,16,17 shows control model fittings carried out using an alternative multi-objective optimization algorithm, namely in these fittings NSGA-II is replaced by NSGA-III algorithm which is more powerful when optimizing more than two objectives (five in our case). We obtained consistent results using both algorithms.

### S6 - Data and Code

Data and code used in this study are available in ModelDB, access to this repository will be provided to the reviewers. The repository will be made open access upon acceptance of the manuscript.

