## Appendix 1, model for "Analysis of cellular and synaptic mechanisms behind spontaneous cortical activity *in vitro*: Insights from optimization of spiking neuronal network models"

### Appendix 1: Details of computational model, equations, fixed parameters, optimized parameters

The spiking neuronal network model is described in a standardized format proposed in Nordlie et al. (2009) in **Tables A1-A-F. Table A1-D** gives all model equations in a compact format with emphasis on model parameters. We described a neuron model according to Izhkevich 2003, 2004. The synaptic model consists of the short-term depression on the presynaptic side (according to Tsodyks et al., 2000), and the synaptic conductances for AMPAR, NMDAR and $GABA_A$R on the postsynaptic side. The AMAR and $GABA_A$R conductances increase instantaneously at each spike and decay exponentially between spikes. The NMDAR is modeled according to Jahr and Stevens (1990, 1991). This is a standard minimal description for the three receptors that has been used previously in the literature (Wang et al., 1999; Wang et al., 2004; Masqualier and Deco, 2013; Lonardoni et al., 2017).

The parameters are listed in **Tables A1-G-I. Table A1-G** gives model parameters that were not fitted to the data but instead selected according to the literature and kept fixed for all simulations. The parameter names, their values and units are given in the table. The last column gives the references used to select the listed parameter values.

**Tables A1-H and A1-I** list the parameters fitted to the experimental data, as described in the paper. The tables give parameter names, the parameter ranges used to select parameter values during model fitting, the physical units, and the selected values used to produce figures in the paper. **Table A1-H** contains the optimal parameters for the five pharmacological conditions obtained using the 'flexible protocol'. These models are discussed in Result Section 2 in the paper. **Table A1-I** shows a model fitted to all pharmacological conditions using the 'constrained protocol'. This model is further analyzed in the Result Section 4 of the paper.

| **A** | **Model summary** |
|---|---|
| **Populations** | Excitatory, inhibitory |
| **Topology** | - |
| **Connectivity** | Random probability of connections ($p$) between each pair of neurons |
| **Neuron model** | Low-dimensional phenomenological model with adaptation and bursting (Izhkevich, 2003; 2004). |
| **Presynaptic** | Short-term depression according to Wang et al. (1999),<br>consistent with short-term depression part in Tsodyks et al. (2000). |
| **Postsynaptic**<br>(syn. receptors) | AMPAR, $GABA_A$R: exponentially decaying conductance (Wang et al. 1999, 2004)<br>NMDAR model: according to Jahr and Stevens (1990, 1991) |
| **External inputs** | 1) White noise<br>2) Short pulses injected into a few randomly selected neurons. |
| Measurements | Spikes, cell membrane potential, ion channel conductances |

| **B** | **Populations** | | |
|---|---|---|---|
| Name | Elements | Size (no. neurons) | Indices: network, connectivity matrix $\mathbf{\Gamma}$ |
| Excitatory (E)<br>Inhibitory (I) | IZH<br>IZH | $N_E = 80$<br>$N_I = N - N_E = 20$ | $i = 1..N_E$<br>indices $i = N_E + 1..N$ |

| **C** | **Connectivity** | |
|---|---|---|
| Source | Target | Pattern |
| $\{E, I\}$ | $\{E, I\}$ | Connectivity matrix $\mathbf{\Gamma} = \{\gamma_{\mathbf{ij}}\}_{\mathbf{i,j=1..N}}$: randomly assigning<br>a connection with probability $p = 0.2$ between each pair of neurons.<br>Binomial distribution of inputs and outputs: $\mathbf{B}(N^2, p)$. |

| **D: Neuron and synapse model** | |
|---|---|
| Type | Integrate-and-fire, postsynaptic channels, presynaptic short-term dynamics |
| Sub-threshold dynamics | |
| GENERAL: | For each neuron $i = 1..N$ in the network |
| 1) Cell membrane | $v_i(t)^{'} = 0.04 \cdot v_i(t)^2 + 5 \cdot v_i(t) + 140 - u_i(t) + I^{syn}(t) + I^{noise}(t) + I^{ext}(t)$ |
| 2) Adaptation | $u_i(t)^{'} = a_i \cdot (b_i \cdot v_i - u_i)$ |
| 3) Spiking and reset | The reset condition: $v_i(t_{sp}^-) \leq V_{th}$ and $v_i(t_{sp}+) > V_{th}$<br>If the reset condition is met then:<br>a) Save the new spike time: $t_{sp} \rightarrow \mathcal{S}$ ($\mathcal{S}$ - the collection of recorded spikes)<br>b) Reset: $v_i(t_{sp}+) \leftarrow c_i, u_i(t_{sp}+) \leftarrow u_i(t_{sp}^-) + d_i$ |
| NEURON MODEL PARAMETERS: | |
| | $\mathcal{P}_i^{neuron} = \{a_i, b_i, c_i, d_i, V_{thr}\}$ |
| Synaptic current | |
| | $I_i^{syn}(t) = \sum_j \sum_X g_{X,ji} \cdot s_{X,ji}(t) \cdot (v_i(t) - E_X), \quad \forall j \in \{1..N\}$ such that $\gamma_{ji} = 1$<br>If $j \leq N_E$ then $X \in \{$AMPAR, NMDAR$\}$<br>If $j > N_E$ then $X =$ GABA$_A$R |
| Presynaptic depression | $y_{Y,ji}(t)^{'} = \frac{1-y_{Y,ji}(t)}{\tau_{D,Y}} - U_{0,ji}^Y \cdot y_{Y,ji}(t_{sp,j}^-) \cdot \delta(t - t_{sp,j}), \quad Y \in \{\text{glu, gaba}\}$ |
| Postsynaptic glutamatergic receptors<br>(excitatory) | $s_{AMPAR,ji}(t)^{'} = -\frac{s_{AMPAR,ji}(t)}{\tau_{AMPAR}} + \sum_j \alpha_{AMPAR} \cdot y_{glu,ji}(t_{sp,j}^-) \cdot (1 - s_{AMPAR,ji}(t_{sp,j})) \cdot \delta(t - t_{sp,j})$<br>$s_{NMDAR,ji}(t) = s_{NMDAR,ji}^*(t) \cdot \left[1 + \frac{[Mg^{2+}]}{3.57} \cdot \exp(-0.062 \cdot \frac{v_i(t)}{mV})\right]^{-1}$<br>$s_{NMDAR,ji}^*(t)^{'} = -\frac{s_{NMDAR,ji}^*(t)}{\tau_{NMDAR,s}} + \alpha_{NMDAR,s} \cdot x_{NMDAR,ji}(t) \cdot (1 - s_{NMDAR,ji}^*(t))$<br>$x_{NMDAR,ji}(t)^{'} = -\frac{x_{NMDAR,ji}(t)}{\tau_{NMDAR,x}} + \sum_j \alpha_{NMDAR,x} \cdot y_{glu,ji}(t_{sp,j}^-) \cdot \delta(t - t_{sp,j})$ |
| GABAergic<br>(inhhibitory) | $s_{GABA_AR,ji}(t) = -\frac{s_{GABA_AR,ji}(t)}{\tau_{GABA_AR}} + \sum_j \alpha_{GABA_AR} \cdot y_{gaba,ji}(t_{sp,j}^-) \cdot (1 - s_{GABA_AR,ji}(t_{sp,j})) \cdot \delta(t - t_{sp,j})$ |
| SYNAPTIC PARAMETERS: | |
| | $\mathcal{P}_{ji}^{syn} = \{g_{AMPAR,ji},\ g_{NMDAR,ji},\ g_{GABA_AR,ji},\ E_{AMPAR},\ E_{NMDAR},\ E_{GABA_AR},$<br>$U_{0,ji}^{glu}, U_{0,ji}^{gaba},\ \tau_{D,glu}, \tau_{D,gaba},\ \tau_{AMPAR},\ \tau_{GABA_AR},$<br>$\tau_{NMDA,s},\ \tau_{NMDA,x},\ \alpha_{AMPAR},\ \alpha_{GABA_AR},\ \alpha_{NMDAR,s},\ \alpha_{NMDAR,x},\ [Mg^{2+}]\}$ |

| **E** | **Inputs** |
|---|---|
| Type | Description |
| Noise | Gaussian noise $I^{noise}(\sigma_{noise})$ injected to each neuron |
| External input | Short pulse $I_i^{ext}(t)$ injected to 10 or 1 randomly selected neuron(s).<br>$I_i^{ext}(t) = A_{inj}$ for $t \leq t_{inj}$; $I^{ext}(t) = 0$ for $t > t_{inj}$. |
| PARAMETERS: | $\mathcal{P}_i^{ext} = \{A_{inj}, t_{inj}\}$, $\mathcal{P}^{noise} = \{\sigma_{noise}\}$ |

| **F Measurements** |
|---|
| 1. Spike raster plots |
| 2. Cell membrane potentials |
| 3. Adaptation variables |
| 4. Dynamics of presynaptic variables |
| 5. Conductances of synaptic receptors |

| **G** | FIXED PARAMETERS | | | |
|---|---|---|---|---|
| Name | Notation | Selected values | Unit | References |
| Spiking threshold | $V_{thr}$ | 30 | mV | Izhkevich (2000) |
| Reversal potential: Glutamate (AMPAR, NMDAR) | $E_{AMPAR}$ $E_{NMDAR}$ | 0 | mV | Latham et al. (2000) Baltz et al. (2011) Masqualier and Deco (2013) Lonardoni et al. (2017) |
| Reversal potential: $\mathrm{GABA_AR}$ | $E_{GABA_AR}$ | -80 | mV | Latham et al. (2000) |
| Time constant: AMPAR | $\tau_{AMPAR}$ | 2 | ms | Gritsun et al. (2010, 2011), Lonardoni et al. (2017) |
| Time constants: NMDAR | $\tau_{NMDAR,s}$ $\tau_{NMDAR,x}$ | 100 2 | ms ms | Wang et al. (1999) Masqualier and Deco (2013) |
| Time constant: $\mathrm{GABA_AR}$ | $\tau_{GABA_AR}$ | 5 | ms | Gritsun et al. (2010) |
| Other AMPAR parameters | $\alpha_{AMPAR}$ | 1 | | |
| Other NMDAR parameters | $\alpha_{NMDAR,s}$ $\alpha_{NMDAR,x}$ | 0.5 1 | kHz | Wang et al. (1999) Masqualier and Deco (2013) |
| Extracellular magnesium concentration | $[Mg^{2+}]$ | 0.69 | nM | Golomb et al. (2010), Markram et al. (2015) |
| Other $\mathrm{GABA_AR}$ paramters | $\alpha_{GABA_AR}$ | 1. | | |
| Gaussian noise (var) | $\sigma_{noise}$ | flex.p.: 1 const.p.: 0 | $\frac{\mathrm{mV}}{\sqrt{\mathrm{s}}}$ | |
| Amplitude of the external pulse current | $A_{inj}$ | flex.p.: 2 const.p.: 2 | $\frac{\mu\mathrm{A}}{cm^2}$ nA | |
| Duration of the pulse current | $t_{inj}$ | flex.p: 10 const.p.: 10 | ms ms | |
| Number of injected (excitatory) neurons | | flex.p.: 1 const.p.: 10 | | |

**Table A1-G:** The parameters listed in this table are kept fixed for all model fitting tasks and all results presented in the paper. The exact values and the references used to select the parameters are indicated. The bottom four rows indicate minor differences in models used in the two model fitting protocols - the flexible protocol (flex.p.) and the constraint protocol (const.p.). Minor differences seen in the table do not affect the presented results (confirmed by running the 'flexible protocol' using both configurations): 1) The noise level was small in both cases and only able to introduce randomness in the cell membrane potential, but it was not able to induce spiking. 2) In both cases, a burst was induced by injecting a short pulse-current into a small subset of excitatory neurons. The amplitude of the current was big enough to drive the injected cells into a short-lived spiking regime (in the absence of recurrent inputs). The spiking quickly propagated across the entire culture.

| **H** | **FITTED PARAMETERS, FLEXIBLE PROTOCOL** | | | | | | | |
|---|---|---|---|---|---|---|---|---|
| Name | Notation | Allowed parameter range while fitting | Unit | **Best parameters per condition:**<br>CTRL | AMPAR blocked | AMPAR, $GABA_AR$ blocked | NMDAR blocked | NMDAR, $GABA_AR$ blocked |
| $a$, mean | $a$ | 0.004, 0.02 | | E: 0.00618<br>I: 0.00708 | E: 0.01071<br>I: 0.00704 | E: 0.01017<br>I: 0.01383 | E: 0.01925<br>I: 0.01779 | E: 0.01902<br>I: 0.01812 |
| $a$, variance | $\sigma_a$ | 0, 0.004 | | E: 0.00196<br>I: 0.00377 | E: 0.00315<br>I: 0.00197 | E: 0.00264<br>I: 0.00065 | E: 0.00048<br>I: 0.00053 | E: 0.00317<br>I: 0.00252 |
| $b$, mean | $b$ | 0.254, 0.2543 | | E: 0.25412<br>I: 0.25424 | E: 0.25409<br>I: 0.25417 | E: 0.25413<br>I: 0.25429 | E: 0.25419<br>I: 0.2542 | E: 0.2542<br>I: 0.25417 |
| $b$, variance | $\sigma_b$ | 0, 0.0001 | | E: $6.904 \cdot 10^{-6}$<br>I: $8.866 \cdot 10^{-5}$ | E: $1.027 \cdot 10^{-5}$<br>I: $9.137 \cdot 10^{-6}$ | E: 0.0001<br>I: 0 | E: $1.841 \cdot 10^{-5}$<br>I: $9.1302 \cdot 10^{-7}$ | E: $1.787 \cdot 10^{-6}$<br>I: $7.008 \cdot 10^{-6}$ |
| $c$, mean | $c$ | -65.0, -46.0 | mV | E: -52.887<br>I: -48.015 | E: -56.602<br>I: -48.207 | E: -64.38<br>I: -56.197 | E: -61.053<br>I: -52.109 | E: -52.274<br>I: -55.339 |
| $c$, variance | $\sigma_c$ | 0, 5 | mV | E: 0.09589<br>I: 1.55394 | E: 3.00054<br>I: 3.17847 | E: 1.42391<br>I: 1.77868 | E: 2.57519<br>I: 2.77453 | E: 0.70889<br>I: 2.34065 |
| $d$, mean | $d$ | 1.39, 2 | $\frac{V}{s}$ | E: 1.9835<br>I: 1.52545 | E: 1.78568<br>I: 1.97387 | E: 1.93706<br>I: 1.8306 | E: 1.46915<br>I: 1.88684 | E: 1.58352<br>I: 1.70825 |
| $d$, variance | $\sigma_d$ | 0, 0.15 | $\frac{V}{s}$ | E: 0.08938<br>I: 0.00877 | E: 0.06311<br>I: 0.15 | E: 0<br>I: 0.04662 | E: 0.10917<br>I: 0.06101 | E: 0.06905<br>I: 0.05797 |
| Conductance, AMPAR | $g_{AMPAR,ji}$ | 0, 1000 | $\frac{\mu S}{cm^2}$ | E: 267.301<br>I: 105.535 | E: 0<br>I: 150.248 | E: 0<br>I: 0 | E: 28.112<br>I: 0 | E: 587.378<br>I: 105.535 |
| Conductance, NMDAR | $g_{NMDAR,ji}$ | 0, 1000 | $\frac{\mu S}{cm^2}$ | E: 2.569<br>I: 30.294 | E: 141.8<br>I: 305.718 | E: 56.482<br>I: 158.459 | E: 0<br>I: 0 | E: 0<br>I: 0 |
| Conductance $GABA_AR$ | $g_{GABA_AR,ji}$ | 0, 1000 | $\frac{\mu S}{cm^2}$ | E: 680.512<br>I: 405.833 | E: 209.891<br>I: 852.976 | E: 0<br>I: 0 | E: 135.533<br>I: 541.927 | E: 0<br>I: 0 |
| Time constant | $\tau_{D,glu/gaba}$ | 0.1, 6.4 | s | EE: 2.883<br>EI: 5.325<br>IE: 0.227<br>II: 2.543 | EE: 2.692<br>EI: 5.207<br>IE: 0.1<br>II: 3.182 | EE: 1.272<br>EI: 5.979<br>IE: 1.37<br>II: 6.161 | EE: 2.411<br>EI: 6.054<br>IE: 2.537<br>II: 3.746 | EE: 1.992<br>EI: 5.65<br>IE: 5.867<br>II: 4.377 |
| Presynaptic resources | $U_{0,ji}^{glu/gaba}$ | 0, 1 | | EE: 0.927<br>EI: 0.2902<br>IE: 0.5419<br>II: 0.1897 | EE: 1<br>EI: 0.1487<br>IE: 0.1487<br>II: 0.4968 | EE: 0.44<br>EI: 0.9074<br>IE: 0.6304<br>II: 0.6662 | EE: 0.922<br>EI: 0.1237<br>IE: 0.7643<br>II: 0.242 | EE: 0.01<br>EI: 0.592<br>IE: 0.5596<br>II: 0.9518 |

**Table A1-H** Optimal parameters per pharmacological condition obtained using the 'flexible' protocol. These models are analyzed in Result Section 2, Fig 4 in the paper. The parameters are rounded to fit to the table format. E - excitatory neurons, I - inhibitory neurons, EE - a synapse from excitatory to excitatory neuron, EI - a synapse from excitatory to inhibitory neuron, IE - a synapse from inhibitory to excitatory neuron, II - a synapse between a pair of inhibitory neurons. These same parameters represented with a higher precision are supplied together with the data and the code used to prepare this paper.

| I | FITTED PARAMETERS, CONSTRAINED PROTOCOL | | | |
|---|---|---|---|---|
| Name | Notation | Allowed range of parameters during fitting | Unit | **Best parameters** |
| $a$, mean | $a$ | 0.004, 0.02 | | E: 0.00884<br>I: 0.00672 |
| $a$, variance | $\sigma_a$ | 0, 0.004 | | E: 0.00001<br>I:0.00008 |
| $b$, mean | $b$ | 0.254, 0.2543 | | E: 0.25428<br>I: 0.25411 |
| $b$, variance | $\sigma_b$ | 0, 0.0001 | | E: 0.00006<br>I: 0.00003 |
| $c$, mean | $c$ | -65.0, -46.0 | mV | E: -60.64335<br>I: -54.17188 |
| $c$, variance | $\sigma_c$ | 0, 5 | mV | E: 1.01475<br>I: 0 |
| $d$, mean | $d$ | 1.39, 2 | $\frac{\text{V}}{\text{s}}$ | E: 1.87622<br>I: 1.9206 |
| $d$, variance | $\sigma_d$ | 0, 0.15 | $\frac{\text{V}}{\text{s}}$ | E: 0.00241<br>I: 0.14878 |
| Conductance, AMPAR | $g_{AMPAR,ji}$ | 0, 1000 | $\frac{\mu\text{S}}{\text{cm}^2}$ | E: 0.19842<br>I: 0.02479 |
| Conductance, NMDAR | $g_{NMDAR,ji}$ | 0, 1000 | $\frac{\mu\text{S}}{\text{cm}^2}$ | E: 0.04295<br>I: 0.06145 |
| Conductance, $\text{GABA}_\text{A}\text{R}$ | $g_{GABA_AR,ji}$ | 0, 1000 | $\frac{\mu\text{S}}{\text{cm}^2}$ | E: 0.73249<br>I: 0.04912 |
| Time constant | $\tau_{D,glu/gaba}$ | 0.1, 6.4 | s | All synapses: 2.62658 |
| Presynaptic resources | $U_{0,ji}^{glu/gaba}$ | 0, 1.0 | | All synapses: 0.10536 |

**Table A1-I** Optimal parameters resulting from the 'constrained' protocol. Models for different pharmacological conditions are obtained by setting the appropriate conductances to zero and using the rest of the parameters from this table. The properties of this model are analyzed in Result Section 4, Fig. 6. E - excitatory neuron, I - inhibitory neuron. All four types of synapses (EE, EI, IE, II) are assigned the same values for the presynaptic parameters ($\tau_D$, $U_0$).
