## Appendix 2, objective functions for "Analysis of cellular and synaptic mechanisms behind spontaneous cortical activity *in vitro*: Insights from optimization of spiking neuronal network models"

### Appendix 2: Burst measures, distance measures and objective functions used in model fitting.

| A: Network burst measures | | | |
|---|---|---|---|
| **Name** | Acronym | Unit | Description |
| **Burst length** | BL | second | Time between the first and the last spike in a network burst |
| **Burst size** | BS | #spikes/#active electrodes | Number of spikes within a burst divided by the number of active electrodes for *in vivo* data. Number of spikes within a burst divided by or the number of neurons in a network for *in silico* data. *In silico* , typically, all the neurons get activated during a burst. |

**Table A2-A:** Summary of burst measures extracted from the experimental *in vitro* data and from the *in silico* data.

| B: Distance measures | | |
|---|---|---|
| **Name** | Equation | Description |
| **Network burst measure** | $B \in \{BL, BS\}$ | See Table S2-A |
| **Experimental (*in vitro* ) data set** | $\{B_l^{exp}\}_{l=1..N_{exp}}$ | All estimations from the experimental data. |
| **Simulated (*in silico* ) data set** | $\{B_l^{sim}\}_{l=1..N_{sim}}$ | $N_{sim}$ estimations obtained from computational model simulations. |
| **Mean value** | $\overline{B} = \frac{1}{N}\sum_{l=1..N} B_l$ where $N \in \{N_{exp}, N_{sim}\}$ | Mean value computed from all data. |
| **Histogram of a data set** | $\mathcal{H}(B) = \text{hist}\left(\{B_l\}_{l=1..N}\right)$ $N \in \{N_{exp}, N_{sim}\}$ | Histogram, approximates distribution of a burst measure. |
| **Distance between means** | $D^m = \lvert\overline{B}^{exp} - \overline{B}^{sim}\rvert$ | Absolute value between means of the experimental and the simulated data sets. |
| **Distance between distributions** | $D^d = JSD\left(\mathcal{H}^{exp}(B), \mathcal{H}^{sim}(B)\right)$ $= \frac{1}{2} \cdot KL\left(\mathcal{H}^{exp}(B) \Vert \mathcal{H}^m(B)\right)$ $+\frac{1}{2} \cdot KL\left(\mathcal{H}^{sim}(B) \Vert \mathcal{H}^m(B)\right)$ | JSD - Jenssen-Shannon divergence between two histograms. |
| | $KL\left(\mathcal{H}, \mathcal{H}^m\right) = -\sum_{bins} \mathcal{H} \log\left(\frac{\mathcal{H}}{\mathcal{H}^m}\right)$ | KL - Kullback-Leibler divergence. |
| | $\mathcal{H}^m = \frac{1}{2}(\mathcal{H}^{exp} + \mathcal{H}^{sim})$ | $\mathcal{H}^m$ - mean between two histograms is computed as the mean in each of the histogram bins. |

**Table A2-B:** Two distance measures (distance between means and distance between distributions) used to compute the objective functions, together with the variables and functions used to define the measures.

| C: Objective functions |
|---|
| **Flexible protocol:** $Obj_1 = JSD\left(\mathcal{H}^{sim}(BL_X), \mathcal{H}^{exp}(BL_X)\right),$ $Obj_2 = JSD\left(\mathcal{H}^{sim}(BS_X), \mathcal{H}^{exp}(BS_X)\right)$ $X \in \{CTRL, AMPAR^{block}, AMPAR^{block}\&GABA_{\mathrm{A}}R^{block}, NMDAR^{block}, NMDAR^{block}\&GABA_{\mathrm{A}}R^{block}\}$ |
| **Constrained protocol:** $Obj_1 = \left\|\overline{BL}^{sim}_{NMDAR^{block}} - \overline{BL}^{exp}_{NMDAR^{block}}\right\| + \left\|\overline{BL}^{sim}_{NMDAR^{block}\&GABA_{\mathrm{A}}R^{block}} - \overline{BL}^{exp}_{NMDAR^{block}\&GABA_{\mathrm{A}}R^{block}}\right\|$ $Obj_2 = \left\|\overline{BL}^{sim}_{AMPAR^{block}} - \overline{BL}^{exp}_{AMPAR^{block}}\right\| + \left\|\overline{BL}^{sim}_{AMPAR^{block}\&GABA_{\mathrm{A}}R^{block}} - \overline{BL}^{exp}_{AMPAR^{block}\&GABA_{\mathrm{A}}R^{block}}\right\|$ $Obj_3 = \left\|\overline{BL}^{sim}_{CTRL} - \overline{BL}^{exp}_{CTRL}\right\|$ $Obj_4 = \left\|\frac{\overline{BS}^{sim}_{NMDAR^{block}}}{\overline{BS}^{sim}_{CTRL}} - \frac{\overline{BS}^{exp}_{NMDAR^{block}}}{\overline{BS}^{exp}_{CTRL}}\right\| + \left\|\frac{\overline{BS}^{sim}_{NMDAR^{block}\&GABA_{\mathrm{A}}R^{block}}}{\overline{BS}^{sim}_{CTRL}} - \frac{\overline{BS}^{exp}_{NMDAR^{block}\&GABA_{\mathrm{A}}R^{block}}}{\overline{BS}^{exp}_{CTRL}}\right\|$ $Obj_5 = \left\|\frac{\overline{BS}^{sim}_{AMPAR^{block}}}{\overline{BS}^{sim}_{CTRL}} - \frac{\overline{BS}^{exp}_{AMPAR^{block}}}{\overline{BS}^{exp}_{CTRL}}\right\| + \left\|\frac{\overline{BS}^{sim}_{AMPAR^{block}\&GABA_{\mathrm{A}}R^{block}}}{\overline{BS}^{sim}_{CTRL}} - \frac{\overline{BS}^{exp}_{AMPAR^{block}\&GABA_{\mathrm{A}}R^{block}}}{\overline{BS}^{exp}_{CTRL}}\right\|$ <br><br> Mean values extracted from the experimental data: <br> $\overline{BL}^{exp}_{CTRL} = 0.69172s$ <br> $\overline{BL}^{exp}_{AMPAR^{block}} = 3.46233s$ $\frac{\overline{BS}^{exp}_{AMPAR^{block}}}{\overline{BS}^{exp}_{CTRL}} = 4.116$ <br> $\overline{BL}^{exp}_{AMPAR^{block}\&GABA_{\mathrm{A}}R^{block}} = 0.93481s$ $\frac{\overline{BS}^{exp}_{AMPAR^{block}\&GABA_{\mathrm{A}}R^{block}}}{\overline{BS}^{exp}_{CTRL}} = 1.71$ <br> $\overline{BL}^{exp}_{NMDAR^{block}} = 0.17s$ $\frac{\overline{BS}^{exp}_{NMDAR^{block}}}{\overline{BS}^{exp}_{CTRL}} = 0.269$ <br> $\overline{BL}^{exp}_{NMDAR^{block}\&GABA_{\mathrm{A}}R^{block}} = 0.39423s$ $\frac{\overline{BL}^{exp}_{NMDAR^{block}\&GABA_{\mathrm{A}}R^{block}}}{\overline{BS}^{exp}_{CTRL}} = 1.979$ |

**Table A2-C:** Two objective functions used in the 'flexible protocol', and five objectives used in the 'constrained protocol'. The mean burst measures extracted from the experimental data are given together with the objectives of the 'constrained protocol'. The histograms of the experimentally obtained burst measures used in the 'flexible protocol' are shown in Fig2 in the paper.
