## Supplementary data 3, flexible protocol for "Analysis of cellular and synaptic mechanisms behind spontaneous cortical activity *in vitro*: Insights from optimization of spiking neuronal network models"

### S3: Supporting data for the ’flexible protocol’.

The supplementary document presents additional data discussed in Result sections 2 and 3. **Fig S3-1** illustrates convergence of the flexible protocol in the test phase, the training phase is shown in Fig3 of the paper. **Fig S3-2** contains three control simulations used to discuss fitting of NMDAR and $GABA_AR$ blocked condition. **Fig S3-3** extends one of the controls, namely it shows results of model fitting using the ’flexible protocol’ when no synaptic blocking is done *in silico*. **Fig S3-4** present the results of sensitivity analysis for all but the BLs in control. **Fig S3-5** shows the dynamics of model variables in case of the best model obtained for each condition. This figure is used to illustrate model behavior and the deviation observed through sensitivity analysis. The figures **Fig S3-6-11** presents the convergence of the ’flexible protocol’ in training and test phase, as well as the properties of the optimal model when a) parameter variability is allowed in synaptic parameters, b) no parameter variability is allowed and thus all neurons or synapses of the same type are identical. The last figure **Fig S3-12** presents intrinsic dynamics of randomly selected neurons in the optimal model when excited with a pulse current.

#### Convergence of the ’flexible protocol’ in the test phase

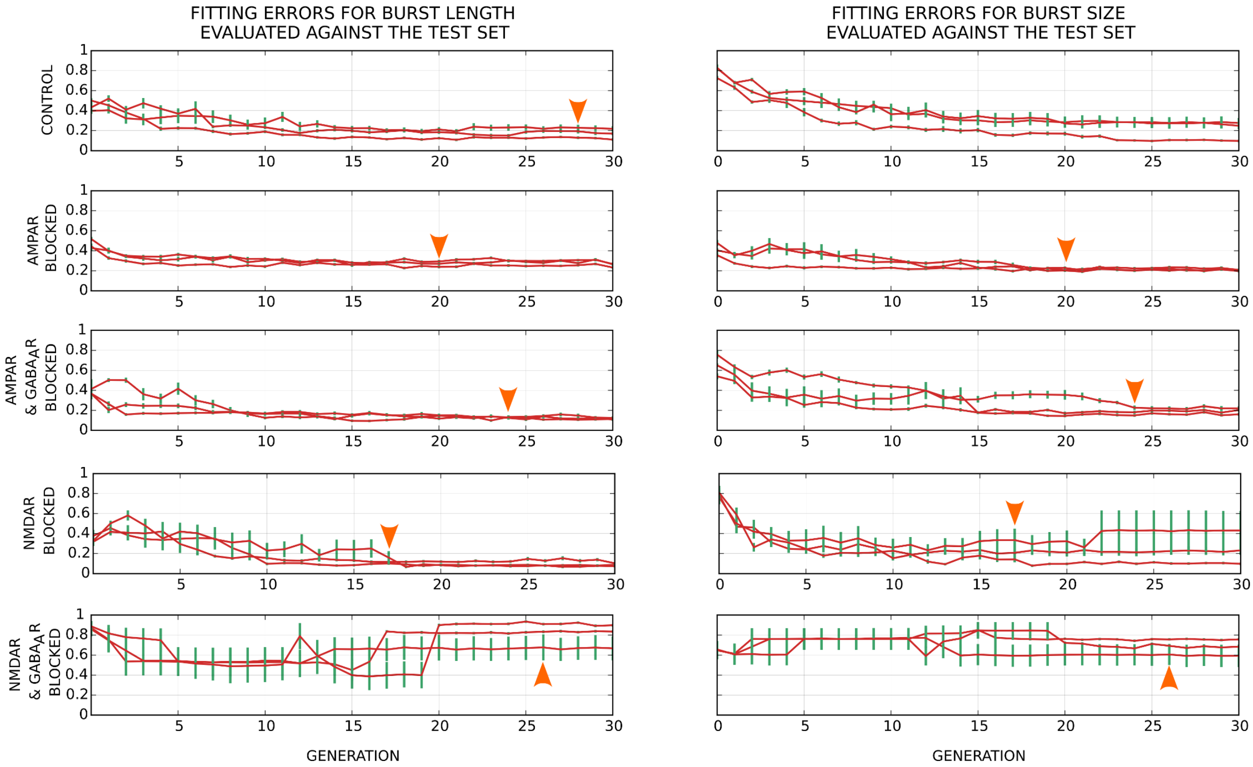


Fig S3-1: This figure complements the results shown in Fig3 (main part of the paper). Fig3 illustrates convergence of the ’flexible algorithm’ during model fitting, i.e. in the training phase. Here, for each generation, the best eight models were selected as described in the paper. The BL and BS distributions are compared to the test data (30% of the experimental data not used during model fitting). Three lines correspond to the three model fitting trials, each initiated using a different random seed. The solid lines are the mean fitting errors, while the whiskers around them correspond to [mean ± variance]. The optimal generations, selected during the training phase, are shown by arrows. Unlike the Fig3, the errors shown here are not always monotonously decreasing with generations, in some fitting trials they increase at later generations (e.g. see one fitting trial in NMDAR blocked condition) suggesting overfitting. However, the optimal selected generations ensure convergence in at least some of the fitting trials.

### Additional model fittings for NMDAR & $\text{GABA}_\text{A}$R blocked condition

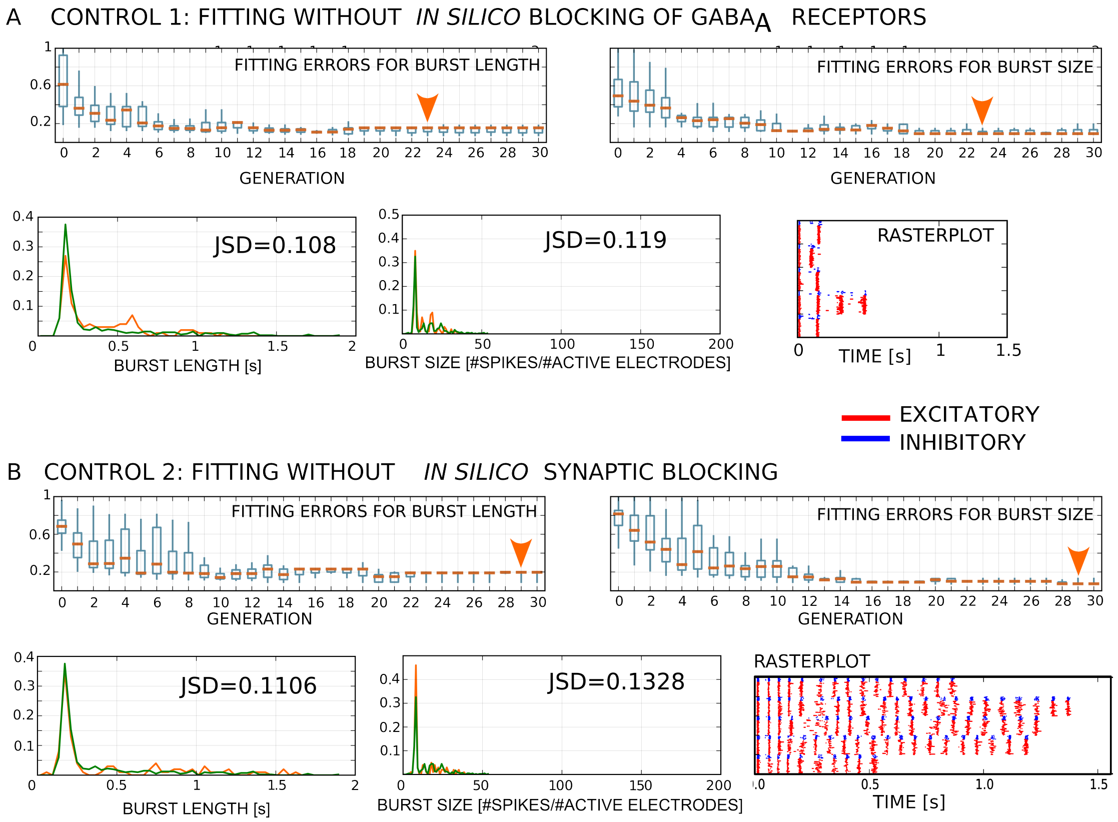


**Fig S3-2:** Control model fittings for NMDAR & $\text{GABA}_\text{A}$R condition. **A:** In Control 1, we assumed an incomplete blocking of $\text{GABA}_\text{A}$ receptors, and fitted the model with active AMPARs and $\text{GABA}_\text{A}$Rs and blocked NMDARs. **B:** In the Control 2, we fitted the entire model without *in silico* blocking to the experimental data obtained from NMDAR & $\text{GABA}_\text{A}$R blocked pharmacological condition. The best generation and the best model were selected following the same steps used to obtain Fig3 in the paper. The best model provided good fit to the experimental data for both BLs and BSs with the distance between experimental and simulated distributions being JSD=0.1106 (for BLs) and JSD=0.1328 (for BSs).

### Fitting the full model to individual experimental conditions

Here, we give additional examples obtained when fitting the full model, with all synaptic receptor being active, to the experimental data. These tests are used as a proof of concept that every experimental data set can be fitted using our approach. Furthermore, we show that every model fitting trial succeed even for a relatively small number of generations (20).

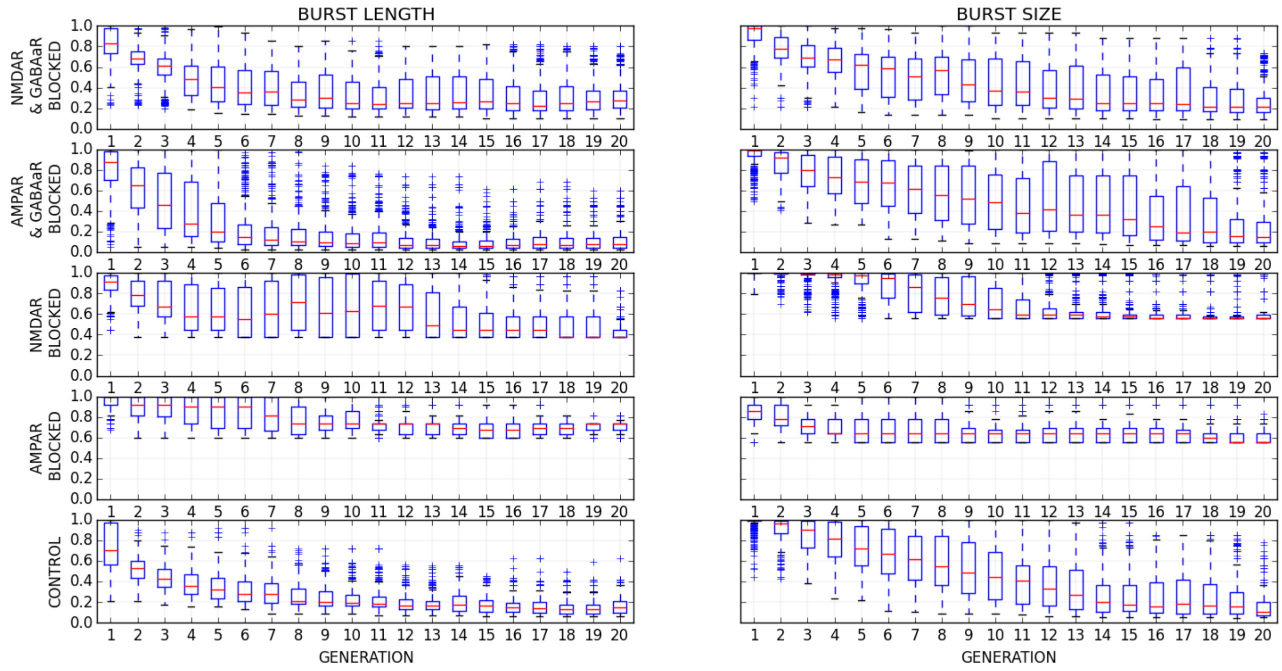


**Fig S3-3:**Convergence of the model fitting algorithm over the course of 20 generations.

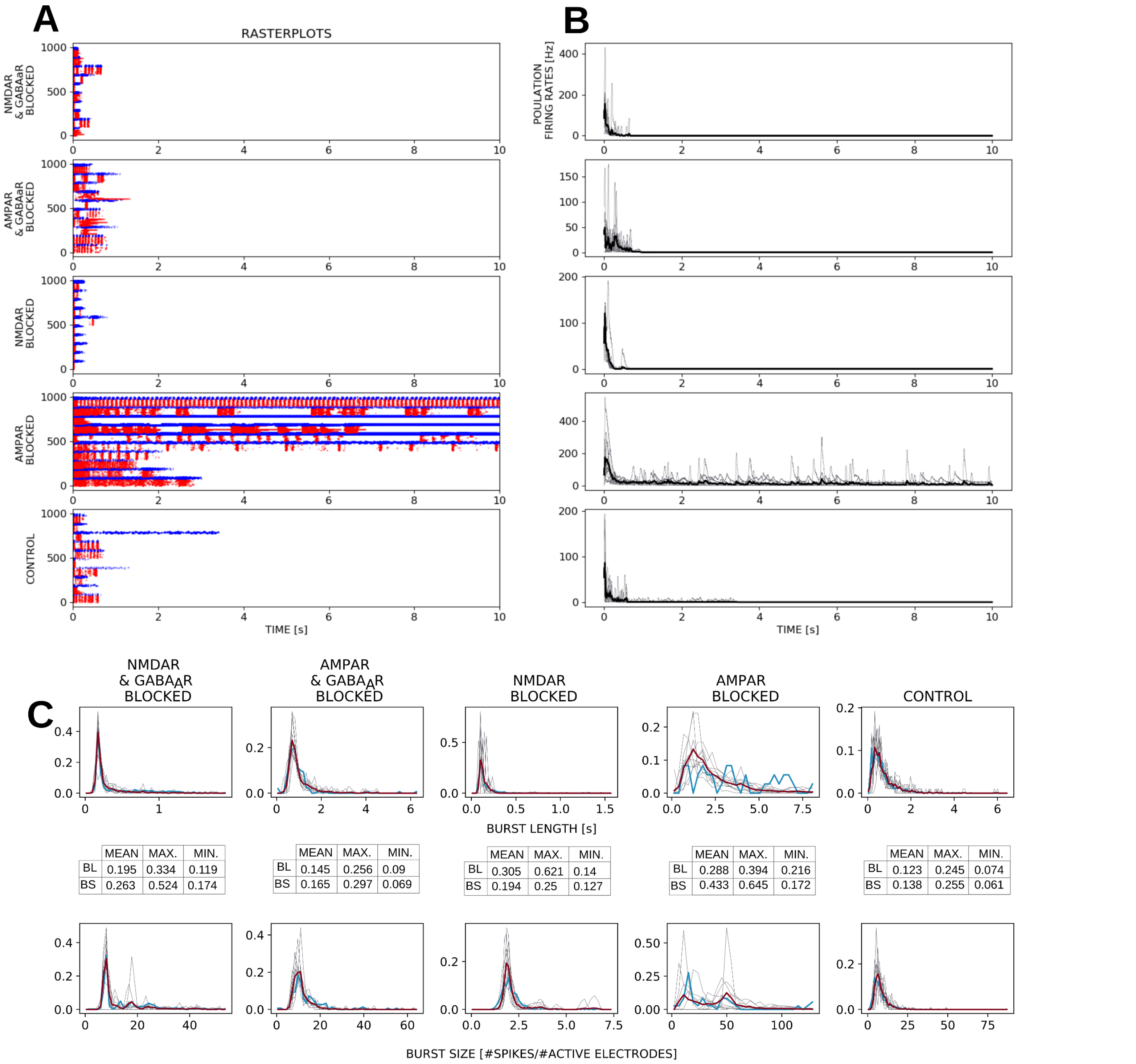


**Fig S3-4:** Analysis of the best models obtained when fitting the full model to each pharmacological condition. In order to show how often model fitting succeeds or fails we present each model fitting trial separately, contrary to Fig 4 when we present the best model obtained from all the trials. **A:** Each rasterplot is obtained as the best result from one model fitting trial (red-excitatory neurons, blue-inhibitory neurons). 10 rasterplots from 10 model fitting trials are plotted together. For the control, AMPAR blocked, NMDAR blocked and AMPAR & GABA$_A$R blocked conditions each fitting trial provides a good fitting result. In case of AMPAR & GABA$_A$R blocked condition, some of the fitting trials result in ceaseless bursts, while others provide good results. **B:** The population rates computed from the rasterplots shown to the left. **C:** BL and BS distribution for the best model obtained from each fitting trial (grey lines), distribution pooled across 10 models (red) and the distribution of the experimental data (blue). The tables in the middle show the minimal, maximal and mean JSD obtained across 10 trials.

### Additional examples of sensitivity analysis, extension of Fig 4N-R

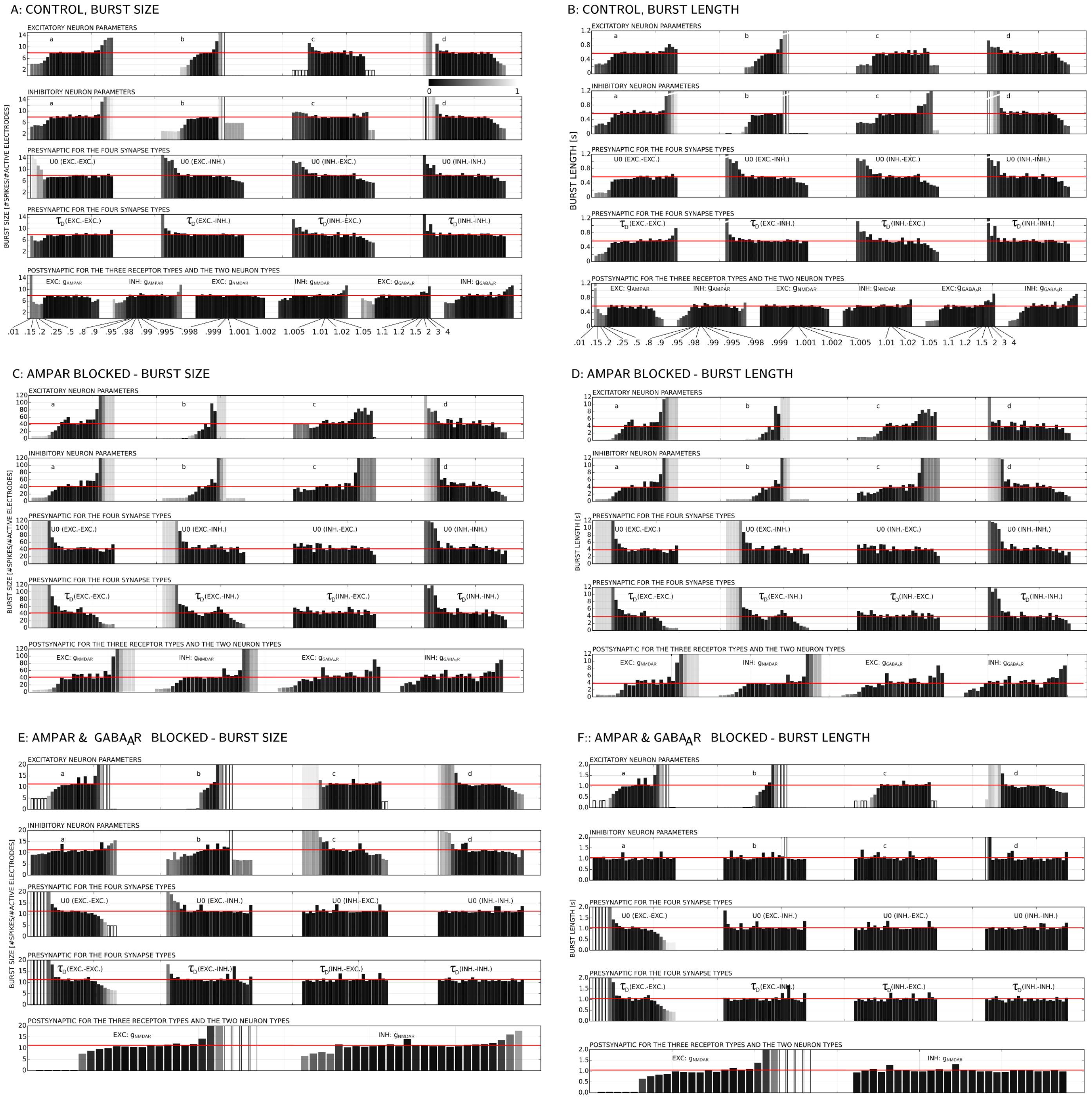

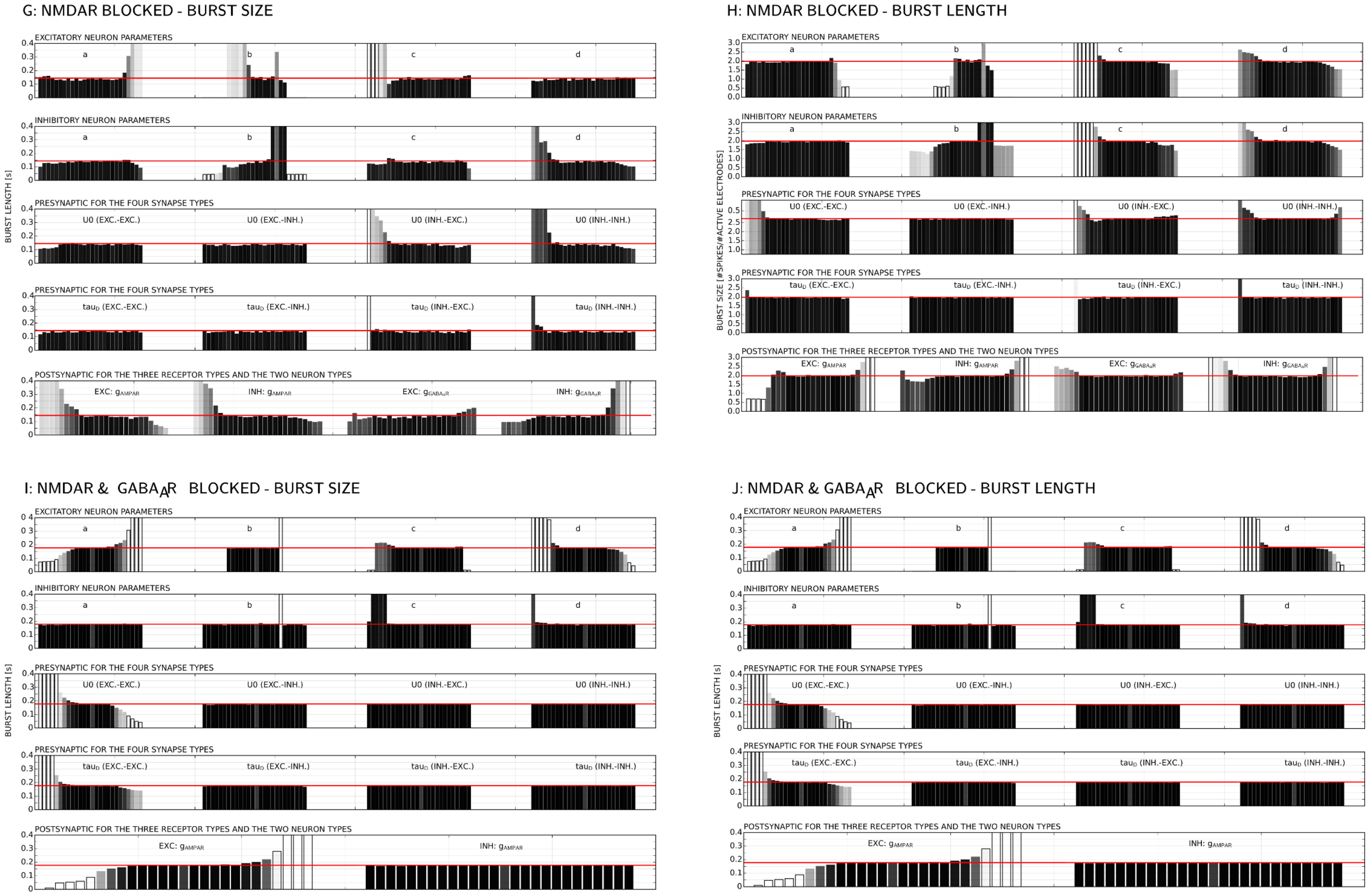


**Fig S3-5:** Sensitivity analysis for flexible protocol. Figure shows the changes in BLs and BSs when varying one parameter at a time. The varied parameter, indicated on top of the corresponding result, is multiplied by a coefficient between 0.01 and 4 (see panel A). The model simulated 100 times to obtain BL and BS distribution. The average BL and BS are shown as bars. The BL and BS of the original model are also shown as bars and additionally indicated by the red horizontal lines. The color of the bar quantifies the distance from the original model (JSD, 0- perfect overlap of distributions, 1-complete mismatch; see the colorbar in the upper right corner of Fig S3-5A). The summary of these results is shown in Fig 4O in the main paper. **A,B:** Control condition (the panel B is repeating Fig 4N-R for easier comparison across pharmacological conditions). **C,D:** AMPAR blocked condition. **E, F:** AMPAR & $GABA_A$R blocked condition. **G, H:** NMDAR blocked condition. **I, J:** NMDAR & $GABA_A$R blocked condition.

#### Supporting data for the analysis of parameter variability

In Fig4 we show a spiking network model in which neuronal excitability (defined by Izhkevich model) is randomized across neuronal population, i.e. the four Izhkevich model parameters are generated from Gaussian distribution (mean and variance fitted to the data) for each neuron. In addition, we fitted the following two models: a) the model where neuronal parameters were the same for all neurons but the synaptic parameters differ across synapses of the same type; b) the model with no randomness in neuronal or synaptic parameters.

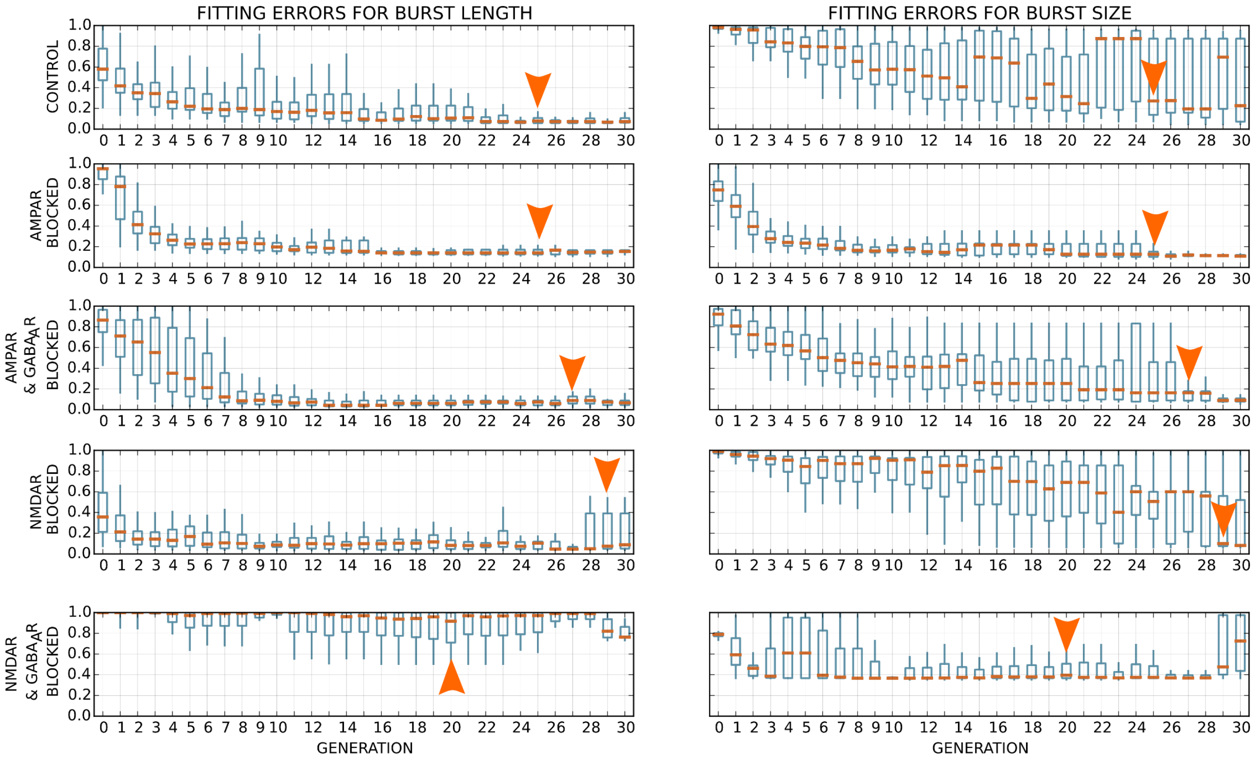


**Fig S3-6: Convergence of the flexible protocol when no variability is allowed between the neurons of the same type (thus, the model is composed of two homogeneous populations of neurons), but the presynaptic and the postsynaptic parameters are allowed to vary. The best generation is selected as before.**

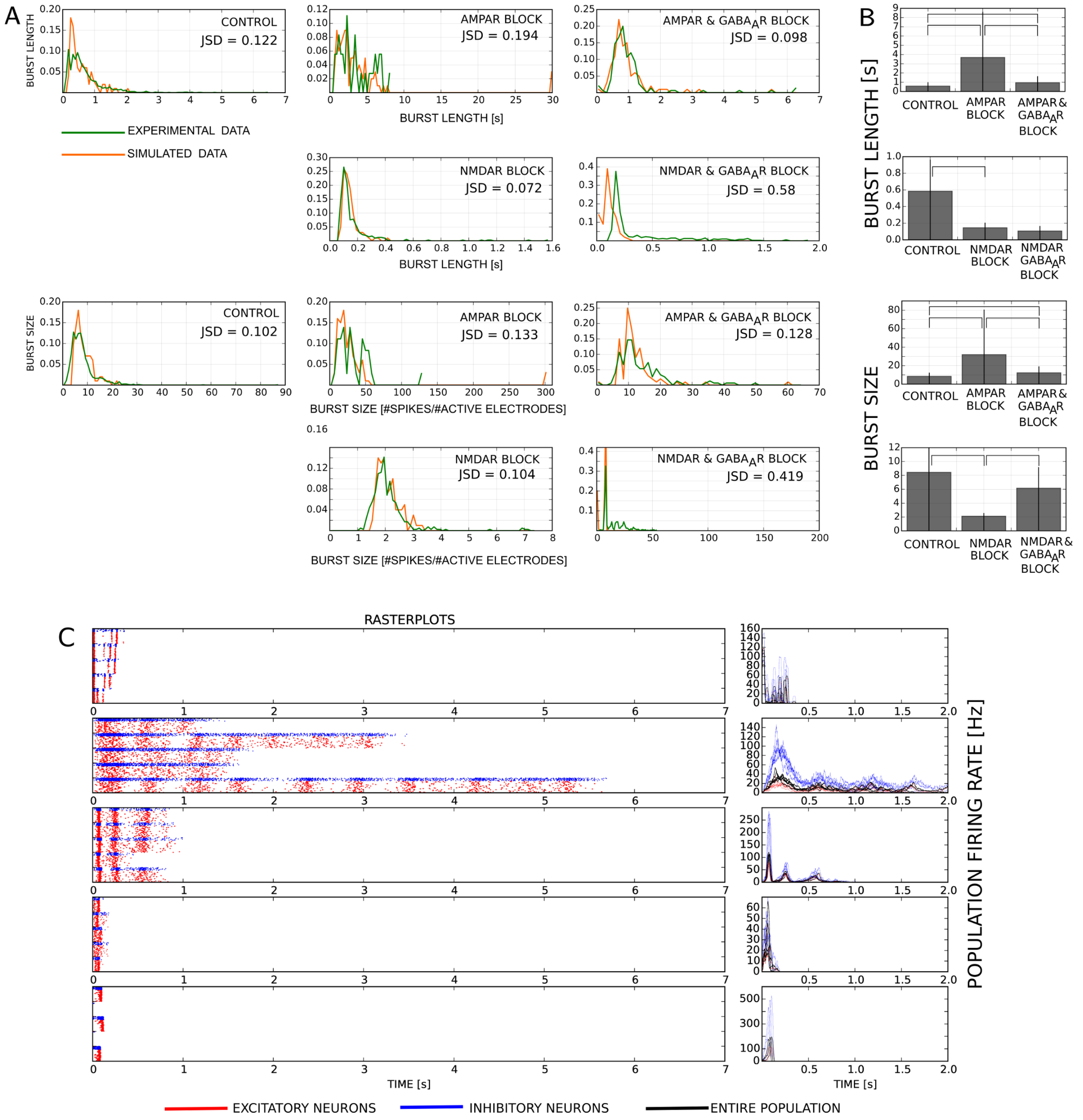


**Fig S3-7:** Best model obtained when no variability is allowed between the neurons of the same type but the presynaptic and the postsynaptic parameters are allowed to vary. **A:** Similarity between BL and BS distributions obtained by simulating the best model (orange) and from the experimental data in Fig 2 (green). JSD between pairs of distributions is shown in each panel. B: Comparison of BL and BS distributions across pharmacological conditions. Bars represent the average values, and whiskers mark [mean ± variance]. The burst measures are compared across pharmacological conditions using 1-sided Kolmogorov-Smirnov test at significance level $p < 0.01$. The angular brackets mark tests that rejected H0 at the given significance level. This model provided a better fit to the experimentally measured BS in NMDAR and GABA$_A$R blocked condition, compared to other models presented in this paper. However the obtained values were still significantly lower than the experimental data, and thus this model failed to reproduce the experimental findings in Fig 2E-H. **C:** Rasterplots and population rates obtained by 5 simulations of the optimal model (connectivity and neuron model parameters randomized). Red - excitatory neurons, blue - inhibitory neurons, black - the entire population.

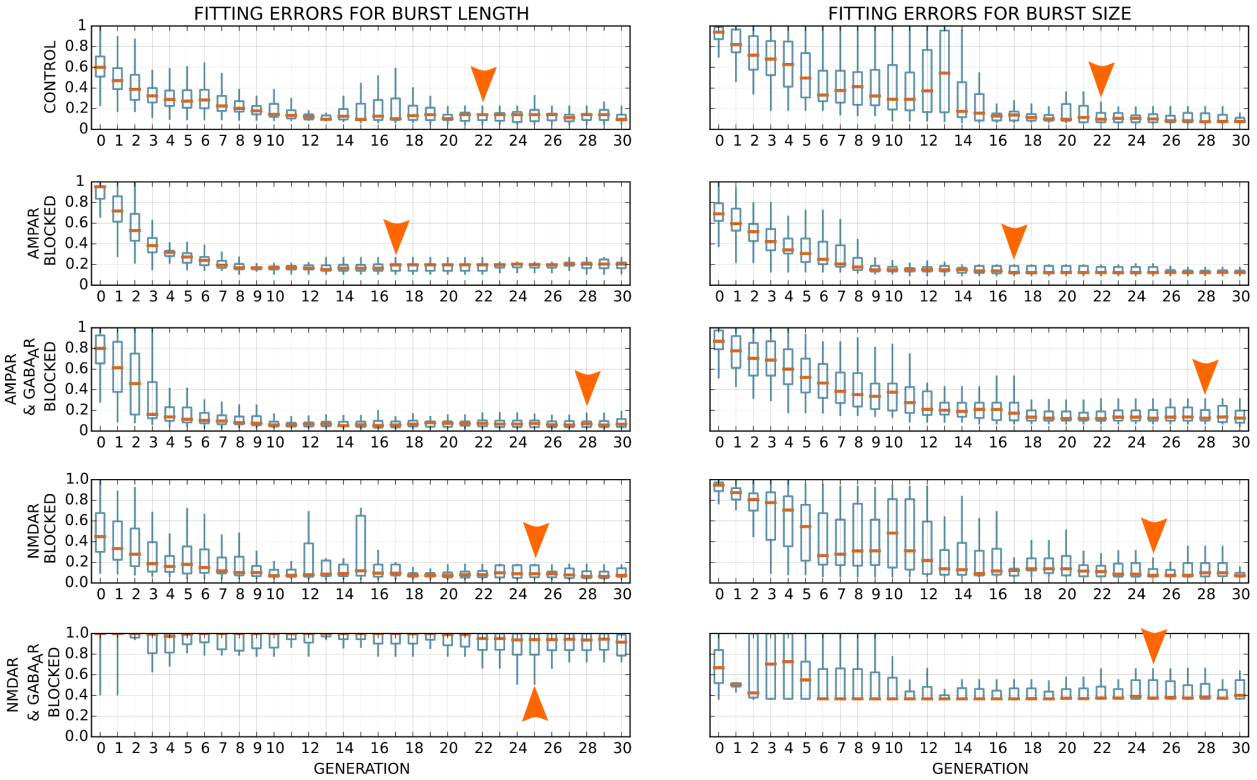


**Fig S3-8:** Convergence of the flexible protocol when no variability is allowed between the neurons of the same type nor between the synapses of the same type. The best generation is selected as before.

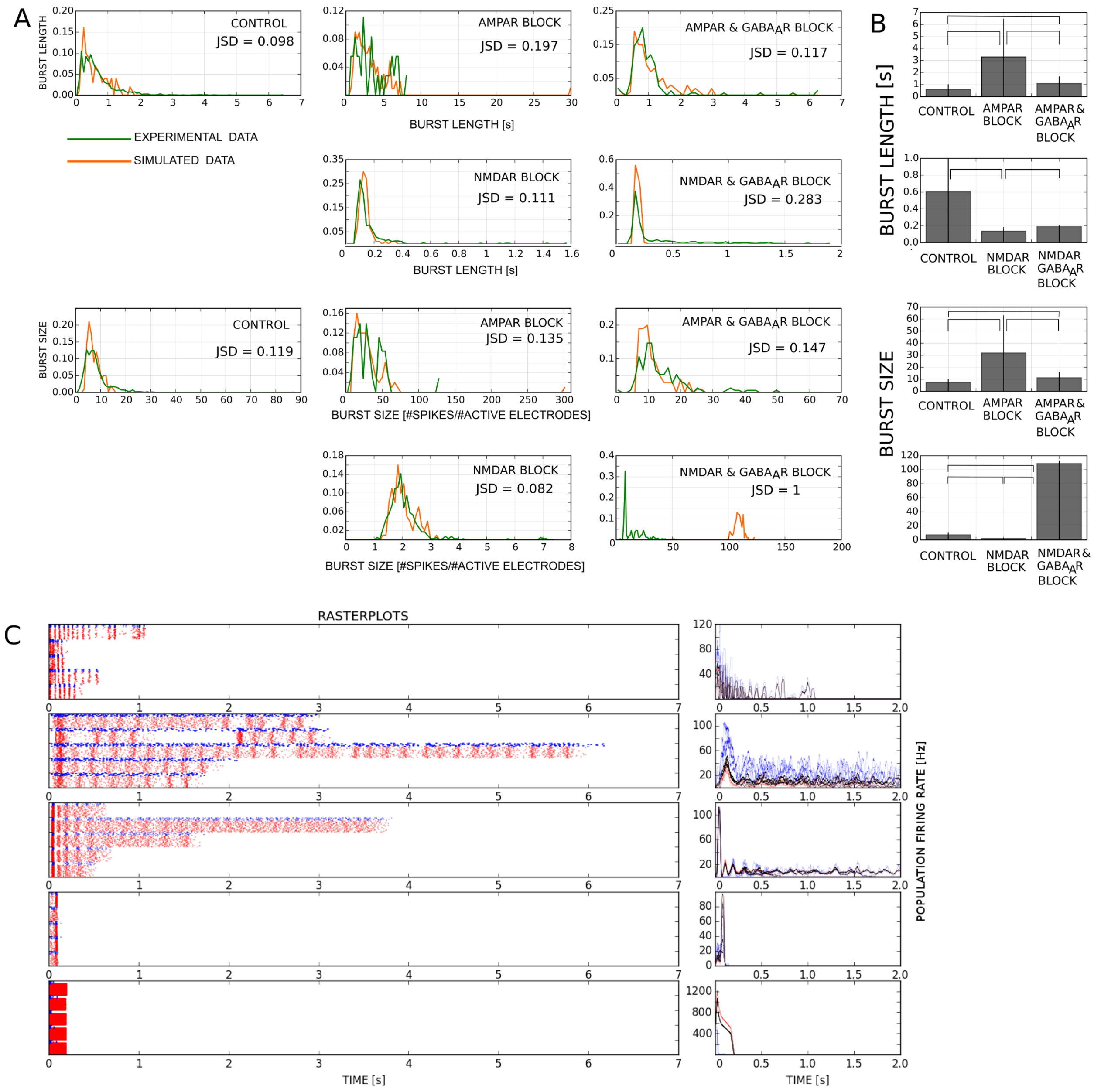


**Fig S3-9:** Best model obtained when no variability is allowed between the neurons of the same type nor between the synapses of the same type. **A:** Similarity between BL and BS distribution for the best model (orange) and the experimental data (green); JSD between distribution shown in each panel. **B:** Comparison of BL and BS distributions across pharmacological conditions. Bars represent the average values, and whiskers mark [mean ± variance]. The burst measures are compared across pharmacological conditions using 1-sided Kolmogorov-Smirnov test at significance level $p < 0.01$. The angular brackets mark tests that rejected H0 at the given significance level. The same significant increases and decreases of BLs and BSs across pharmacological conditions were found in the experimental data in Fig 2. **C:** Rasterplots and population rates obtained by 5 simulations of the optimal model (connectivity and neuron model parameters randomized). Red - excitatory neurons, blue - inhibitory neurons, black - the entire population.

### Spiking patterns of optimized neurons

We examined the spiking patters of excitatory and inhibitory neuron models obtained using flexible protocol. Upon selecting the optimal model per condition we randomly selected five excitatory and five inhibitory neurons and stimulated them for 900ms with a pulse current of $2\frac{\mu A}{cm^2}$. Supplementary Fig A3-10 shows the cell membrane potentials (in mV) obtained for one model fitting trial where we allowed variability of neuronal model parameters. In most of the conditions, the preference for tonic spiking or bursting was visible across a neuronal type (excitatory or inhibitory) but some exceptions also existed (e.g. both tonic spiking and bursting neurons appeared among the inhibitory neurons of NMDAR blocked condition). The preference for specific spiking patterns across experimental conditions was not evident. Furthermore, different model fitting trials resulted in different spiking patterns across neuronal types and experimental conditions (data not shown). The same conclusion was obtained when we excluded variability in neuronal model parameters. Clearly, the considered model fitting task, where one spiking network model is trained to reproduce one experimental condition, permits substantial flexibility when selecting the optimal solution which results in variable optimal models which do not exhibit visible similarity in their dynamical regimes.

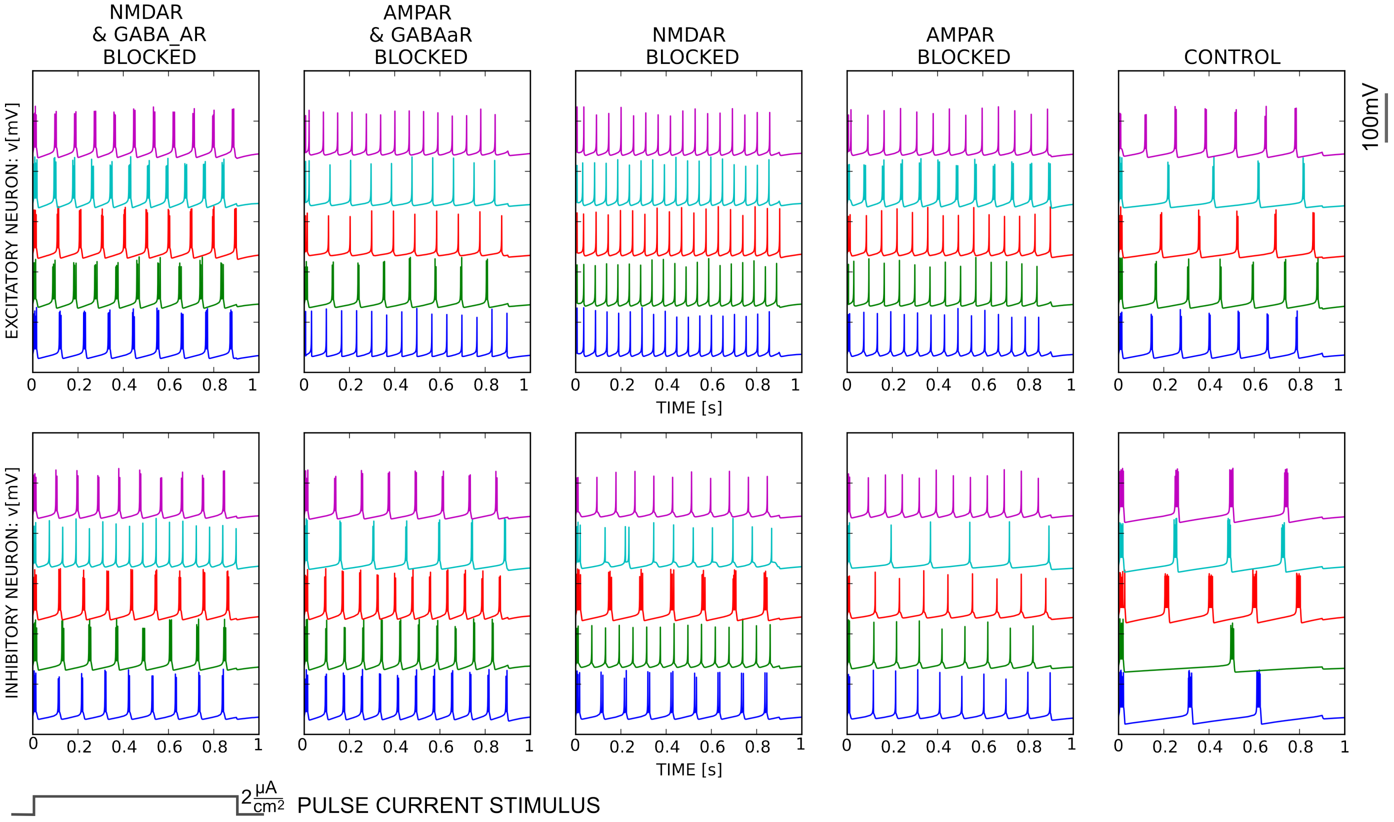


**Fig S3-10** illustrates the spiking patterns of excitatory and inhibitory neurons in models fitted individual experimental conditions. Top panels: five randomly selected excitatory neurons; bottom panels: five randomly selected inhibitory neurons. The experimental conditions are indicated above the two corresponding panels. The voltage traces (in mV) are obtained by stimulating isolated neurons with a pulse current of $2\frac{\mu A}{cm^2}$.

### Dynamics of model variables

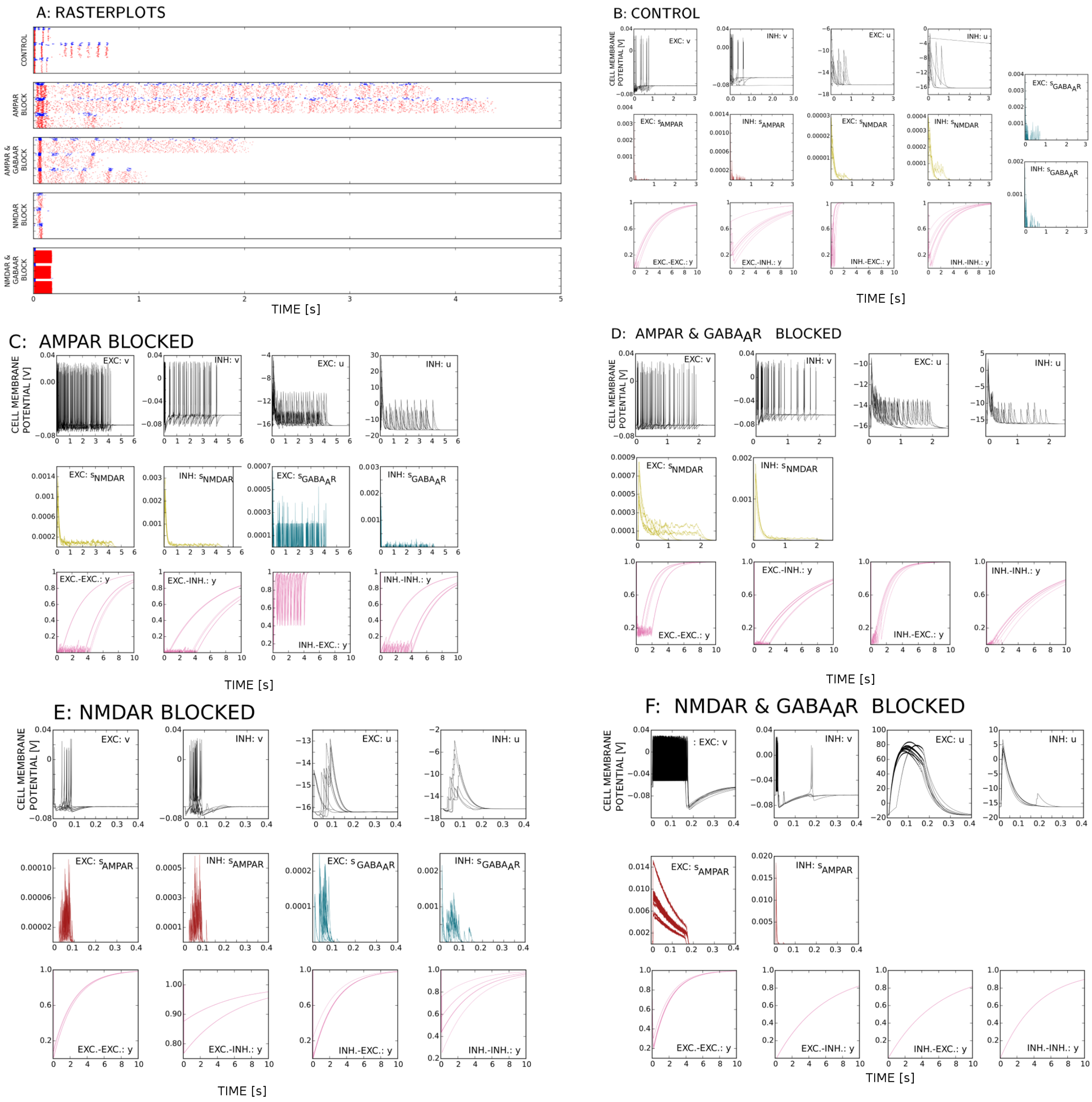


**Fig S3-11** Time course of model variables in a model with variability in neuronal parameters. **A:** Rasterplots from Fig 4 are shown again for completeness. **B:** Control condition. **C** AMPAR blocked condition. **D:** AMPAR and GABA$_A$R blocked condition. **E:** NMDAR blocked condition. **F:** NMDAR and GABA$_A$R blocked condition The panels show variables v and u (Izhkevich model) for the random three excitatory and three inhibitory neurons (top row); AMPAR, NMDAR and GABA$_A$R conductances for the random three synapses (middle row), and the variable representing the availability of presynaptic resources for the random three synapses (bottom row). The variables show variation in dynamics of individual neurons and synapses. The variability in neuronal dynamics might be caused by input difference, but also from randomized model parameters. The variation in synaptic dynamics results solely from difference in inputs as all synapses of the same type are identical.
