## Supplementary data 4 for "Analysis of cellular and synaptic mechanisms behind spontaneous cortical activity *in vitro*: Insights from optimization of spiking neuronal network models"

### S4: Naïve approach in fitting one model to all pharmacological conditions

In addition to the tests shown in the paper, we examined to what extent the trends observed in the experimental data can be reproduced using the minimally complex model fitting routine and the minimal training data set. To do this, we optimized a model to the training set extracted from the control data, and then performed *in silico* blocking of synaptic receptors equivalent, in the context of this model, to the experiments with *in vitro* receptor blocking. We tested this approach using ten models from Section 2 optimized to control condition and obtained starting from ten different random number generator seeds. Blocking of all synaptic conductances of a certain type by setting the peak conductance to zero resulted in unrealistic short distributions of burst lengths and small normalized burst sizes. An exception was fitting of small burst sizes of NMDAR blocked condition which appeared larger in simulated data. However, this simple blocking dramatically changed the level of synaptic inputs to each neuron which inevitably altered the network level activity. Thus, we compensated the removed synaptic inputs by tuning the peak conductances of the remaining active receptor types according to the following rules:

- In experiments involving glutamatergic and gabaergic blocking, i.e. NMDAR & $\mathrm{GABA_AR}$ or AMPAR & $\mathrm{GABA_AR}$ blocking, for each synapse $j, i = 1..N$ the peak conductance of the remaining glutamatergic receptor (glu = AMPAR or NMDAR) was scaled to account for the lost synaptic currents according to the following equation. Here, $g^*_{glu,ji}$ is the peak conductance after blocking while $g_{glu,ji}$ is the peak conductance in control condition. For details about notation see model description in Appendix 1.

$$g^*_{glu,ji} = g_{AMPAR,ji} + g_{NMDAR,ji} - g_{GABA_AR,ji} \cdot \frac{N - N_E}{N_E}$$

- For *in silico* blocking of a single glutamatergic receptor, AMPAR or NMDAR blocking, the peak conductance of the remaining glutamatergic receptor was set to the sum of peak AMPAR and NMDAR conductance values from control condition.

$$g^*_{glu,ji} = g_{AMPAR,ji} + g_{NMDAR,ji}$$

Figure 4 summarizes the results. The top row in Figure 4 shows the distributions of burst lengths in five considered experiments while the bottom row contains distributions of burst sizes. Grey lines correspond to individual models, red lines are the distributions obtained by pooling data obtained using all ten models, while blue lines are the distributions obtained from experimental data. The distances between experimental and simulated data are given in the upper right corner of each panel for each of the ten models. The average distances for control condition (the condition used to optimized models) are 0.191 for burst length and 0.151 for burst sizes, while the average distances for other conditions (obtained after in silico blocking) are 0.76 and 0.845 (burst length and size for NMDAR and $\mathrm{GABA_AR}$ blocking), 0.73 and 0.76 (BL and BS for AMPAR and $\mathrm{GABA_AR}$ blocking), 0.7 and 0.95 (BL and BS for NMDAR blocking), 0.73 and 0.79 (BL and BS for AMPAR blocking). Although some models achieved realistic values of burst measures for some conditions, no model managed to approach the experimental data for more than one experimental condition other than control. These tests suggest that sole *in silico* blocking of synaptic receptors cannot reproduce experimental findings as such. Accurate results require either the use of large data sets that contain enough information on all conditions of interest, or a very carefully constructed model manually tuned to reproduce all conditions and data.

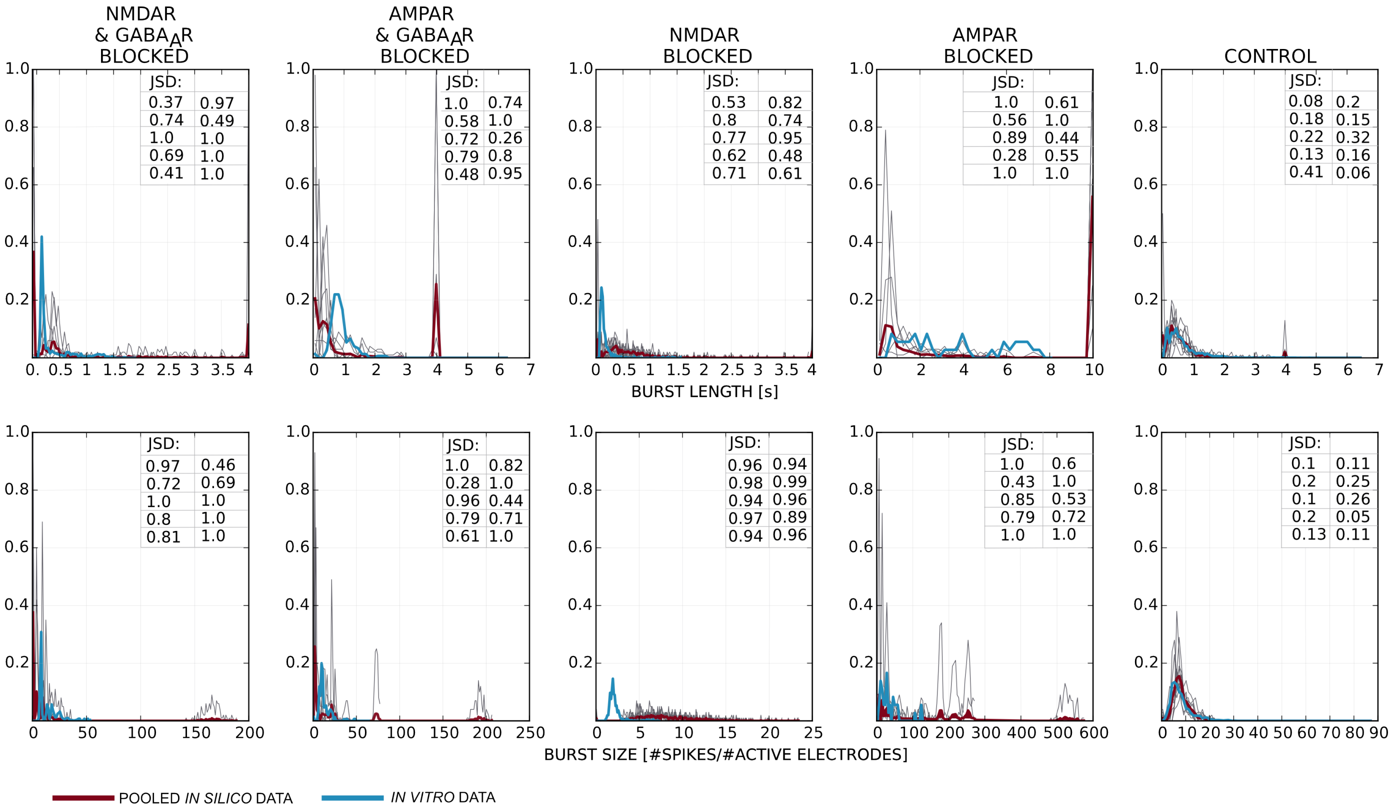


The figure summarizes attempt to reproduce all experimental conditions by fitting a single model to control condition, and then performing *in silico* blocking of appropriate synaptic receptors. Upper row - distribution of burst lengths, bottom row - distribution of burst sizes. Each column corresponds to one experimental condition indicated at the top of the column. Grey lines - the results of 10 models optimized to the control data, red - all 10 models pooled together, blue - experimental *in silico* data. The similarity between the experimental and *in silico* expressed using JSD (JSD=0 for identical histograms, JSD=1 for maximally divergent histograms) is presented in the upper right corner of each panel.
