## Supplementary data 5, constrained protocol for "Analysis of cellular and synaptic mechanisms behind spontaneous cortical activity *in vitro*: Insights from optimization of spiking neuronal network models"

### Supplementary material S5

September 29, 2021

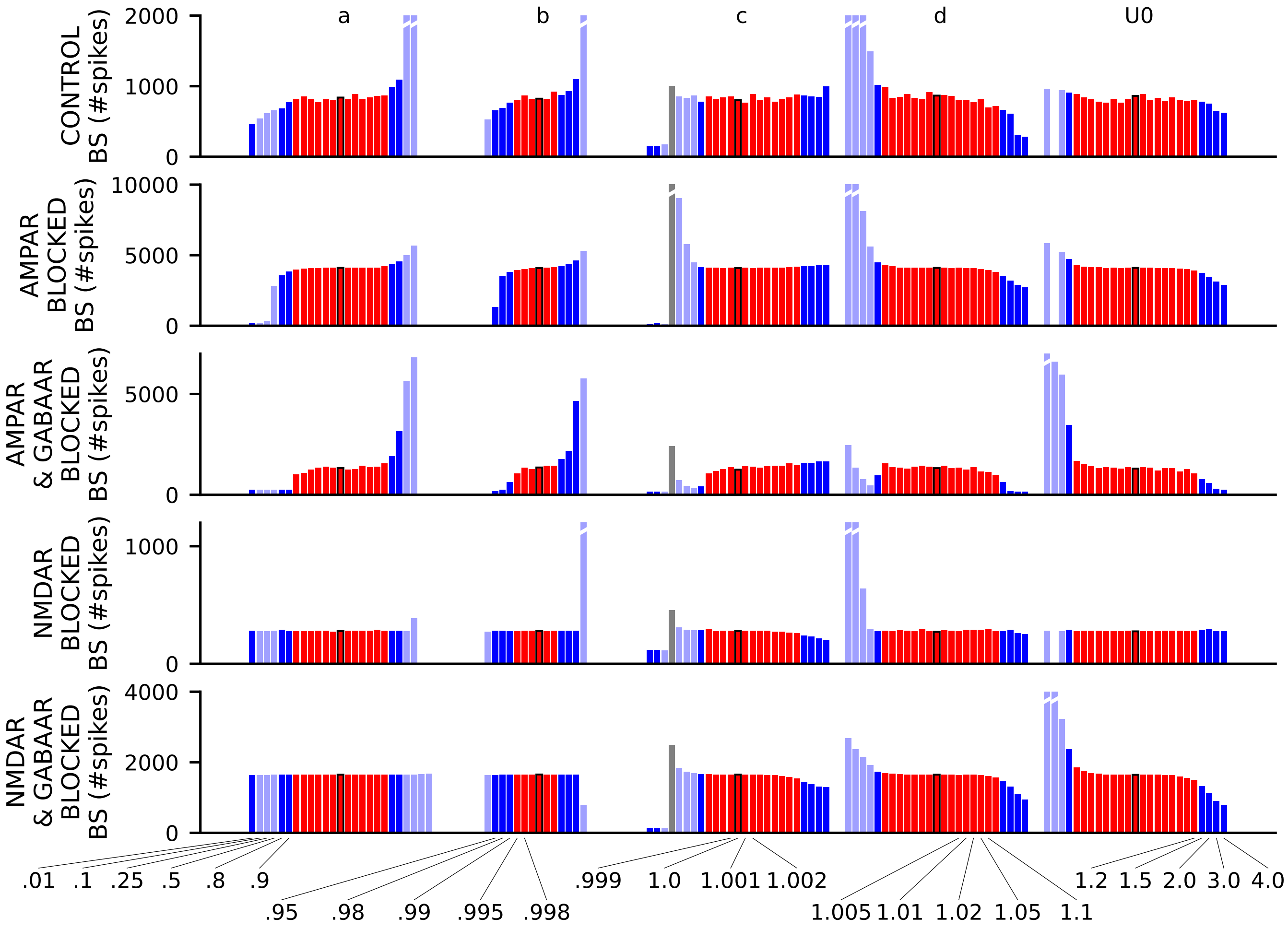


Figure S5-1: Burst size sensitivity analysis of the network model obtained with the multi-objective optimization task where all mechanisms were included. The five panels show the predicted burst sizes (in number of spikes) in the five different pharmacological conditions. See Figure 7M for the corresponding results on burst lengths.

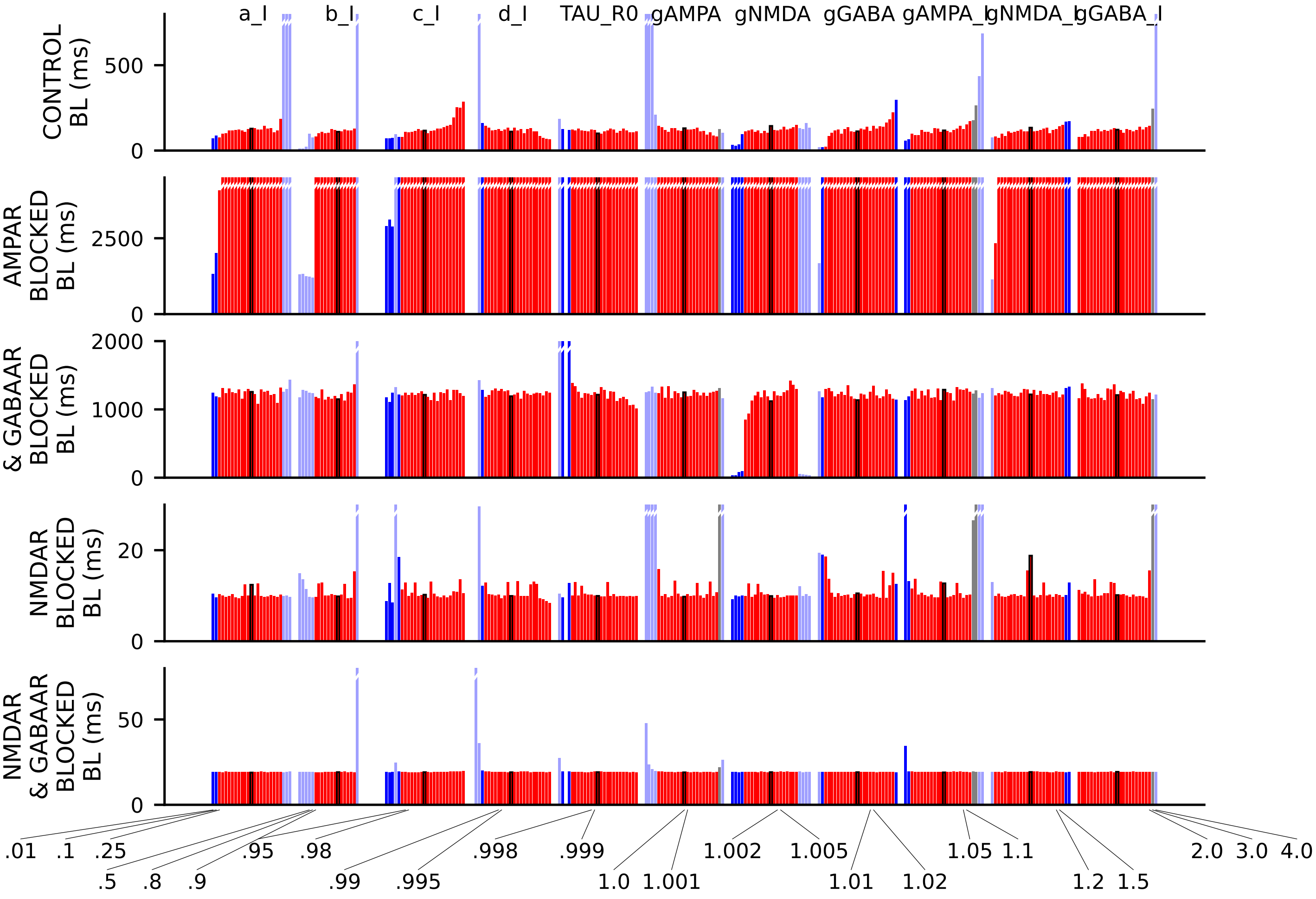


Figure S5-2: Burst length sensitivity analysis of the network model obtained with the multi-objective optimization task where all mechanisms were included. The five panels show the predicted burst lengths (in msec) in the five different pharmacological conditions. One of the eleven parameters were multiplied with a coefficient ranging from 0.01 to 1.0 (unaltered, i.e., from the optimization; highlighted with black edges) and all the way to 4.0. The exact coefficients are listed at the bottom of the figure. The eleven varied parameters were the Izhikevich model parameters of the inhibitory neuron $a_I$ (left), $b_I$, $c_I$, and $d_I$, the time constant of recovery from synaptic depression $\tau_{R0}$, the synaptic conductances to the excitatory neurons ($g_{AMPA}$, $g_{NMDA}$, $g_{GABA}$), and the synaptic conductances to the inhibitory neurons ($g_{AMPA,I}$, $g_{NMDA,I}$, $g_{GABA,I}$) (right), See Figure 7M for the corresponding data on the sensitivity with respect to the other five model parameters.

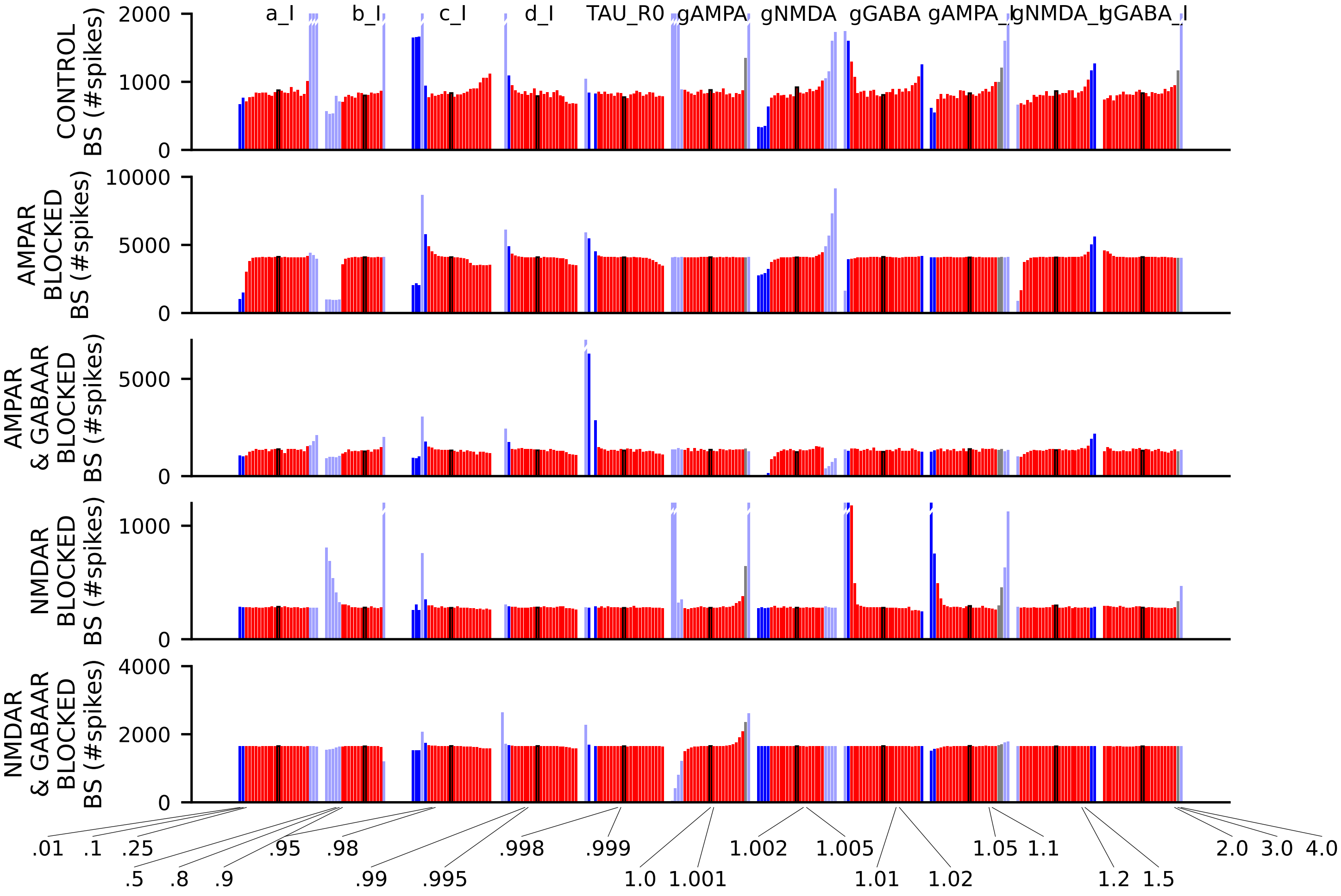


Figure S5-3: Burst size sensitivity analysis of the network model obtained with the multi-objective optimization task where all mechanisms were included. The five panels show the predicted burst sizes (in number of spikes) in the five different pharmacological conditions. The eleven varied parameters were as in Figure S5-2. See Figure S5-2 for the corresponding data on burst lengths.

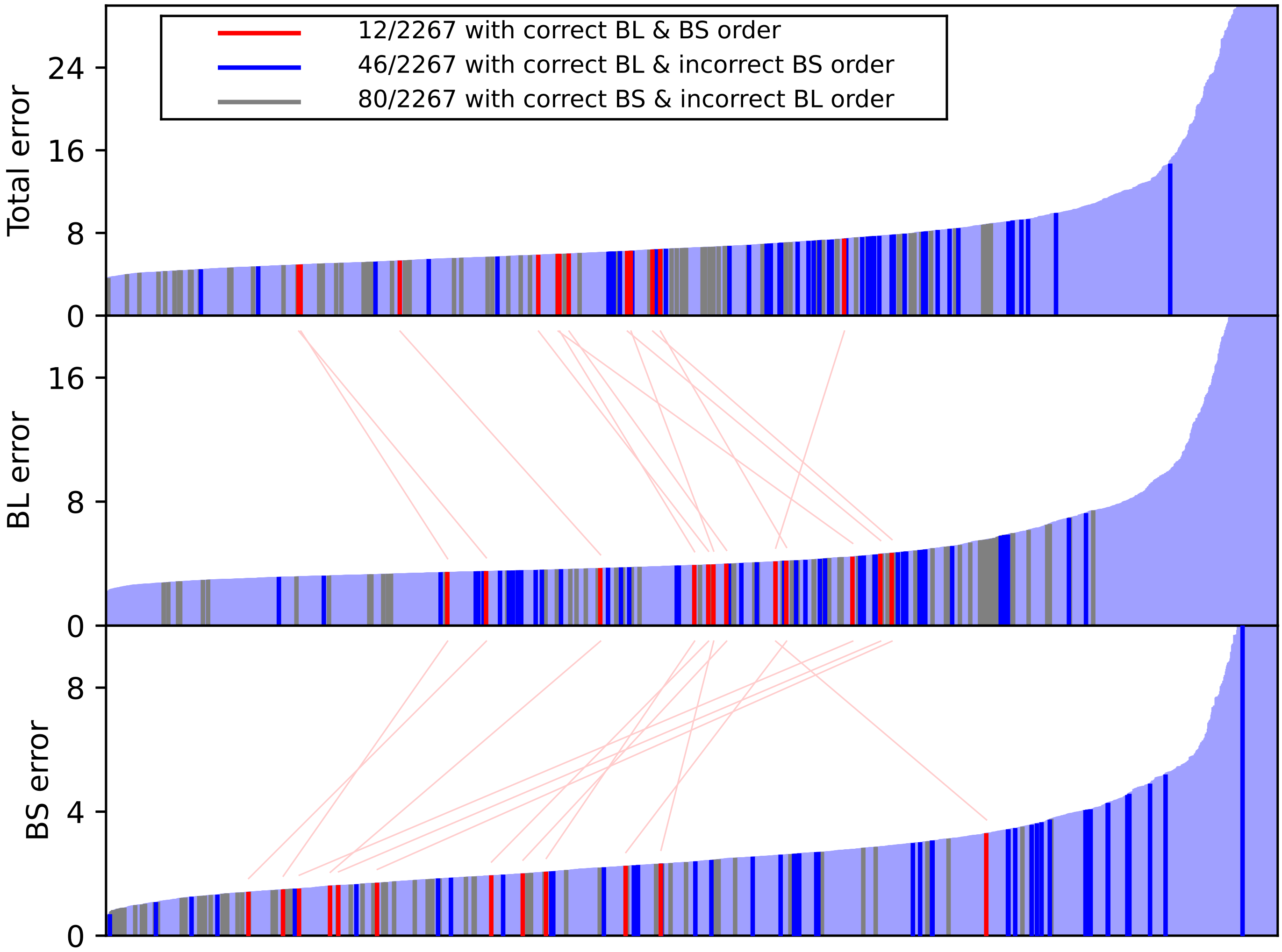


Figure S5-4: Fitness error values from final parameter sets from 10 repetitions of the model fitting when synaptic depression was excluded ($U0$ and $\tau_{R0}$ not fitted, all synapses were static). A total of 2267 unique parameter sets were obtained from the multi-objective optimization. See Figure 6B for the corresponding data from optimization where synaptic depression was included.

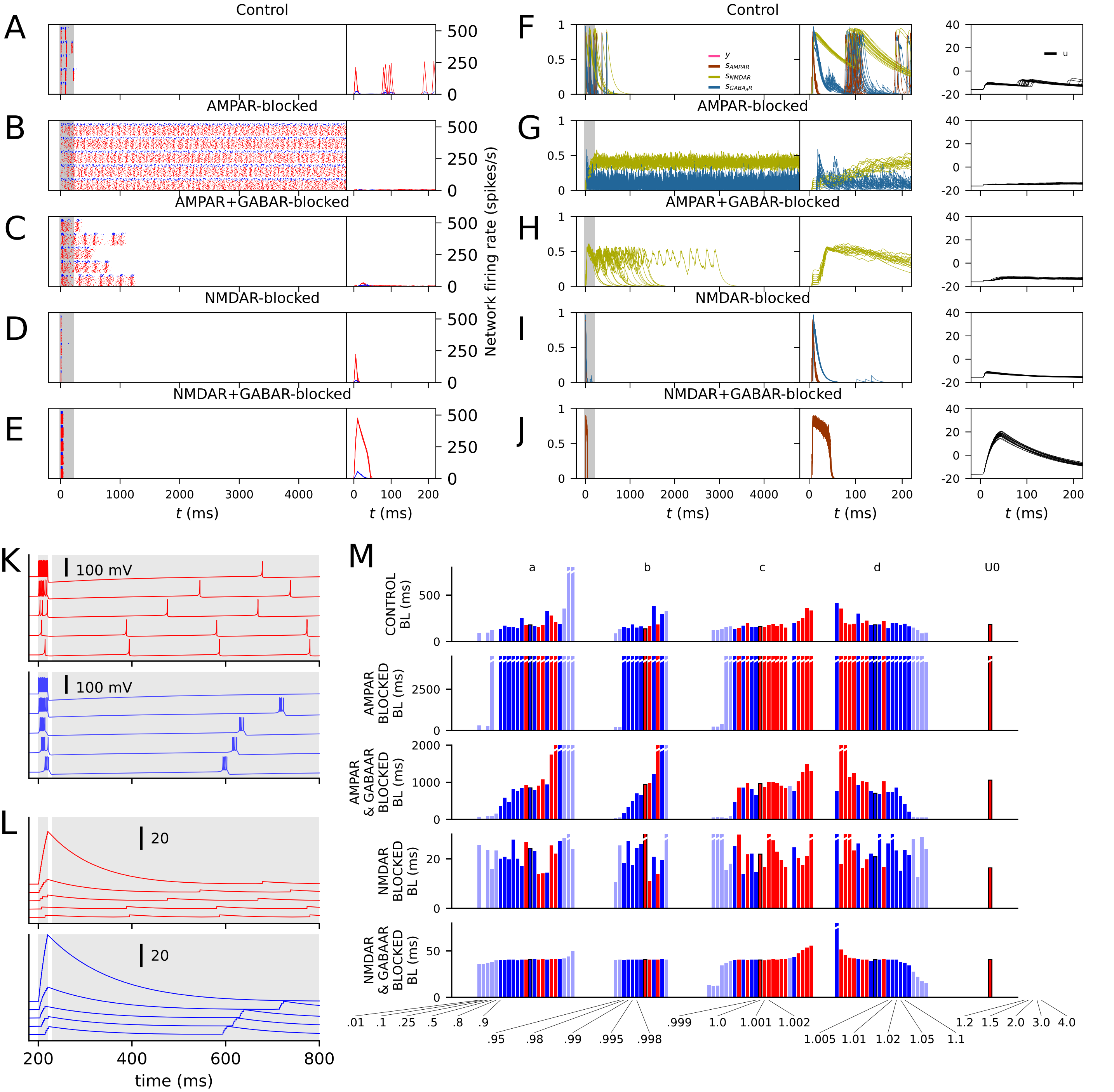


Figure S5-5: Illustration of the network model obtained from the multi-objective optimization where synaptic depression was excluded. See Figure 7 for the corresponding data from optimization where synaptic depression was included.

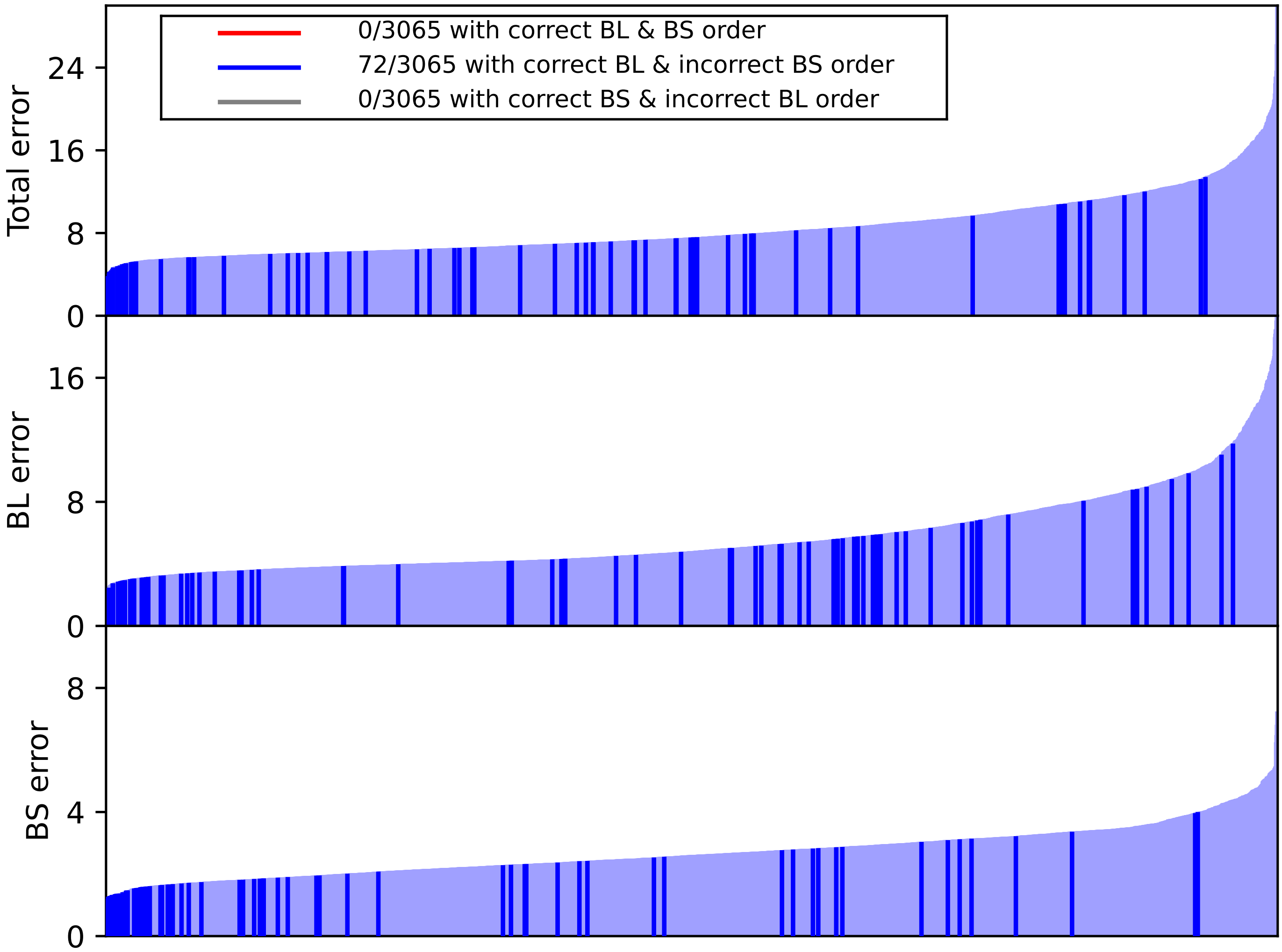


Figure S5-6: Fitness error values from final parameter sets from 10 repetitions of the model fitting when adaptation from both excitatory and inhibitory neurons was excluded (the tonic firing parameters $a$=0.02, $b$=0.254, $c$=-65, $d$=2 were used for all neurons). A total of 3065 unique parameter sets were obtained from the multi-objective optimization. See Figure 6B for the corresponding data from optimization where the Izhikevich parameters were fitted along the rest of the model parameters.

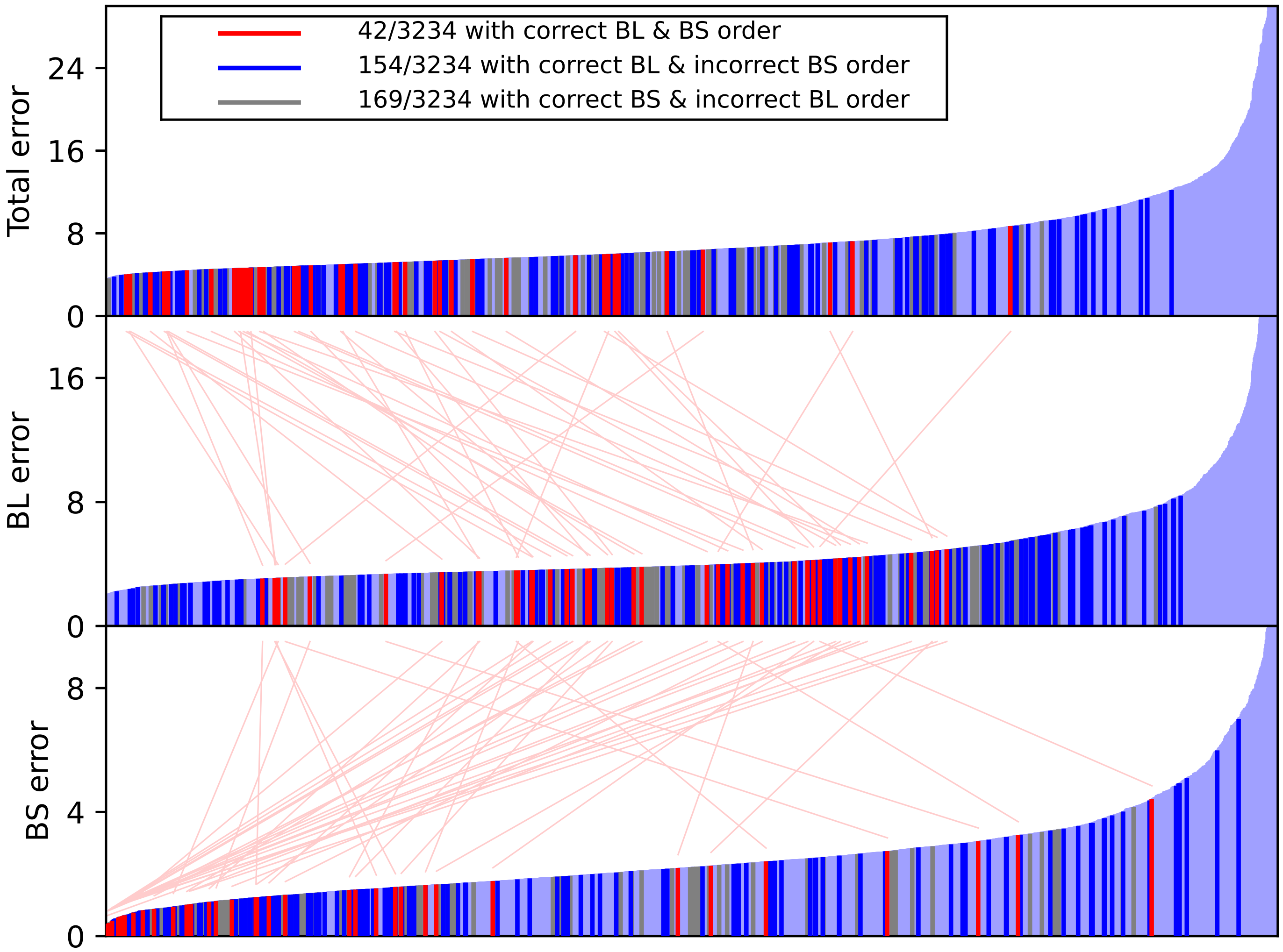


Figure S5-7: Fitness error values from final parameter sets from 10 repetitions of the model fitting when variability in Izhikevich parameters was excluded (all excitatory neurons had equal Izhikevich model parameters, and inhibitory neurons likewise). A total of 3234 unique parameter sets were obtained from the multi-objective optimization. See Figure 6B for the corresponding data from optimization where the variability in Izhikevich parameters was included.

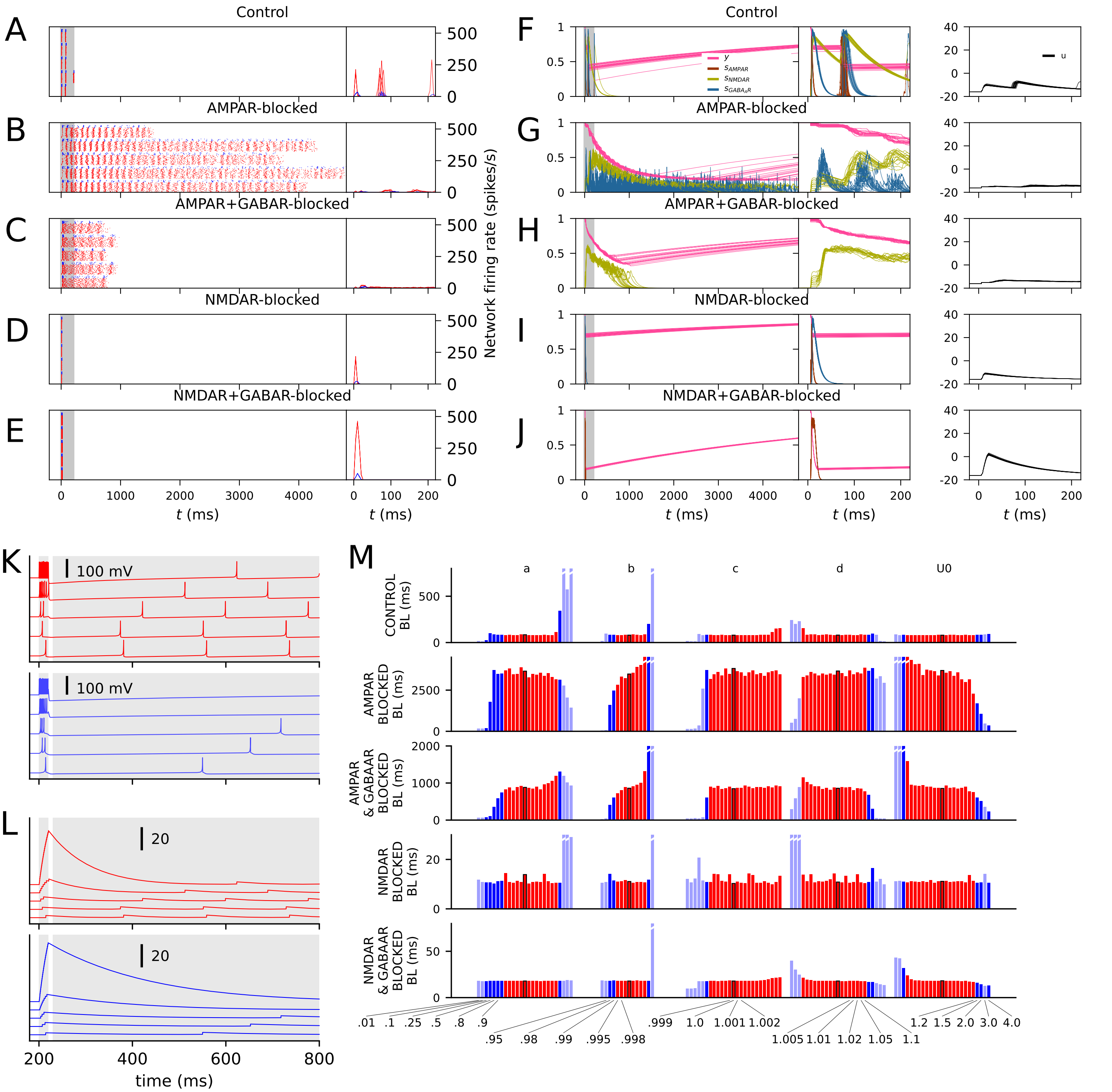


Figure S5-8: Illustration of the network model obtained from the multi-objective optimization where variability in Izhikevich parameters was excluded. See Figure 7 for the corresponding data from optimization where the variability in Izhikevich parameters was included.

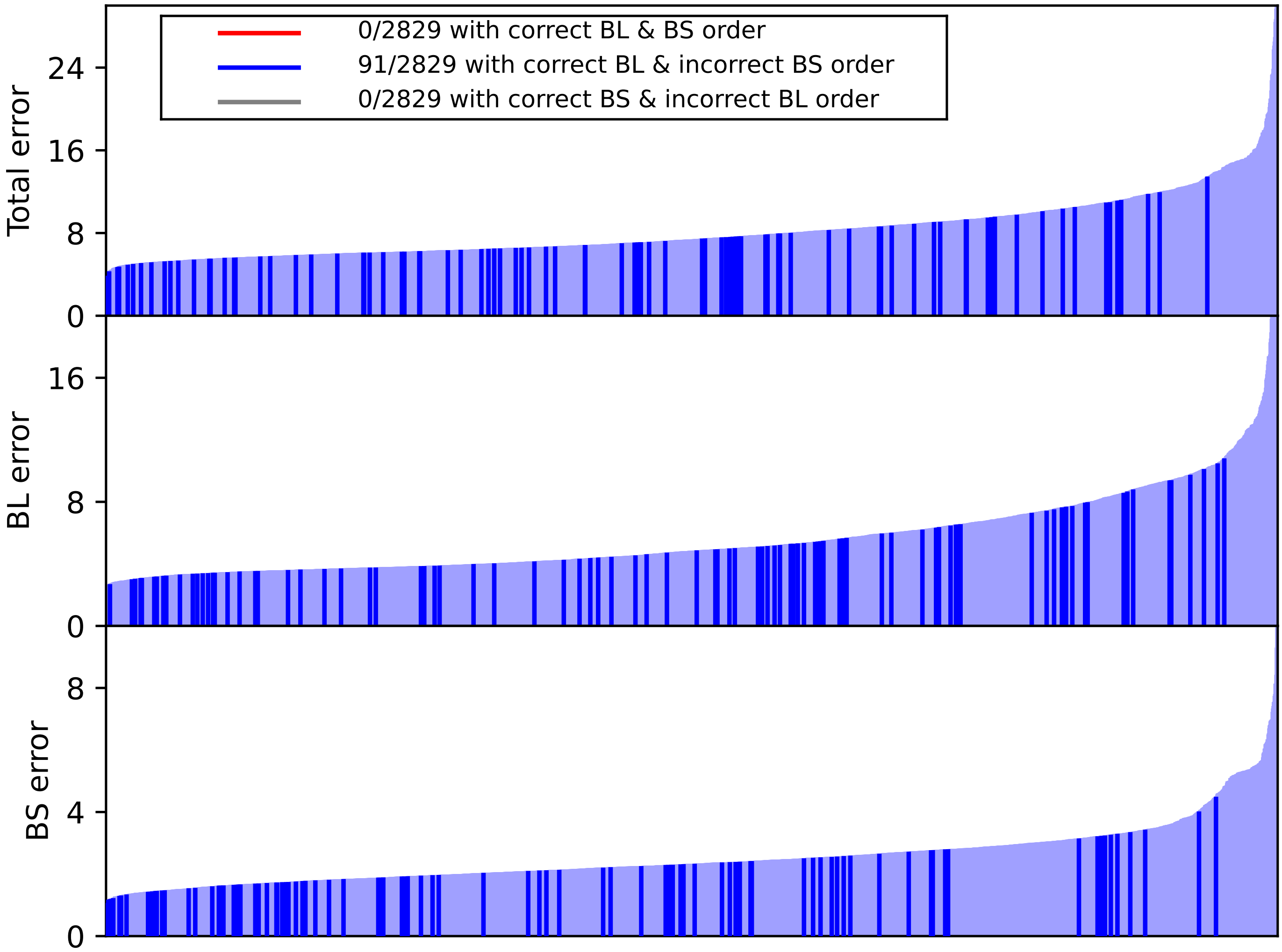


Figure S5-9: Fitness error values from final parameter sets from 10 repetitions of the model fitting when adaptation from both excitatory and inhibitory neurons was excluded (the tonic firing parameters $a$=0.02, $b$=0.254, $c$=-65, $d$=2 were used for all neurons), but the maximum allowed values for synaptic conductances were 150% larger than in other fittings. A total of 2829 unique parameter sets were obtained from the multi-objective optimization. See Figure 6B for the corresponding data from optimization where synaptic depression was included.

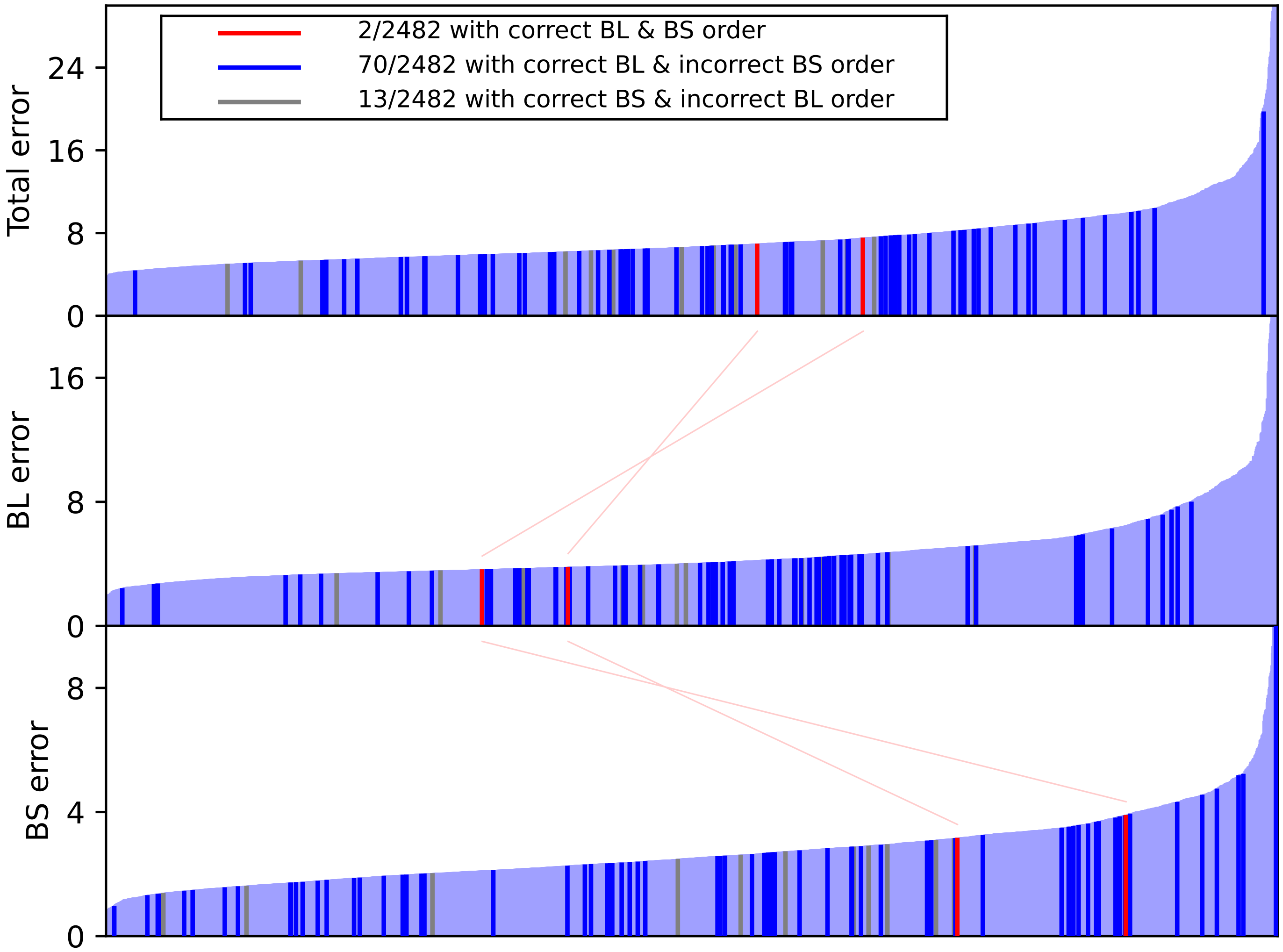


Figure S5-10: Fitness error values from final parameter sets from 10 repetitions of the model fitting when adaptation from inhibitory neurons was excluded (the tonic firing parameters $a$=0.02, $b$=0.254, $c$=-65, $d$=2 were used for inhibitory neurons). A total of 2482 unique parameter sets were obtained from the multi-objective optimization. See Figure 6B for the corresponding data from optimization where the Izhikevich parameters of the inhibitory neurons were fitted along the rest of the model parameters.

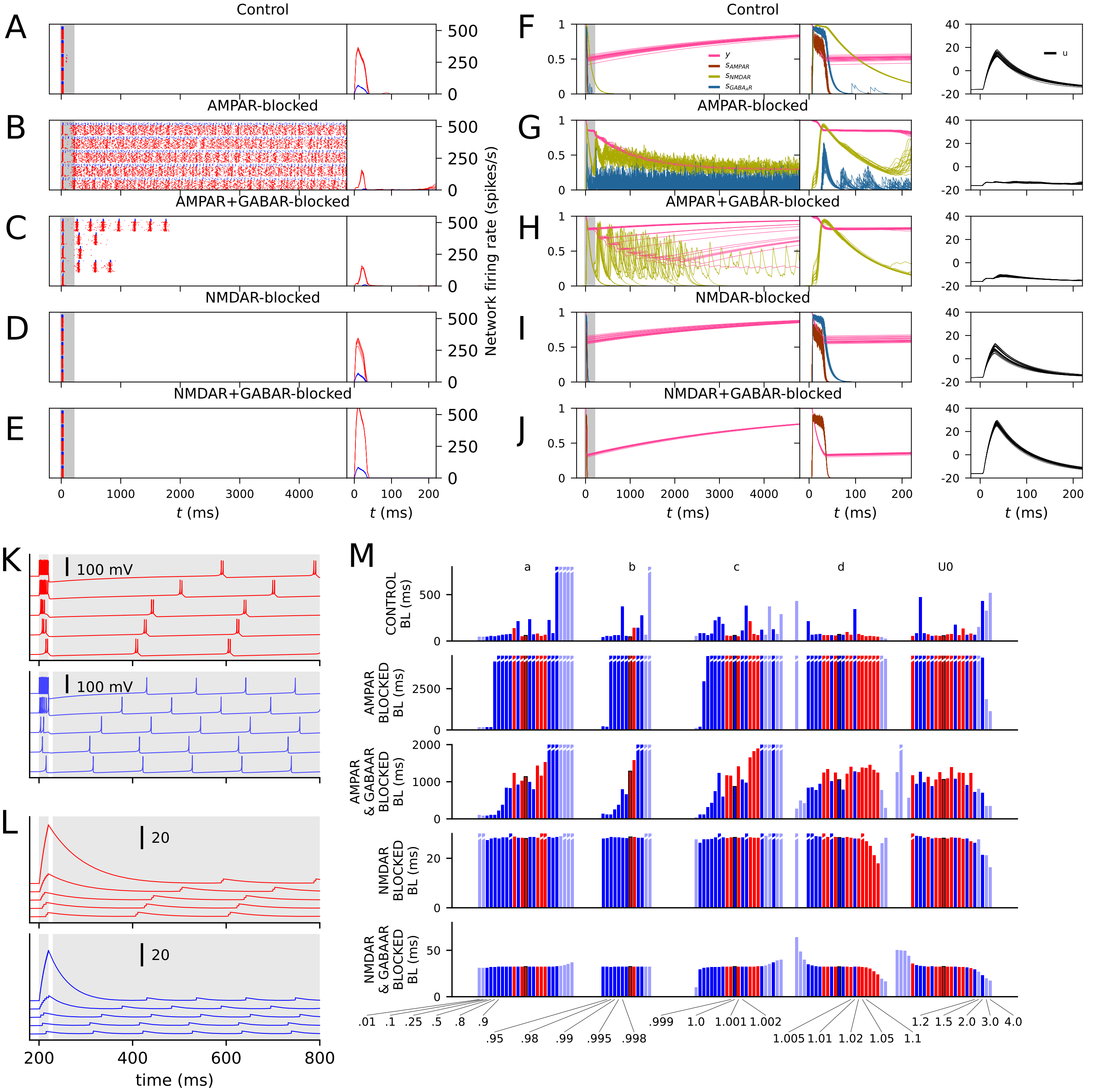


Figure S5-11: Illustration of the network model obtained from the multi-objective optimization where adaptation from inhibitory neurons was excluded (the tonic firing parameters $a$=0.02, $b$=0.254, $c$=-65, $d$=2 were used for inhibitory neurons). See Figure 7 for the corresponding data from optimization where the Izhikevich parameters of the inhibitory neurons were fitted along the rest of the model parameters.

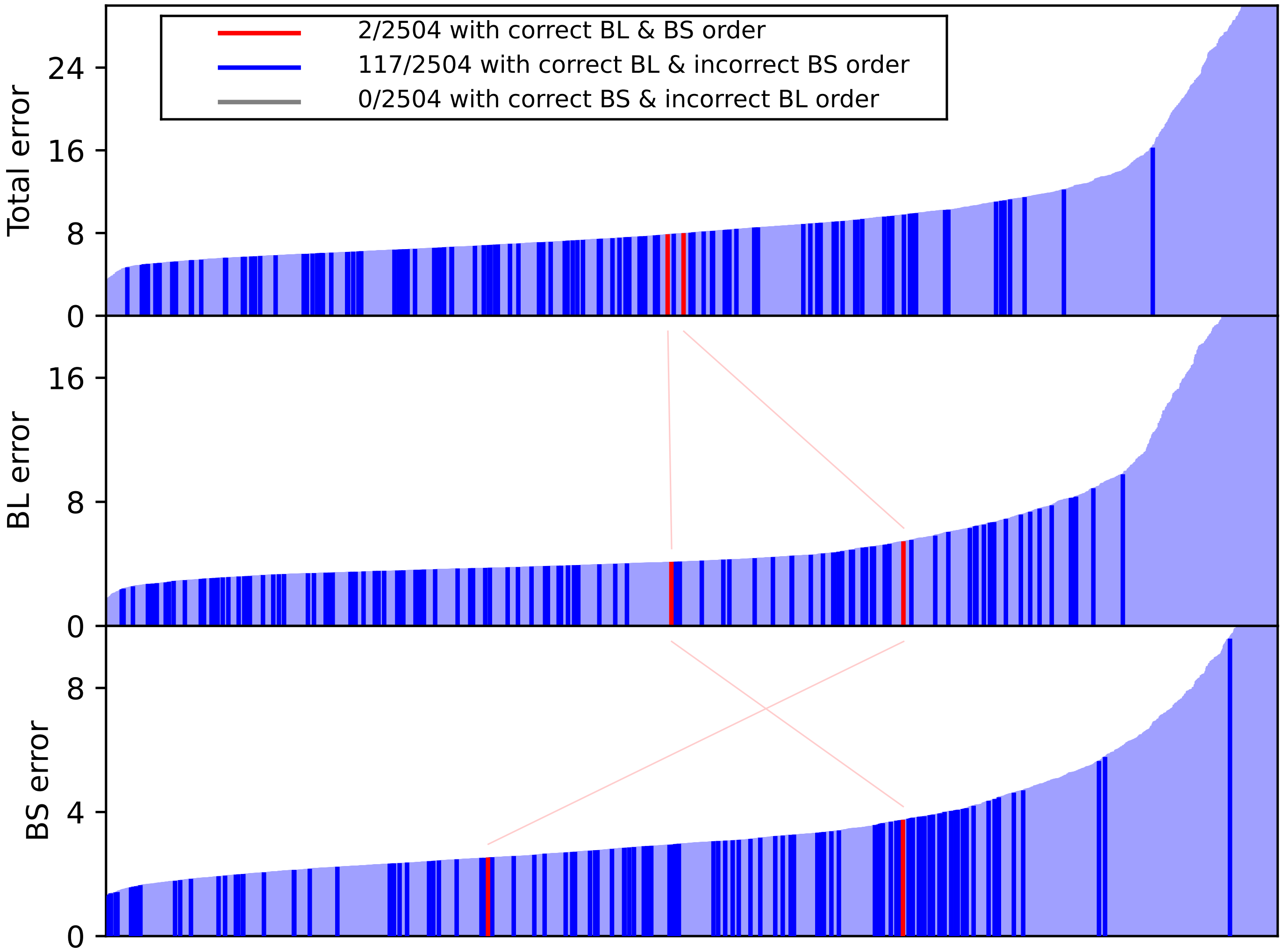


Figure S5-12: Fitness error values from final parameter sets from 10 repetitions of the model fitting when adaptation from excitatory neurons was excluded (the tonic firing parameters $a$=0.02, $b$=0.254, $c$=-65, $d$=2 were used for excitatory neurons). A total of 2504 unique parameter sets were obtained from the multi-objective optimization. See Figure 6B for the corresponding data from optimization where the Izhikevich parameters of the excitatory neurons were fitted along the rest of the model parameters.

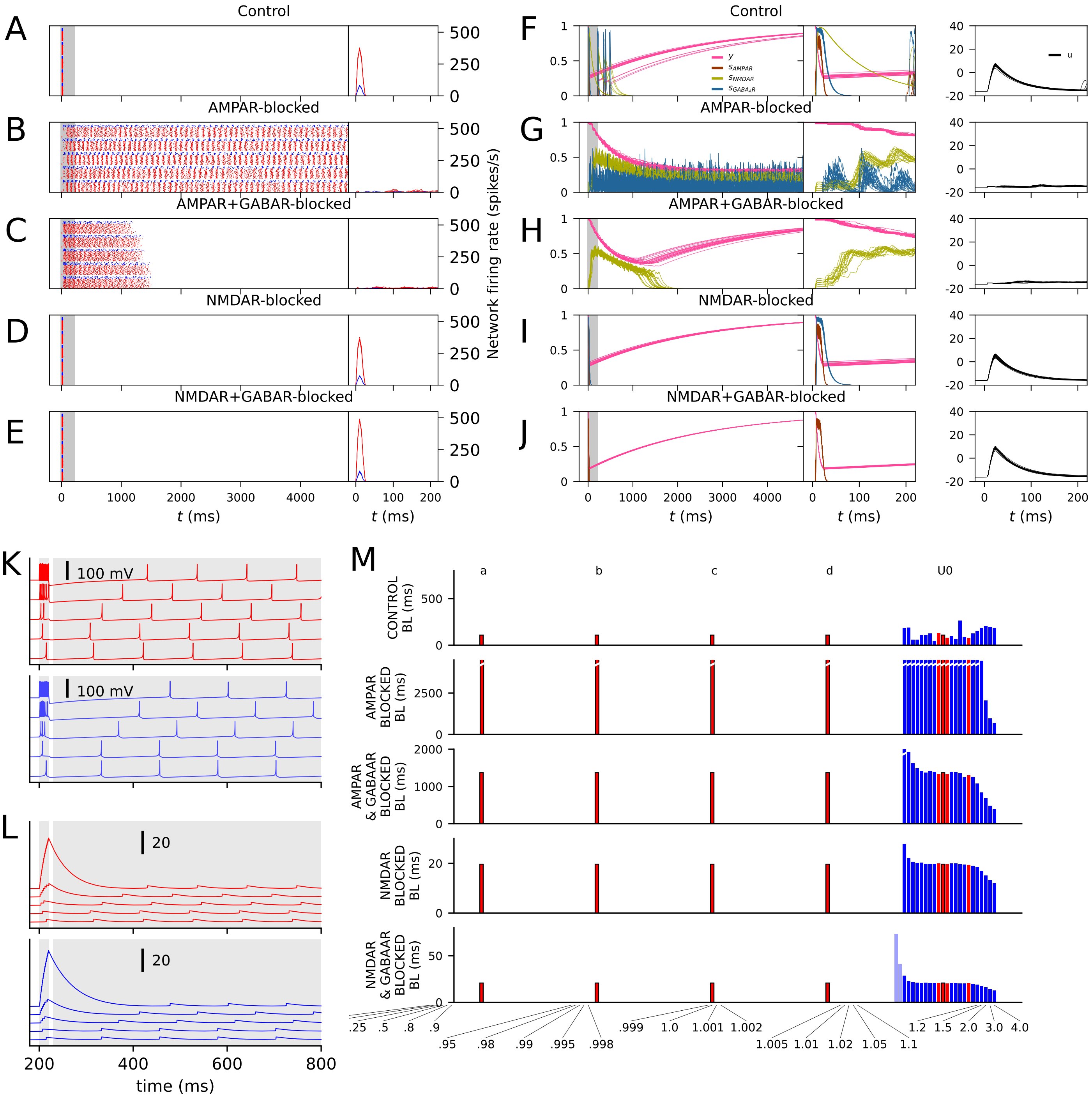


Figure S5-13: Illustration of the network model obtained from the multi-objective optimization where adaptation from excitatory neurons was excluded (the tonic firing parameters $a$=0.02, $b$=0.254, $c$=-65, $d$=2 were used for excitatory neurons). See Figure 7 for the corresponding data from optimization where the Izhikevich parameters of the excitatory neurons were fitted along the rest of the model parameters.

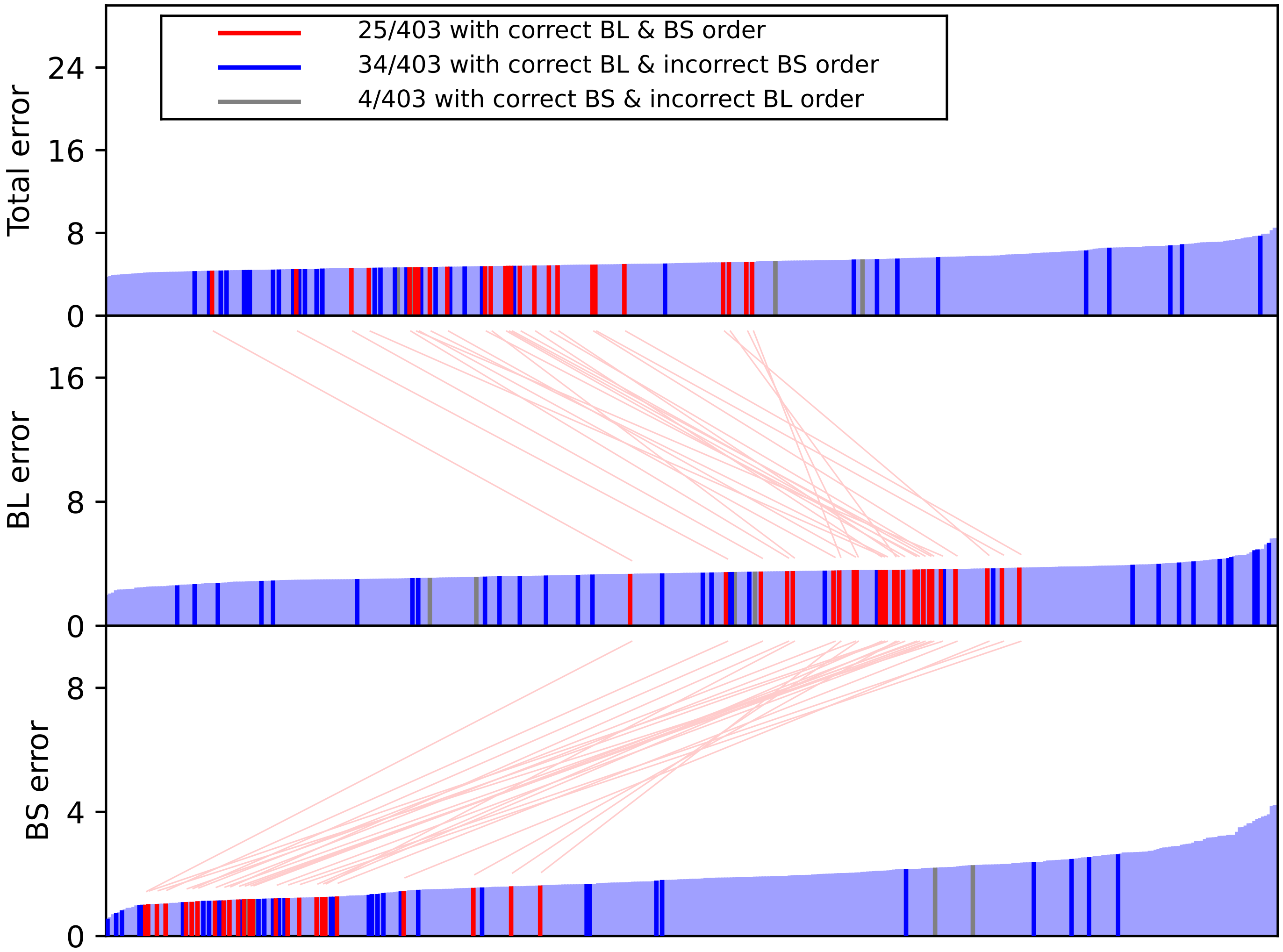


Figure S5-14: Fitness error values from final parameter sets from 5 repetitions of fitting of the model with NSGA-III. A total of 403 unique parameter sets were obtained from the multi-objective optimization. See Figure 6B for the corresponding data from optimization with NSGA-II.

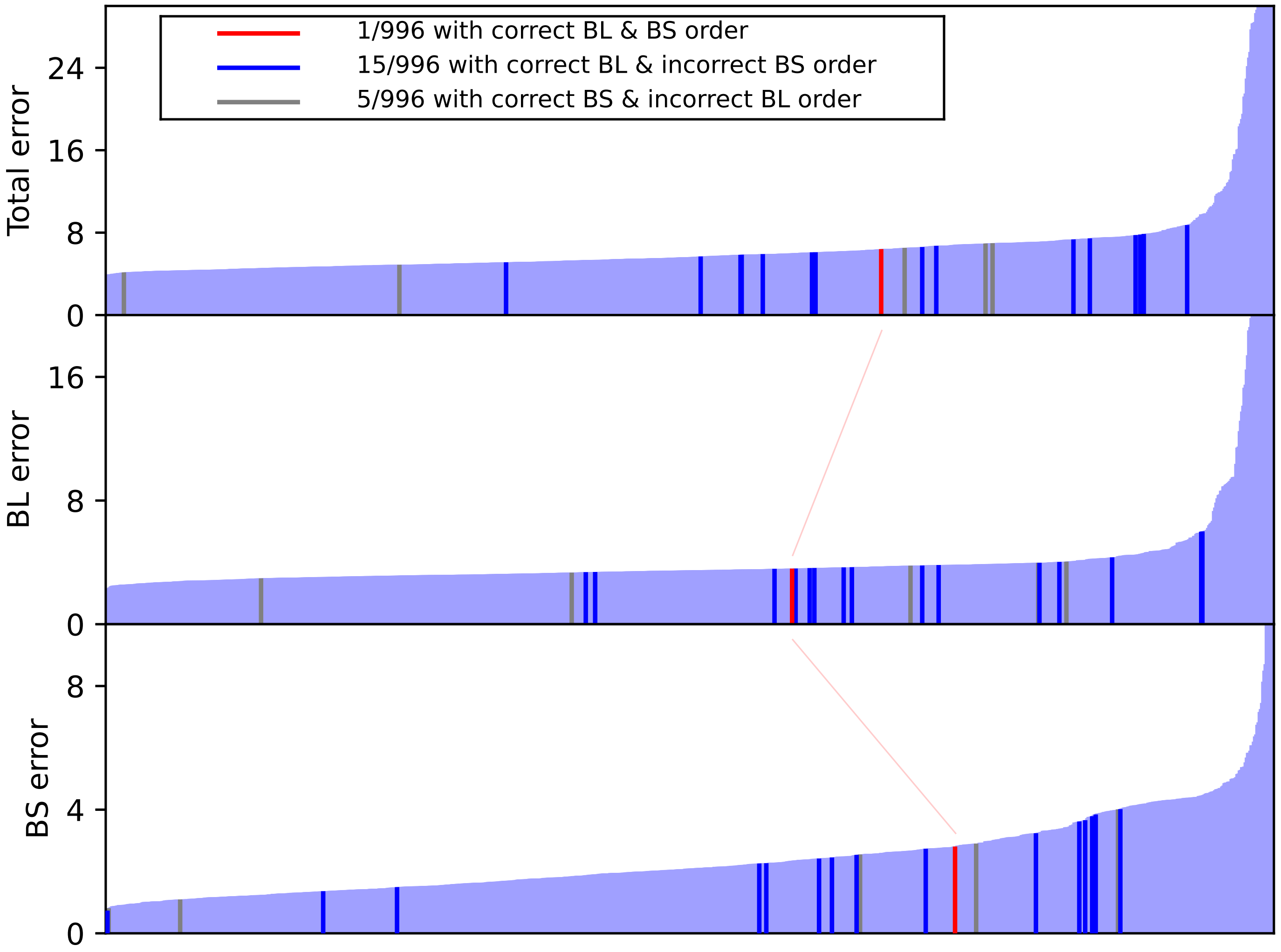


Figure S5-15: Fitness error values from final parameter sets from 5 repetitions of fitting of the model with NSGA-III when synaptic depression was excluded ($U0$ and $\tau_{R0}$ not fitted, all synapses were static). A total of 996 unique parameter sets were obtained from the multi-objective optimization. See Figure S5-14 for the corresponding data from optimization where synaptic depression was included.

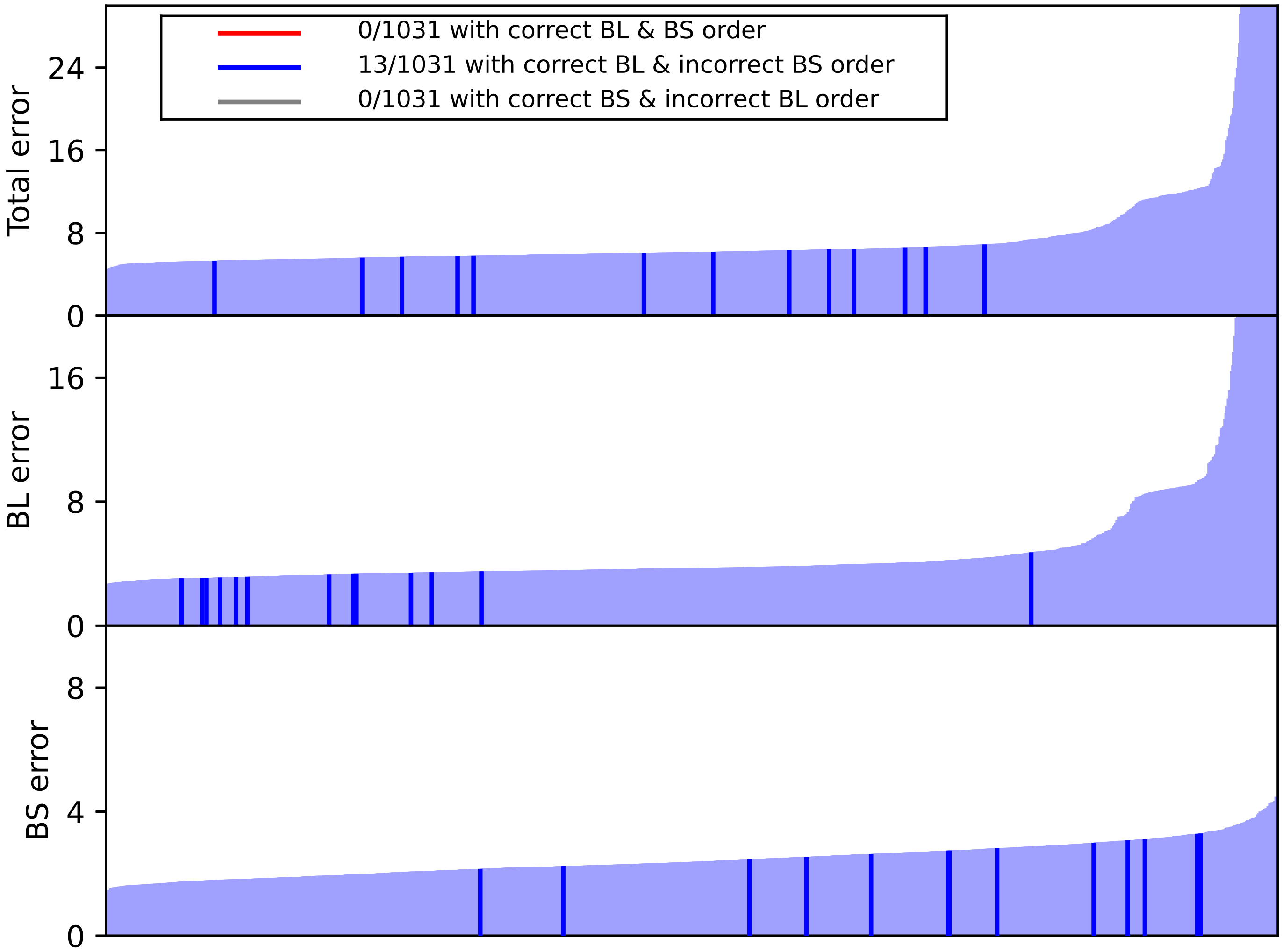


Figure S5-16: Fitness error values from final parameter sets from 5 repetitions of fitting of the model with NSGA-III when adaptation from both excitatory and inhibitory neurons was excluded. A total of 1031 unique parameter sets were obtained from the multi-objective optimization. See Figure S5-14 for the corresponding data from optimization where adaptation was included.

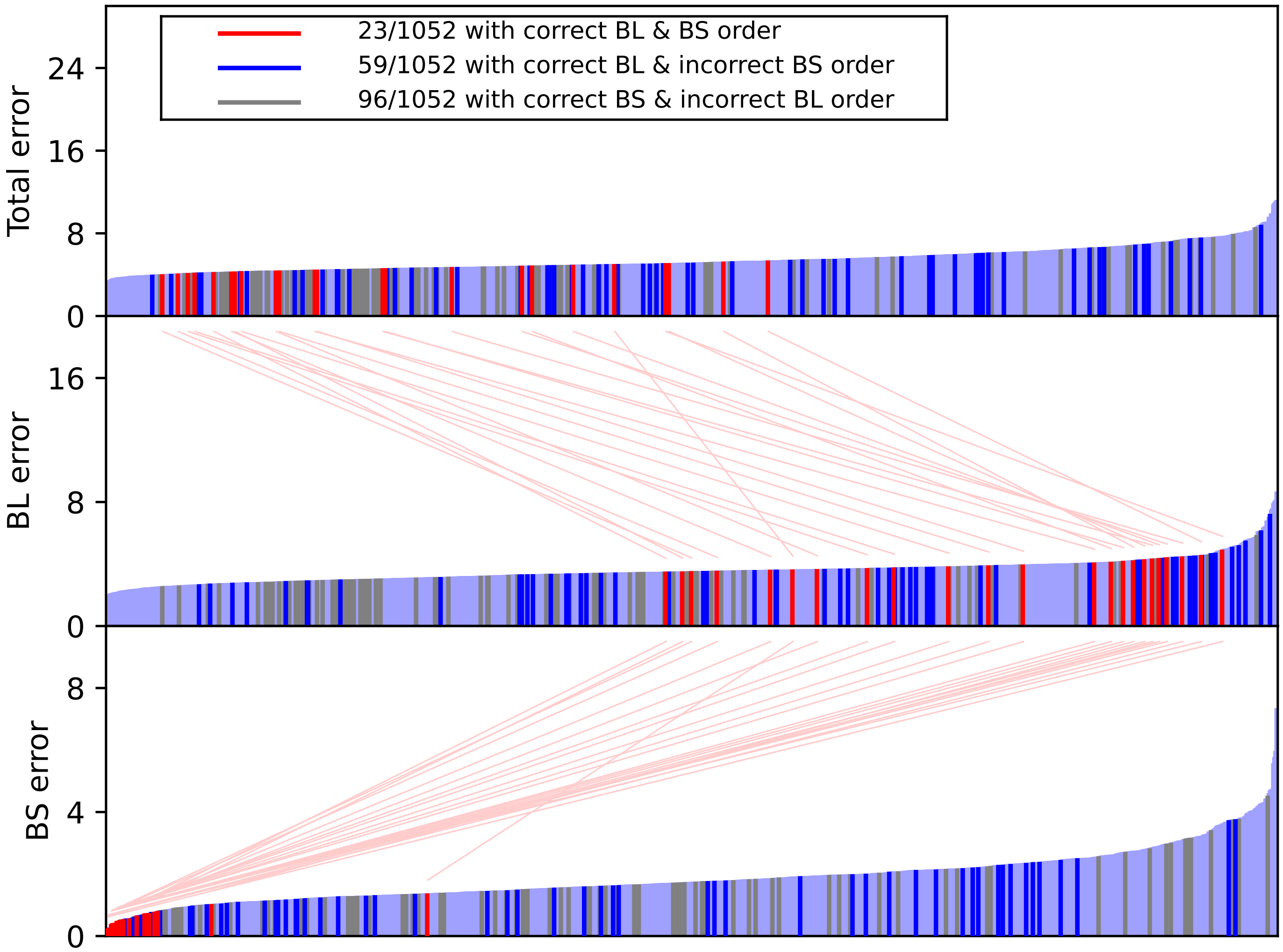


Figure S5-17: Fitness error values from final parameter sets from 5 repetitions of fitting of the model with NSGA-III when variability in Izhikevich parameters was excluded. A total of 1052 unique parameter sets were obtained from the multi-objective optimization. See Figure S5-14 for the corresponding data from optimization where variability in Izhikevich parameters was included.
