## Supplementary document 6, description of data and code for "Analysis of cellular and synaptic mechanisms behind spontaneous cortical activity *in vitro*: Insights from optimization of spiking neuronal network models"

S6: Python code, experimental and simulated data

Here, we summarize the material used to produce the results presented in the paper and the appendices. The Python scripts used to present the experimental data, implement the computational model, train and test the model, and prepare the result figures are available in modelDB. The repository is currently private and the accession code will be shared with reviewers. Upon acceptance, the code will be made public.

The experimental data set needed to reproduce the results can be found in:

flexible_protocol/Experimental_BL_BS_histograms.p (saved in the format compatible with Python 2.7)

flexible_protocol/Experimental_BL_BS_histograms_for_Python3.py (the format compatible with Python 3.3)

The code used to implement the flexible protocol, analyze the results and prepare figures (Fig 3, 4, 5) shown in the paper and the Appendices 3 and 4 is saved in the folder 'flexible_protocol'. All code is written in Python. Implementation and simulation of spiking neuronal network models is done in Brian2 simulator.

The code used to implement the constrained protocol, analyze the results and prepare figures (Fig 6,7,8) and the Appendix 5 is saved in the folder 'constrained_protocol'. The code is written in Python and implementation of the spiking neuronal network model is done in Neuron simulator, version 7.5-7.7.

The genetic algorithm, used for optimization, is a publicly available Python package presented in Bahl et al (2012) and also used in Mäki-Marttunen et al (2017). The package can be downloaded from: https://projects.g-node.org/emoo/

The following Python scripts are used to plot individual result figures:

| Figure | modelDB folder | Code |
|---|---|---|
| Figure 2 | flexible_protocol | Fig2.py |
| Figure3 | flexible_protocol | convergence_best_model_Fig3_Fig4.py |
| Figure 4<br>Fig 4A-H<br>Fig 4I-M<br>Fig 4N, Fig A3-5<br>Fig 4O | flexible_protocol | <br>convergence_best_model_Fig3_Fig4.py<br>Fig4.py<br>Fig4_plot_sensitivity.py<br>Fig4O_sensitivity_summary.py |
| Figure 5<br>Fig 5A-J<br>Fig 5K,L | flexible_protocol | <br>Fig5.py<br>Fig5_KL.py |
| Figure 6<br>Fig 6A<br>Fig 6B | constrained_protocol | <br>drawfig0a.py<br>drawfig0b.py |

| | | |
|---|---|---|
| Figure 7<br>Fig 7A-E<br>Fig 7F-J<br>Fig 7K-L<br>Fig 7M | constrained_protocol | <br>drawfig1.py<br>drawfig2.py<br>drawfig3.py<br>drawfig4.py |
| Figure 8 | constrained_protocol | Drawfig5.py |
